## Supplementary_file_1 for "Single cell RNA sequencing unravels the transcriptional network underlying zebrafish retina regeneration"

**Cluster 1**

| | names | scores | logfoldchanges | pvals | pvals_adj |
|---|---|---|---|---|---|
| **0** | glula | 202.4346 | 5.833523273 | 0 | 0 |
| **1** | slc1a2b | 183.2414 | 5.140877247 | 0 | 0 |
| **2** | cahz | 182.9701 | 6.424218178 | 0 | 0 |
| **3** | selenop | 180.7526 | 5.330976486 | 0 | 0 |
| **4** | rlbp1a | 172.976 | 5.283491135 | 0 | 0 |
| **5** | aqp1a.1 | 170.692 | 6.171694279 | 0 | 0 |
| **6** | nfasca | 153.0779 | 4.269580364 | 0 | 0 |
| **7** | cdo1 | 152.254 | 5.08609581 | 0 | 0 |
| **8** | gstp1 | 151.2817 | 4.877549171 | 0 | 0 |
| **9** | fosab | 149.3541 | 5.28233242 | 0 | 0 |
| **10** | fxyd6l | 137.2993 | 3.659704685 | 0 | 0 |
| **11** | si:ch211-152c2.3 | 137.0068 | 6.221270084 | 0 | 0 |
| **12** | ptgdsb.2 | 136.8515 | 5.723503113 | 0 | 0 |
| **13** | jdp2b | 136.2845 | 5.292197227 | 0 | 0 |
| **14** | spock3 | 134.9536 | 4.483845234 | 0 | 0 |
| **15** | sparc | 132.7677 | 4.232047558 | 0 | 0 |
| **16** | fosb | 131.2225 | 4.369260311 | 0 | 0 |
| **17** | efhd1 | 130.836 | 4.415273666 | 0 | 0 |
| **18** | zfp36l1b | 130.4942 | 4.800184727 | 0 | 0 |
| **19** | eno1a | 128.9211 | 3.307034016 | 0 | 0 |
| **20** | icn | 127.1163 | 4.933670044 | 0 | 0 |
| **21** | fosl1a | 126.1979 | 4.024051666 | 0 | 0 |
| **22** | mt2 | 125.033 | 5.786454678 | 0 | 0 |
| **23** | ppp1r14aa | 123.0459 | 4.849192142 | 0 | 0 |
| **24** | s100a10b | 120.6196 | 4.405580044 | 0 | 0 |
| **25** | rhbg | 120.5902 | 4.047341824 | 0 | 0 |
| **26** | atp1a1b | 117.981 | 4.247701168 | 0 | 0 |
| **27** | zgc:165604 | 115.436 | 3.359636307 | 0 | 0 |
| **28** | pik3ip1 | 112.6725 | 4.668477535 | 0 | 0 |
| **29** | btg2 | 111.3905 | 3.155260086 | 0 | 0 |
| **30** | aplp2 | 108.6828 | 3.532805204 | 0 | 0 |
| **31** | mcl1b | 108.0181 | 3.970383883 | 0 | 0 |
| **32** | cavin2a | 105.313 | 4.867988586 | 0 | 0 |
| **33** | apoeb | 104.8825 | 4.027953148 | 0 | 0 |
| **34** | prdx6 | 103.3325 | 3.429146051 | 0 | 0 |
| **35** | jun | 101.6418 | 3.013286352 | 0 | 0 |
| **36** | socs3a | 101.591 | 4.512181282 | 0 | 0 |
| **37** | mcl1a | 101.506 | 3.701190948 | 0 | 0 |
| **38** | col15a1b | 100.4798 | 5.353167534 | 0 | 0 |
| **39** | si:dkey-16p21.8 | 99.58262 | 3.896369696 | 0 | 0 |
| **40** | mCherry | 96.94631 | 3.514686584 | 0 | 0 |
| **41** | slc38a4 | 95.86198 | 4.357030869 | 0 | 0 |
| **42** | junba | 95.83254 | 3.064756632 | 0 | 0 |
| **43** | insig1 | 95.09448 | 3.546940327 | 0 | 0 |
| **44** | ubb | 94.82858 | 3.611829519 | 0 | 0 |
| **45** | dap1b | 94.21604 | 3.266350746 | 0 | 0 |
| **46** | atp1b4 | 94.20795 | 6.214066982 | 0 | 0 |
| **47** | dab2ipa | 93.97268 | 3.303300381 | 0 | 0 |
| **48** | atp1b3a | 93.81084 | 3.831693649 | 0 | 0 |
| **49** | hsp70l | 93.31802 | 5.357698441 | 0 | 0 |
| **50** | epas1b | 92.86622 | 4.177621365 | 0 | 0 |
| **51** | hsp70.2 | 92.45098 | 5.219595909 | 0 | 0 |
| **52** | junbb | 92.05714 | 3.120440245 | 0 | 0 |
| **53** | hsp70.3 | 90.29752 | 5.169946194 | 0 | 0 |
| **54** | mycb | 89.90348 | 3.781744003 | 0 | 0 |
| **55** | pfkfb3 | 89.82876 | 4.429637909 | 0 | 0 |
| **56** | jund | 89.80586 | 3.046759844 | 0 | 0 |
| **57** | gulp1a | 88.92162 | 4.507949829 | 0 | 0 |
| **58** | dhrs13l1 | 87.82293 | 4.125034332 | 0 | 0 |
| **59** | tob1b | 87.59043 | 3.636733055 | 0 | 0 |
| **60** | gpm6bb | 87.52592 | 4.23235178 | 0 | 0 |
| **61** | glulb | 86.93468 | 2.942313433 | 0 | 0 |
| **62** | nrgna | 86.5343 | 3.377605915 | 0 | 0 |
| **63** | hsp90aa1.2 | 85.88276 | 4.621142387 | 0 | 0 |
| **64** | itm2ba | 85.22511 | 1.974311709 | 0 | 0 |
| **65** | hyal6 | 84.91393 | 4.75174427 | 0 | 0 |
| **66** | CU929418.2 | 84.85066 | 3.718532324 | 0 | 0 |
| **67** | tob1a | 83.87309 | 3.637747049 | 0 | 0 |
| **68** | si:dkey-56f14.7 | 82.75861 | 3.836972952 | 0 | 0 |
| **69** | lin7a | 82.7543 | 4.844590664 | 0 | 0 |
| **70** | gyg1b | 82.66338 | 4.9946208 | 0 | 0 |
| **71** | f3b | 82.60305 | 3.78447938 | 0 | 0 |
| **72** | csrp2 | 82.12587 | 4.869801998 | 0 | 0 |
| **73** | gpd1b | 81.33405 | 5.089282513 | 0 | 0 |
| **74** | zgc:153704 | 80.99913 | 5.608815193 | 0 | 0 |
| **75** | vat1 | 80.49884 | 2.980295897 | 0 | 0 |
| **76** | gadd45ba | 80.35832 | 2.952223539 | 0 | 0 |
| **77** | gapdhs | 79.81317 | 1.74020493 | 0 | 0 |
| **78** | tmem176l.1 | 78.8969 | 2.263581991 | 0 | 0 |
| **79** | zgc:92606 | 78.59956 | 2.632120371 | 0 | 0 |
| **80** | rtn4a | 78.45934 | 1.951094747 | 0 | 0 |
| **81** | cnn3a | 78.35567 | 2.745401144 | 0 | 0 |
| **82** | ppp1r15a | 77.76933 | 3.812073708 | 0 | 0 |
| **83** | gpm6ab | 77.76002 | 2.387986183 | 0 | 0 |
| **84** | zgc:195173 | 77.67789 | 4.041749001 | 0 | 0 |
| **85** | scarb2a | 77.54933 | 2.747274637 | 0 | 0 |
| **86** | espn | 77.12132 | 4.101651669 | 0 | 0 |
| **87** | alkbh3-1 | 76.7421 | 3.688959837 | 0 | 0 |
| **88** | hsd11b2 | 76.62229 | 4.310726643 | 0 | 0 |
| **89** | CR383676.1 | 76.35896 | 1.118119597 | 0 | 0 |
| **90** | anks4b | 75.93623 | 4.868053913 | 0 | 0 |
| **91** | si:ch1073-303k11.2 | 75.85842 | 3.036639452 | 0 | 0 |
| **92** | dnajb1b | 75.45119 | 4.250008583 | 0 | 0 |
| **93** | clstn1 | 75.32479 | 2.615636587 | 0 | 0 |
| **94** | cebpd | 74.94839 | 4.392705917 | 0 | 0 |
| **95** | map4l | 74.88611 | 3.423485041 | 0 | 0 |
| **96** | hmgcs1 | 74.86773 | 4.08097887 | 0 | 0 |
| **97** | hsp70.1 | 74.67686 | 4.996319771 | 0 | 0 |
| **98** | slc43a2b | 74.32397 | 4.363460541 | 0 | 0 |
| **99** | acbd7 | 73.97585 | 2.28858614 | 0 | 0 |

**Cluster 2**

| | names | scores | logfoldchanges | pvals | pvals_adj |
|---|---|---|---|---|---|
| 0 | apoeb | 106.334 | 4.605007648 | 0 | 0 |
| 1 | fosab | 68.48776 | 3.710303307 | 2.6E-284 | 2E-281 |
| 2 | glula | 62.38844 | 3.501479864 | 1.2E-248 | 6.4E-246 |
| 3 | rlbp1a | 62.38046 | 3.2717309 | 1.1E-235 | 4.6E-233 |
| 4 | fosb | 53.34531 | 3.075199604 | 1.6E-192 | 4E-190 |
| 5 | nfasca | 52.18295 | 2.64123559 | 7.8E-192 | 1.9E-189 |
| 6 | slc1a2b | 51.11511 | 2.606746197 | 2.9E-196 | 7.5E-194 |
| 7 | sparc | 50.24355 | 2.870394707 | 3.4E-180 | 7E-178 |
| 8 | cahz | 48.28159 | 2.865404606 | 5E-190 | 1.2E-187 |
| 9 | mt2 | 46.54424 | 4.235199928 | 1.7E-162 | 2.8E-160 |
| 10 | jdp2b | 44.74947 | 3.268232346 | 5.8E-164 | 9.8E-162 |
| 11 | btg2 | 41.3894 | 2.187958717 | 1.5E-156 | 2.1E-154 |
| 12 | chrdl2 | 41.05654 | 5.588739395 | 6.7E-135 | 6.1E-133 |
| 13 | rdh10a | 40.69147 | 5.249956608 | 4.8E-134 | 4.3E-132 |
| 14 | jun | 40.10576 | 2.230064869 | 2.3E-149 | 2.7E-147 |
| 15 | zfp36l1b | 39.33615 | 2.821576118 | 2.1E-138 | 2E-136 |
| 16 | selenop | 38.59664 | 2.610793352 | 1.4E-141 | 1.5E-139 |
| 17 | sncga | 38.21304 | 3.161438704 | 5.7E-130 | 4.8E-128 |
| 18 | atp1a1b | 36.81063 | 2.349395037 | 7.9E-132 | 6.9E-130 |
| 19 | gstp1 | 34.9228 | 2.291466475 | 5.4E-127 | 4.4E-125 |
| 20 | fxyd6l | 34.06401 | 1.8824296 | 9.1E-124 | 7E-122 |
| 21 | atp1b3a | 34.0034 | 2.600070715 | 2.1E-116 | 1.5E-114 |
| 22 | aldh1a3 | 33.464 | 4.511420727 | 9E-111 | 5.9E-109 |
| 23 | ptgdsb.2 | 32.2485 | 2.588564873 | 3.1E-114 | 2.1E-112 |
| 24 | cdo1 | 31.22018 | 2.199245214 | 3.4E-109 | 2.1E-107 |
| 25 | zgc:195173 | 29.73603 | 2.713517904 | 1.2E-100 | 6.5E-99 |
| 26 | nrgna | 28.88848 | 2.129214764 | 5.08E-98 | 2.76E-96 |
| 27 | aplp2 | 28.61648 | 1.900037289 | 5.36E-98 | 2.91E-96 |
| 28 | jund | 28.43835 | 1.990542293 | 9.79E-97 | 5.21E-95 |
| 29 | CU467822.1 | 28.16533 | 1.877335429 | 2.17E-95 | 1.13E-93 |
| 30 | tob1a | 28.04329 | 2.44776845 | 2.06E-93 | 1.04E-91 |
| 31 | stc2a | 27.92295 | 3.924168348 | 1.04E-91 | 5.1E-90 |
| 32 | zgc:153704 | 27.90702 | 2.947695494 | 1.79E-93 | 9.05E-92 |
| 33 | si:dkey-164f24.2 | 27.81779 | 3.10580492 | 2.96E-91 | 1.43E-89 |
| 34 | f3b | 27.6234 | 2.782377243 | 3.66E-92 | 1.8E-90 |
| 35 | eno1a | 27.36835 | 1.494156957 | 6.32E-96 | 3.34E-94 |
| 36 | ppp1r14aa | 27.29414 | 2.183029413 | 1.32E-91 | 6.43E-90 |
| 37 | si:ch211-152c2.3 | 27.2213 | 2.457709312 | 1.02E-92 | 5.07E-91 |
| 38 | junbb | 27.09918 | 2.1152637 | 5.76E-91 | 2.77E-89 |
| 39 | socs3a | 27.0586 | 2.826738119 | 5.78E-90 | 2.72E-88 |
| 40 | hspb1 | 26.31068 | 3.335918665 | 2.92E-86 | 1.31E-84 |
| 41 | s100a10b | 25.6166 | 1.922732711 | 2.63E-86 | 1.18E-84 |
| 42 | mcl1a | 25.44027 | 2.156741381 | 1.41E-84 | 6.06E-83 |
| 43 | efhd1 | 25.02043 | 1.881289244 | 7.04E-83 | 2.97E-81 |
| 44 | junba | 24.98672 | 1.958903074 | 1.18E-82 | 4.94E-81 |
| 45 | egr1 | 24.31924 | 2.771734953 | 6.89E-78 | 2.7E-76 |
| 46 | clstn1 | 23.9734 | 1.831335425 | 2.58E-77 | 1.01E-75 |
| 47 | mstnb | 23.95243 | 4.543766975 | 2.23E-75 | 8.39E-74 |
| 48 | fosl1a | 23.76232 | 2.227057695 | 6.55E-77 | 2.54E-75 |
| 49 | slc38a4 | 23.59912 | 2.157702446 | 1.06E-75 | 4.02E-74 |
| 50 | epas1b | 23.56985 | 2.195223093 | 1.75E-75 | 6.61E-74 |
| 51 | sik2b | 22.82044 | 2.199272394 | 3.89E-72 | 1.4E-70 |
| 52 | si:dkey-16p21.8 | 22.3226 | 1.943249702 | 1.3E-70 | 4.48E-69 |
| 53 | rsrp1 | 21.95772 | 1.676834702 | 4.46E-69 | 1.49E-67 |
| 54 | ptmaa | 21.81192 | 1.292658091 | 9.55E-70 | 3.23E-68 |
| 55 | pik3ip1 | 21.65113 | 1.894111395 | 2.76E-68 | 9.08E-67 |
| 56 | cavin2a | 21.4658 | 2.047474861 | 6.24E-67 | 2E-65 |
| 57 | prdx6 | 21.35275 | 1.577369809 | 3.09E-67 | 9.95E-66 |
| 58 | dab2ipa | 21.30005 | 1.702703357 | 1.78E-66 | 5.64E-65 |
| 59 | mcl1b | 21.23481 | 1.918084979 | 3.29E-66 | 1.04E-64 |
| 60 | gyg1b | 21.21754 | 2.125426292 | 8.7E-66 | 2.72E-64 |
| 61 | gulp1a | 20.91881 | 2.07951808 | 1.94E-64 | 5.9E-63 |
| 62 | bgnb | 20.61047 | 2.36639452 | 8.28E-63 | 2.44E-61 |
| 63 | rbpms2b | 20.46741 | 1.680673957 | 5.11E-63 | 1.51E-61 |
| 64 | dhrs13l1 | 19.99698 | 1.883863926 | 1.15E-60 | 3.29E-59 |
| 65 | insig1 | 19.91047 | 1.677845478 | 4.61E-61 | 1.33E-59 |
| 66 | pfkfb3 | 19.65537 | 2.199845314 | 4.34E-59 | 1.21E-57 |
| 67 | slc4a4a | 19.38198 | 2.021410227 | 7.17E-58 | 1.95E-56 |
| 68 | dusp2 | 19.2915 | 2.530094862 | 2.47E-57 | 6.65E-56 |
| 69 | mCherry | 19.25415 | 1.687426805 | 4.87E-58 | 1.33E-56 |
| 70 | hopx | 19.24791 | 2.067633152 | 2.24E-57 | 6.04E-56 |
| 71 | btg1 | 18.99416 | 1.188444495 | 7.23E-57 | 1.92E-55 |
| 72 | spock3 | 18.86499 | 1.408797503 | 3.18E-57 | 8.52E-56 |
| 73 | cebpd | 18.65459 | 2.186013222 | 6.72E-55 | 1.72E-53 |
| 74 | mycb | 18.38472 | 2.029505014 | 6.12E-54 | 1.54E-52 |
| 75 | tob1b | 18.14173 | 1.812660456 | 5.46E-53 | 1.35E-51 |
| 76 | col15a1b | 18.06737 | 1.744888663 | 7.38E-53 | 1.82E-51 |
| 77 | gpd1b | 17.80615 | 1.976151824 | 2.01E-51 | 4.81E-50 |
| 78 | slc43a2b | 17.60483 | 2.06780386 | 2.03E-50 | 4.73E-49 |
| 79 | ndrg3a | 17.5949 | 2.007047415 | 2.4E-50 | 5.6E-49 |
| 80 | ivns1abpa | 17.35951 | 1.565198064 | 1.41E-49 | 3.24E-48 |
| 81 | alkbh3-1 | 17.25975 | 1.818648338 | 2.55E-49 | 5.82E-48 |
| 82 | CU929418.2 | 17.22066 | 1.727148533 | 5.2E-49 | 1.18E-47 |
| 83 | espn | 17.01364 | 1.804711223 | 3.76E-48 | 8.37E-47 |
| 84 | itm2ba | 16.92506 | 1.058800936 | 3.64E-48 | 8.11E-47 |
| 85 | map4l | 16.77126 | 1.681953549 | 3.57E-47 | 7.72E-46 |
| 86 | cav1 | 16.64918 | 2.327598572 | 2.45E-46 | 5.22E-45 |
| 87 | csrp2 | 16.62286 | 1.980470061 | 2.09E-46 | 4.45E-45 |
| 88 | rhbg | 16.41214 | 1.405569553 | 4.45E-46 | 9.41E-45 |
| 89 | gnai2b | 16.38775 | 2.707526445 | 3.58E-45 | 7.41E-44 |
| 90 | dap1b | 16.18257 | 1.355489254 | 5.09E-45 | 1.05E-43 |
| 91 | hsp90aa1.2 | 16.12121 | 2.22889924 | 1.04E-44 | 2.13E-43 |
| 92 | pim1 | 16.09024 | 1.19142127 | 5.05E-45 | 1.04E-43 |
| 93 | nocta | 15.97082 | 1.431319594 | 4.53E-44 | 9.1E-43 |
| 94 | tmem176l.1 | 15.77423 | 1.026697874 | 1.41E-43 | 2.79E-42 |
| 95 | im:7152348 | 15.72226 | 2.020355463 | 1.22E-42 | 2.36E-41 |
| 96 | rtn4a | 15.69392 | 0.852889717 | 1.14E-43 | 2.26E-42 |
| 97 | vegfaa | 15.52684 | 1.801396132 | 6.71E-42 | 1.26E-40 |
| 98 | mdka | 15.52096 | 0.896473527 | 4.28E-43 | 8.41E-42 |
| 99 | rpz5 | 15.33037 | 1.8974365 | 3.62E-41 | 6.64E-40 |

**Cluster 3**

| | names | scores | logfoldchanges | pvals | pvals_adj |
|---|---|---|---|---|---|
| **0** | her15.1-1 | 58.02736 | 4.225009918 | 0 | 0 |
| **1** | si:ch73-335l21.4 | 57.05906 | 3.122822285 | 0 | 0 |
| **2** | cxcl18b | 52.75472 | 4.977982521 | 6.5E-285 | 3E-281 |
| **3** | marcksl1b | 52.34999 | 2.360349655 | 0 | 8.2E-306 |
| **4** | mmp9 | 50.80667 | 4.913655758 | 4.3E-273 | 1.2E-269 |
| **5** | stm | 48.47406 | 8.312149048 | 2.7E-255 | 5.4E-252 |
| **6** | cdh2 | 45.71982 | 1.957092166 | 2.5E-256 | 5.4E-253 |
| **7** | rplp1 | 45.05664 | 1.426795006 | 1E-279 | 3.2E-276 |
| **8** | sox4a-1 | 44.04139 | 3.711538792 | 2.6E-233 | 3.2E-230 |
| **9** | txn | 43.90092 | 3.346035957 | 1.3E-235 | 1.7E-232 |
| **10** | her9 | 43.68443 | 3.637079 | 1.9E-232 | 2.2E-229 |
| **11** | si:dkey-151g10.6 | 42.14235 | 1.381598711 | 8.2E-247 | 1.3E-243 |
| **12** | fosab | 41.71621 | 2.003262281 | 2.2E-282 | 7.8E-279 |
| **13** | si:ch211-222l21.1 | 41.42971 | 1.932316303 | 3.7E-243 | 5.7E-240 |
| **14** | rplp2l | 41.2953 | 1.51043427 | 7.7E-243 | 1.1E-239 |
| **15** | pim1 | 41.13966 | 1.882244825 | 4.6E-237 | 6.3E-234 |
| **16** | ncl | 41.0663 | 2.076798201 | 1.1E-220 | 8.8E-218 |
| **17** | id1 | 39.84898 | 2.902092695 | 1.3E-213 | 8.9E-211 |
| **18** | rps11 | 39.72352 | 1.308057666 | 5.5E-228 | 5.8E-225 |
| **19** | hspa5 | 39.58265 | 2.044267178 | 4.2E-221 | 3.5E-218 |
| **20** | hbegfa | 39.39593 | 3.19938302 | 8.1E-205 | 4.8E-202 |
| **21** | rpl27 | 39.37078 | 1.390632391 | 2.8E-222 | 2.5E-219 |
| **22** | rtn4a | 39.29338 | 1.448977113 | 5.9E-224 | 5.4E-221 |
| **23** | rps14 | 39.26819 | 1.29278338 | 2.2E-225 | 2.2E-222 |
| **24** | six3b | 38.62426 | 1.994689345 | 1.6E-209 | 9.8E-207 |
| **25** | rps8a | 38.4972 | 1.238654375 | 2.8E-224 | 2.7E-221 |
| **26** | rpl23 | 38.42531 | 1.288289309 | 2.4E-220 | 1.9E-217 |
| **27** | rpl9 | 38.19319 | 1.353270411 | 7.3E-217 | 5.6E-214 |
| **28** | rps27a | 37.9284 | 1.250830531 | 4.8E-215 | 3.6E-212 |
| **29** | rps25 | 37.88544 | 1.387264848 | 1E-211 | 6.6E-209 |
| **30** | rps15a | 37.86848 | 1.308765411 | 9E-213 | 6.1E-210 |
| **31** | rps23 | 37.85899 | 1.287877679 | 2.6E-214 | 1.9E-211 |
| **32** | rplp0 | 37.41143 | 1.22599256 | 3E-214 | 2.1E-211 |
| **33** | rps26l | 37.09454 | 1.370550513 | 8.7E-204 | 5E-201 |
| **34** | her15.1 | 36.9349 | 3.666859627 | 1.7E-186 | 6.8E-184 |
| **35** | rpl36a | 36.76678 | 1.341036201 | 1.7E-201 | 9.4E-199 |
| **36** | foxj1a | 36.76667 | 3.513093472 | 1.5E-185 | 5.4E-183 |
| **37** | uba52 | 36.4925 | 1.22524631 | 2.3E-200 | 1.2E-197 |
| **38** | tomm20a | 36.4276 | 2.381071091 | 2.4E-185 | 8.8E-183 |
| **39** | zgc:158343 | 36.32206 | 2.799567223 | 6.7E-188 | 2.8E-185 |
| **40** | rps2 | 36.2913 | 1.176840901 | 9.9E-207 | 6.1E-204 |
| **41** | her12 | 36.14827 | 4.560902596 | 3E-180 | 9.7E-178 |
| **42** | rps15 | 35.9535 | 1.225199699 | 1.5E-197 | 7.9E-195 |
| **43** | hsp90aa1.2 | 35.66359 | 2.108299017 | 2.3E-205 | 1.4E-202 |
| **44** | myl6 | 35.63562 | 2.198964357 | 3.6E-182 | 1.2E-179 |
| **45** | rps24 | 35.56178 | 1.204318881 | 1.3E-194 | 6.7E-192 |
| **46** | her4.2 | 35.55307 | 3.947669744 | 4E-177 | 1.2E-174 |
| **47** | rpl35 | 35.52575 | 1.269109488 | 8.3E-191 | 3.7E-188 |
| **48** | eef2b | 35.24443 | 1.252345443 | 1.7E-187 | 7E-185 |
| **49** | rps19 | 35.18133 | 1.196424603 | 5.4E-194 | 2.6E-191 |
| **50** | rpl17 | 35.08038 | 1.169332743 | 1.2E-193 | 5.8E-191 |
| **51** | rpl32 | 35.0037 | 1.212828875 | 4.3E-191 | 1.9E-188 |
| **52** | her6 | 34.67548 | 2.641201735 | 3.1E-176 | 9.1E-174 |
| **53** | mych | 34.61427 | 2.114996672 | 1.1E-177 | 3.4E-175 |
| **54** | rps10 | 34.58969 | 1.129958391 | 3.3E-191 | 1.5E-188 |
| **55** | cd63 | 34.58833 | 2.145318508 | 2E-173 | 5.3E-171 |
| **56** | TXN | 34.55949 | 1.499872923 | 5.4E-178 | 1.7E-175 |
| **57** | rpl28 | 34.5156 | 1.169592142 | 3.8E-187 | 1.5E-184 |
| **58** | rps13 | 34.39772 | 1.18578136 | 8.7E-186 | 3.2E-183 |
| **59** | rps7 | 34.37388 | 1.129494667 | 6.1E-189 | 2.6E-186 |
| **60** | rpsa | 34.21595 | 1.146769404 | 2.6E-186 | 1E-183 |
| **61** | rpl19 | 34.15697 | 1.127461433 | 1.8E-187 | 7.4E-185 |
| **62** | sgk1 | 34.14804 | 2.012307405 | 6.1E-175 | 1.7E-172 |
| **63** | rpl7 | 34.12462 | 1.149122 | 9E-185 | 3.2E-182 |
| **64** | rps5 | 34.07727 | 1.135283351 | 3.8E-187 | 1.5E-184 |
| **65** | RPS17 | 34.06856 | 1.173175216 | 5.9E-181 | 2E-178 |
| **66** | her4.1 | 34.03872 | 4.49159956 | 2.2E-166 | 4.7E-164 |
| **67** | rps9 | 33.97892 | 1.105675817 | 5.1E-186 | 1.9E-183 |
| **68** | rpl14 | 33.95321 | 1.165776968 | 3.3E-179 | 1E-176 |
| **69** | rpl36 | 33.75732 | 1.332930803 | 6.2E-175 | 1.7E-172 |
| **70** | atp1b1a | 33.72487 | 1.963865161 | 5.5E-171 | 1.3E-168 |
| **71** | rps12 | 33.72482 | 1.174200892 | 5.5E-182 | 1.9E-179 |
| **72** | rpl18a | 33.65269 | 1.162939429 | 2.6E-180 | 8.4E-178 |
| **73** | krt8 | 33.58528 | 2.3538661 | 2.1E-168 | 4.6E-166 |
| **74** | rpl23a | 33.49052 | 1.208750963 | 1.7E-176 | 5.2E-174 |
| **75** | ier2b | 33.46938 | 2.473191261 | 2.5E-166 | 5.3E-164 |
| **76** | hsp70.3 | 33.42881 | 2.135173321 | 7.8E-181 | 2.5E-178 |
| **77** | cirbpa | 33.37631 | 1.243928194 | 3.2E-175 | 9.1E-173 |
| **78** | si:dkey-7j14.6 | 33.31284 | 1.477175593 | 1.7E-171 | 4.2E-169 |
| **79** | rpl35a | 33.18785 | 1.189859033 | 3.8E-174 | 1E-171 |
| **80** | nme2b.1 | 33.15488 | 1.203778625 | 6.1E-174 | 1.6E-171 |
| **81** | RPL37A | 33.13859 | 1.198282361 | 5.6E-173 | 1.4E-170 |
| **82** | rpl13 | 33.09223 | 1.106846213 | 6.8E-177 | 2E-174 |
| **83** | hsp90ab1 | 33.00584 | 0.964442253 | 8.4E-171 | 2E-168 |
| **84** | crlf1a | 32.99549 | 2.786085606 | 3.8E-163 | 7.8E-161 |
| **85** | eef1a1l1 | 32.97499 | 0.900468767 | 3.6E-173 | 9.2E-171 |
| **86** | tubb4b | 32.90364 | 1.166003585 | 5.3E-169 | 1.2E-166 |
| **87** | rpl10 | 32.86886 | 1.094585538 | 2.9E-175 | 8.2E-173 |
| **88** | rpl31 | 32.85209 | 1.156771302 | 7.7E-171 | 1.8E-168 |
| **89** | rpl13a | 32.73288 | 1.072018266 | 5.6E-172 | 1.4E-169 |
| **90** | rpl39 | 32.71383 | 1.118948698 | 3.5E-171 | 8.5E-169 |
| **91** | rps3a | 32.60934 | 1.066306233 | 1.7E-174 | 4.5E-172 |
| **92** | rpl7a | 32.46098 | 1.091044545 | 6.2E-173 | 1.5E-170 |
| **93** | her4.2-1 | 32.34126 | 4.645433903 | 2.1E-154 | 3.9E-152 |
| **94** | vmp1 | 32.28925 | 2.58438158 | 1.1E-157 | 2.1E-155 |
| **95** | rpl21 | 32.22758 | 1.119826913 | 5.3E-169 | 1.2E-166 |
| **96** | rpl10a | 32.224 | 1.058582544 | 1.9E-170 | 4.3E-168 |
| **97** | ppdpfb | 32.20901 | 1.185377955 | 1.5E-166 | 3.4E-164 |
| **98** | rps4x | 32.0587 | 1.095569491 | 3.9E-169 | 8.9E-167 |
| **99** | rpl15 | 31.79554 | 1.086902499 | 2.2E-168 | 4.9E-166 |

**Cluster 4**

| | names | scores | logfoldchanges | pvals | pvals_adj |
|---|---|---|---|---|---|
| **0** | f3a | 55.36212 | 3.084530592 | 4.8E-282 | 6E-279 |
| **1** | crlf1a | 48.00781 | 4.067601204 | 2.3E-230 | 1.9E-227 |
| **2** | fabp7a | 45.29711 | 1.886966348 | 2.9E-248 | 2.7E-245 |
| **3** | mdka | 43.44316 | 1.958930135 | 1.6E-225 | 1.3E-222 |
| **4** | vmp1 | 42.79474 | 2.970813513 | 5.3E-208 | 3.5E-205 |
| **5** | marcksl1a | 42.75191 | 2.058850288 | 9.1E-211 | 6.3E-208 |
| **6** | crabp1a | 36.21816 | 1.700720549 | 5.3E-175 | 2.5E-172 |
| **7** | clcf1 | 35.68691 | 3.765312433 | 1.5E-161 | 5.3E-159 |
| **8** | myl6 | 35.67439 | 2.276098728 | 4.5E-166 | 1.7E-163 |
| **9** | txn | 33.94541 | 3.097076654 | 1.9E-153 | 6.1E-151 |
| **10** | hbegfa | 33.53465 | 3.055268526 | 6.5E-151 | 2E-148 |
| **11** | cdh2 | 31.51853 | 1.473041654 | 4.7E-144 | 1.3E-141 |
| **12** | zic2b | 31.50673 | 3.065355301 | 1.7E-138 | 4.2E-136 |
| **13** | akap12b | 31.32176 | 2.762203693 | 1.5E-137 | 3.8E-135 |
| **14** | ppdpfb | 30.08066 | 1.160496831 | 2.7E-139 | 6.8E-137 |
| **15** | flna | 29.97908 | 2.376906633 | 2.4E-130 | 5.2E-128 |
| **16** | cd82a | 29.05429 | 1.853296876 | 3.1E-126 | 6.1E-124 |
| **17** | six3b | 29.03836 | 1.83422184 | 1.5E-127 | 3.1E-125 |
| **18** | dusp5 | 28.49575 | 2.876734734 | 6.6E-121 | 1.2E-118 |
| **19** | cnn2 | 28.39768 | 2.205649614 | 6.5E-122 | 1.2E-119 |
| **20** | acbd7 | 28.14837 | 1.591598868 | 6.3E-123 | 1.2E-120 |
| **21** | rpl12 | 27.13593 | 0.976159751 | 4.4E-121 | 8.1E-119 |
| **22** | gfap | 26.47637 | 1.711139798 | 4.3E-111 | 6.8E-109 |
| **23** | si:ch211-251b21.1 | 26.15443 | 3.095375776 | 1.2E-106 | 1.8E-104 |
| **24** | CR383676.1 | 26.15266 | 0.728779912 | 1.7E-112 | 2.7E-110 |
| **25** | marcksl1b | 26.13877 | 1.290313482 | 1.6E-112 | 2.7E-110 |
| **26** | rpl9 | 25.55742 | 1.026162028 | 6.8E-110 | 1.1E-107 |
| **27** | si:dkey-238o13.4 | 24.85637 | 2.669303894 | 1E-99 | 1.37E-97 |
| **28** | lepb | 24.81931 | 3.228264093 | 7.7E-99 | 1.01E-96 |
| **29** | ckbb | 24.4378 | 1.224566102 | 2E-100 | 2.73E-98 |
| **30** | bzw1b | 24.03344 | 1.352009296 | 1.85E-97 | 2.35E-95 |
| **31** | tgif1 | 23.72293 | 1.649936795 | 8.82E-94 | 1.01E-91 |
| **32** | mvp | 23.71933 | 1.615901113 | 3.57E-94 | 4.13E-92 |
| **33** | rasgef1bb | 23.52336 | 1.74011147 | 1.99E-92 | 2.18E-90 |
| **34** | nme2b.1 | 23.25942 | 0.99786979 | 1.5E-93 | 1.7E-91 |
| **35** | rps2 | 22.98646 | 0.882615745 | 1.06E-93 | 1.21E-91 |
| **36** | col18a1a | 22.78006 | 2.001247644 | 1.45E-87 | 1.45E-85 |
| **37** | elovl1b | 22.62617 | 2.455638885 | 2.51E-86 | 2.44E-84 |
| **38** | rplp0 | 22.11063 | 0.830475926 | 3.82E-88 | 3.86E-86 |
| **39** | zgc:165604 | 21.60006 | 1.371934414 | 1.68E-82 | 1.5E-80 |
| **40** | mdkb | 21.37778 | 1.696121693 | 5.39E-80 | 4.67E-78 |
| **41** | tjp2b | 21.34831 | 2.25809598 | 4.89E-79 | 4.11E-77 |
| **42** | cd81a | 20.93864 | 1.194727063 | 9.01E-78 | 7.32E-76 |
| **43** | atp1b1a | 20.68841 | 1.644294381 | 7.48E-76 | 5.85E-74 |
| **44** | krt8 | 20.54859 | 1.769300938 | 3.38E-75 | 2.58E-73 |
| **45** | sgk1 | 20.44717 | 1.40393579 | 2.52E-75 | 1.93E-73 |
| **46** | si:dkey-7j14.6 | 20.43421 | 1.045555592 | 8.56E-76 | 6.68E-74 |
| **47** | myl9b | 19.57335 | 1.285054445 | 5.23E-70 | 3.53E-68 |
| **48** | rpl19 | 19.47244 | 0.725687742 | 1.04E-71 | 7.27E-70 |
| **49** | rpl7a | 19.40669 | 0.719799817 | 2.43E-71 | 1.68E-69 |
| **50** | si:ch1073-303k11.2 | 19.36223 | 1.230882287 | 1.51E-69 | 1.01E-67 |
| **51** | sall1b | 19.31191 | 3.526893139 | 4.3E-67 | 2.7E-65 |
| **52** | rplp1 | 19.22264 | 0.731488287 | 4.53E-70 | 3.06E-68 |
| **53** | mmp9 | 19.20126 | 2.362295389 | 4.2E-67 | 2.64E-65 |
| **54** | rtn4a | 19.05221 | 0.876076937 | 3.93E-68 | 2.53E-66 |
| **55** | c7b | 18.77744 | 2.772310495 | 2.14E-64 | 1.29E-62 |
| **56** | fam129bb | 18.77638 | 2.289850235 | 1.91E-64 | 1.16E-62 |
| **57** | nrp2b | 18.6152 | 1.960113049 | 1.01E-63 | 5.97E-62 |
| **58** | pim1 | 18.60892 | 0.977517307 | 6.05E-66 | 3.74E-64 |
| **59** | tpt1 | 18.462 | 0.726840198 | 1.35E-64 | 8.2E-63 |
| **60** | gpm6aa | 18.2658 | 0.829822958 | 2.7E-63 | 1.57E-61 |
| **61** | rps9 | 18.14201 | 0.692920327 | 1.85E-63 | 1.09E-61 |
| **62** | aplp1 | 18.09368 | 1.95404613 | 7.77E-61 | 4.25E-59 |
| **63** | igfbp5b | 18.02501 | 2.394520521 | 2.58E-60 | 1.37E-58 |
| **64** | yap1 | 17.98789 | 1.520968795 | 1.63E-60 | 8.77E-59 |
| **65** | zgc:86709 | 17.95627 | 3.522382021 | 1.02E-59 | 5.28E-58 |
| **66** | rpl32 | 17.87215 | 0.67593658 | 6.02E-62 | 3.4E-60 |
| **67** | pros1 | 17.85444 | 1.845410347 | 1.62E-59 | 8.35E-58 |
| **68** | zgc:153867 | 17.84432 | 0.912852943 | 2.91E-60 | 1.54E-58 |
| **69** | rps14 | 17.80692 | 0.660975337 | 1.87E-61 | 1.04E-59 |
| **70** | rps3a | 17.74341 | 0.665450156 | 3.27E-61 | 1.8E-59 |
| **71** | grb10b | 17.70638 | 1.505983472 | 6.4E-59 | 3.23E-57 |
| **72** | myl12.1 | 17.66671 | 0.779660761 | 1.73E-59 | 8.92E-58 |
| **73** | il11b | 17.64211 | 4.81022644 | 6.46E-58 | 3.19E-56 |
| **74** | lgals2a | 17.63276 | 2.190256596 | 2.21E-58 | 1.1E-56 |
| **75** | pprc1 | 17.50554 | 1.910521984 | 1.08E-57 | 5.28E-56 |
| **76** | rack1 | 17.49477 | 0.660355985 | 7.32E-60 | 3.82E-58 |
| **77** | irf2 | 17.48677 | 2.837457895 | 2.75E-57 | 1.34E-55 |
| **78** | tpm4a | 17.30012 | 2.23798275 | 1.81E-56 | 8.62E-55 |
| **79** | eef1g | 17.23352 | 0.694715261 | 2.84E-58 | 1.41E-56 |
| **80** | cnn3a | 17.06765 | 1.188982844 | 6.57E-56 | 3.08E-54 |
| **81** | rps11 | 17.03423 | 0.663609684 | 9.59E-57 | 4.63E-55 |
| **82** | rpl11 | 17.00854 | 0.683452547 | 1.02E-56 | 4.92E-55 |
| **83** | rps10 | 16.9364 | 0.657839417 | 1.83E-56 | 8.69E-55 |
| **84** | rpl36a | 16.53119 | 0.678871751 | 7.15E-54 | 3.17E-52 |
| **85** | rpl23 | 16.49108 | 0.650538802 | 7.07E-54 | 3.14E-52 |
| **86** | rps8a | 16.48499 | 0.622688115 | 4.62E-54 | 2.06E-52 |
| **87** | fabp3 | 16.45695 | 0.90865171 | 5.86E-53 | 2.55E-51 |
| **88** | si:dkey-79d12.5 | 16.39497 | 1.804925442 | 9.38E-52 | 3.91E-50 |
| **89** | mych | 16.39429 | 1.198716283 | 2.18E-52 | 9.28E-51 |
| **90** | klf7b | 16.3875 | 1.336815 | 5.6E-52 | 2.36E-50 |
| **91** | actb1 | 16.37879 | 0.509622693 | 3.39E-53 | 1.49E-51 |
| **92** | nat8l | 16.33082 | 1.99587059 | 2.45E-51 | 1.01E-49 |
| **93** | fosl2 | 16.27498 | 1.075386524 | 1.59E-51 | 6.6E-50 |
| **94** | npdc1b | 16.27261 | 2.002556562 | 4.74E-51 | 1.93E-49 |
| **95** | rps15a | 16.17297 | 0.677108884 | 7.86E-52 | 3.29E-50 |
| **96** | kitlgb | 16.02795 | 3.64020896 | 1.57E-49 | 6.14E-48 |
| **97** | dbn1 | 15.98582 | 1.42545867 | 1.08E-49 | 4.22E-48 |
| **98** | rps12 | 15.98538 | 0.654640138 | 5.07E-51 | 2.06E-49 |
| **99** | rpl8 | 15.96231 | 0.584302187 | 4.56E-51 | 1.86E-49 |

**Cluster 5**

| | names | scores | logfoldchanges | pvals | pvals_adj |
|---|---|---|---|---|---|
| 0 | hmgn2 | 68.39232 | 2.729560614 | 0 | 0 |
| 1 | hmgb2b | 63.28001 | 2.253685951 | 0 | 0 |
| 2 | rpl9 | 61.13248 | 1.683425903 | 0 | 0 |
| 3 | rpl12 | 60.21627 | 1.503011346 | 0 | 0 |
| 4 | rplp0 | 59.62198 | 1.53966856 | 0 | 0 |
| 5 | rplp1 | 59.1498 | 1.471200943 | 0 | 0 |
| 6 | rps2 | 56.12161 | 1.469377637 | 0 | 0 |
| 7 | h2afvb | 55.04961 | 1.873373985 | 0 | 0 |
| 8 | rps9 | 54.23639 | 1.381539941 | 0 | 0 |
| 9 | rplp2l | 54.04665 | 1.505197406 | 0 | 0 |
| 10 | rps12 | 53.88076 | 1.405890107 | 0 | 0 |
| 11 | rps3a | 53.59622 | 1.372345328 | 0 | 0 |
| 12 | rpl23 | 53.5159 | 1.383931756 | 0 | 0 |
| 13 | rpl10a | 53.37649 | 1.363327026 | 0 | 0 |
| 14 | rps10 | 52.57711 | 1.35702312 | 0 | 0 |
| 15 | rpl7a | 52.35942 | 1.369402647 | 0 | 0 |
| 16 | rpl11 | 52.08962 | 1.363907933 | 0 | 0 |
| 17 | rpl3 | 51.9273 | 1.341996551 | 0 | 0 |
| 18 | rps8a | 51.79038 | 1.363577724 | 0 | 0 |
| 19 | rpl32 | 51.48996 | 1.337257028 | 0 | 0 |
| 20 | si:dkey-151g10.6 | 51.31477 | 1.310818791 | 0 | 0 |
| 21 | rpl15 | 51.17108 | 1.409645677 | 0 | 0 |
| 22 | rpsa | 50.93291 | 1.367236614 | 0 | 0 |
| 23 | rpl10 | 50.88318 | 1.30215776 | 0 | 0 |
| 24 | hmgb2a | 50.71026 | 2.166218996 | 0 | 0 |
| 25 | rpl21 | 50.68056 | 1.339955926 | 0 | 0 |
| 26 | rps7 | 50.56798 | 1.311057687 | 0 | 0 |
| 27 | rps19 | 50.27814 | 1.334269285 | 0 | 0 |
| 28 | ran | 49.99977 | 1.585142136 | 0 | 0 |
| 29 | rpl8 | 49.80364 | 1.286369324 | 0 | 0 |
| 30 | eef1g | 49.54037 | 1.367532253 | 0 | 0 |
| 31 | si:ch211-222l21.1 | 49.48146 | 2.151708603 | 0 | 0 |
| 32 | rps14 | 49.47591 | 1.266065836 | 0 | 0 |
| 33 | rps6 | 49.35487 | 1.386384249 | 0 | 0 |
| 34 | rps5 | 49.35282 | 1.321283817 | 0 | 0 |
| 35 | rps15a | 49.28632 | 1.346520066 | 0 | 0 |
| 36 | rps24 | 49.27847 | 1.26134336 | 0 | 0 |
| 37 | rpl19 | 49.21072 | 1.316273451 | 0 | 0 |
| 38 | rpl13 | 49.04952 | 1.298480511 | 0 | 0 |
| 39 | rps23 | 48.86997 | 1.293151021 | 0 | 0 |
| 40 | rpl18a | 48.85421 | 1.300501704 | 0 | 0 |
| 41 | rps4x | 48.74708 | 1.315247774 | 0 | 0 |
| 42 | rps27a | 48.62541 | 1.287004828 | 0 | 0 |
| 43 | rpl17 | 48.2268 | 1.263866067 | 0 | 0 |
| 44 | rps16 | 48.21977 | 1.255934238 | 0 | 0 |
| 45 | rps11 | 48.07288 | 1.259169221 | 0 | 0 |
| 46 | rpl28 | 48.00758 | 1.242783189 | 0 | 0 |
| 47 | rps13 | 47.75944 | 1.283989072 | 0 | 0 |
| 48 | nme2b.1 | 47.68089 | 1.425987959 | 0 | 0 |
| 49 | rpl18 | 47.59703 | 1.262466431 | 0 | 0 |
| 50 | rpl13a | 47.48776 | 1.201495647 | 0 | 0 |
| 51 | rps25 | 47.39968 | 1.329605579 | 0 | 0 |
| 52 | rps3 | 47.33768 | 1.295441389 | 0 | 0 |
| 53 | rps27.1 | 47.14485 | 1.247563481 | 0 | 0 |
| 54 | rpl7 | 47.1441 | 1.279641867 | 0 | 0 |
| 55 | eef1b2 | 47.06527 | 1.361621618 | 0 | 0 |
| 56 | rack1 | 47.02077 | 1.274111271 | 0 | 0 |
| 57 | fabp3 | 46.95059 | 1.62801671 | 0 | 0 |
| 58 | tpt1 | 46.93792 | 1.252314687 | 0 | 0 |
| 59 | eef1a1l1 | 46.3877 | 1.034739852 | 0 | 0 |
| 60 | rps15 | 46.00953 | 1.244218349 | 0 | 0 |
| 61 | rpl5a | 45.88334 | 1.392658472 | 0 | 0 |
| 62 | eef2b | 45.78035 | 1.255832314 | 0 | 0 |
| 63 | si:ch211-288g17.3 | 45.4481 | 1.849825144 | 0 | 0 |
| 64 | rpl36a | 45.25935 | 1.283270121 | 0 | 0 |
| 65 | hmgb1b | 45.25665 | 1.614726186 | 0 | 0 |
| 66 | rpl27 | 45.01738 | 1.249033451 | 0 | 0 |
| 67 | rpl23a | 44.91151 | 1.277434468 | 0 | 0 |
| 68 | rpl24 | 44.5558 | 1.199122429 | 0 | 0 |
| 69 | rpl35 | 44.42942 | 1.212393403 | 0 | 0 |
| 70 | crabp1a | 44.30365 | 1.685112119 | 2.6E-299 | 3.8E-297 |
| 71 | cirbpa | 44.06984 | 1.354929805 | 0 | 9.8E-307 |
| 72 | rpl35a | 43.63777 | 1.238346338 | 0 | 6.4E-308 |
| 73 | rpl39 | 43.45329 | 1.187108159 | 0 | 5.7E-307 |
| 74 | hmga1a | 43.04143 | 1.828811288 | 3E-290 | 4.1E-288 |
| 75 | ppiaa | 42.9143 | 1.288335204 | 1.3E-298 | 1.8E-296 |
| 76 | RPS17 | 42.75465 | 1.163946748 | 3.7E-301 | 5.5E-299 |
| 77 | faua | 42.63739 | 1.165846944 | 2.3E-307 | 3.5E-305 |
| 78 | uba52 | 42.33688 | 1.111465693 | 5.5E-300 | 8E-298 |
| 79 | fabp7a | 42.30542 | 1.598576546 | 2.4E-293 | 3.4E-291 |
| 80 | rpl30 | 41.64565 | 1.158921361 | 4.2E-289 | 5.7E-287 |
| 81 | rpl34 | 41.45354 | 1.149294615 | 4.5E-286 | 6E-284 |
| 82 | rpl31 | 41.24366 | 1.106783748 | 2E-286 | 2.7E-284 |
| 83 | serbp1a | 41.22321 | 1.21775043 | 6.7E-275 | 8.5E-273 |
| 84 | RPL37A | 40.6989 | 1.172426224 | 1.4E-276 | 1.7E-274 |
| 85 | pcna | 40.53197 | 2.519945621 | 1.7E-254 | 2E-252 |
| 86 | cirbpb | 40.42265 | 1.056654692 | 1E-272 | 1.3E-270 |
| 87 | rps26l | 40.22673 | 1.208529234 | 4.5E-272 | 5.6E-270 |
| 88 | h3f3b.1 | 40.20798 | 1.706689119 | 1.8E-266 | 2.2E-264 |
| 89 | seta | 40.18583 | 1.59227562 | 4.8E-257 | 5.7E-255 |
| 90 | si:dkey-238o13.4 | 40.04027 | 3.349406719 | 7.2E-241 | 7.8E-239 |
| 91 | snrpd1 | 40.01326 | 1.582401395 | 2E-257 | 2.3E-255 |
| 92 | naca | 39.92827 | 1.137397766 | 5.1E-267 | 6.2E-265 |
| 93 | snrpb | 39.54791 | 1.488209486 | 5.7E-254 | 6.6E-252 |
| 94 | rpl4 | 39.33717 | 1.294695735 | 1.4E-257 | 1.7E-255 |
| 95 | rpl22l1 | 39.03199 | 1.241069674 | 1.2E-255 | 1.4E-253 |
| 96 | rps17 | 38.826 | 1.253696561 | 6.7E-254 | 7.7E-252 |
| 97 | rpl37-1 | 38.64377 | 1.188984632 | 1.5E-252 | 1.7E-250 |
| 98 | hnrnpabb | 38.45846 | 1.344951391 | 1E-244 | 1.2E-242 |
| 99 | marcksb | 38.33159 | 1.568653822 | 5.9E-241 | 6.3E-239 |

**Cluster 6**

| | names | scores | logfoldchanges | pvals | pvals_adj |
|---|---|---|---|---|---|
| **0** | hmgb2a | 87.25797 | 3.047536612 | 0 | 0 |
| **1** | hmgn2 | 84.35824 | 3.128717184 | 0 | 0 |
| **2** | si:ch211-222l21.1 | 74.9759 | 2.752526045 | 0 | 0 |
| **3** | h3f3b.1 | 71.59824 | 2.711129189 | 0 | 0 |
| **4** | hmgb2b | 68.31348 | 2.523654461 | 0 | 0 |
| **5** | hmga1a | 67.3632 | 2.501763821 | 0 | 0 |
| **6** | tubb2b | 66.21985 | 2.726396322 | 0 | 0 |
| **7** | h2afvb | 55.49695 | 2.118937969 | 0 | 0 |
| **8** | mki67 | 51.94138 | 3.309105635 | 1.9E-281 | 3.3E-279 |
| **9** | si:ch73-281n10.2 | 50.23926 | 2.130485058 | 1.9E-292 | 3.5E-290 |
| **10** | stmn1a | 49.73759 | 2.934290886 | 1.7E-272 | 2.9E-270 |
| **11** | si:ch211-288g17.3 | 47.28036 | 2.094866514 | 6.5E-269 | 1.1E-266 |
| **12** | tuba8l4 | 46.34163 | 1.940417409 | 8.8E-262 | 1.4E-259 |
| **13** | seta | 45.18188 | 1.879339457 | 5E-250 | 7.6E-248 |
| **14** | dek | 45.16863 | 2.623578072 | 1E-240 | 1.5E-238 |
| **15** | hmgb1b | 42.74128 | 1.741088986 | 8.6E-236 | 1.2E-233 |
| **16** | cirbpa | 42.6862 | 1.440792322 | 2.7E-244 | 4E-242 |
| **17** | ran | 42.34833 | 1.5820961 | 3.6E-236 | 5.1E-234 |
| **18** | hmgb1a | 41.80008 | 1.640624285 | 1.1E-225 | 1.5E-223 |
| **19** | ptmab | 40.86359 | 1.430267453 | 1.1E-228 | 1.4E-226 |
| **20** | cbx3a | 40.40691 | 1.952654839 | 1.3E-212 | 1.6E-210 |
| **21** | snrpd1 | 39.98124 | 1.81624794 | 1.8E-212 | 2.2E-210 |
| **22** | hnrnpaba | 39.28574 | 1.372292995 | 4.5E-211 | 5.5E-209 |
| **23** | lbr | 38.72216 | 2.611174583 | 2E-196 | 2.2E-194 |
| **24** | cirbpb | 36.51932 | 1.116986752 | 4.8E-196 | 5.4E-194 |
| **25** | hnrnpabb | 36.44706 | 1.465348244 | 2E-190 | 2.1E-188 |
| **26** | hnrnpa0l-1 | 36.42182 | 1.236985326 | 2E-191 | 2.2E-189 |
| **27** | rpsa | 35.40435 | 1.168693781 | 8.5E-192 | 9.2E-190 |
| **28** | rplp0 | 35.37712 | 1.172370195 | 2.3E-192 | 2.5E-190 |
| **29** | rplp1 | 35.33271 | 1.132003188 | 2.5E-194 | 2.7E-192 |
| **30** | tubb4b | 34.73749 | 1.308233619 | 4.7E-176 | 4.5E-174 |
| **31** | snrpb | 34.64986 | 1.570525169 | 1.4E-175 | 1.3E-173 |
| **32** | stmn1b | 34.09365 | 1.798956037 | 6.7E-175 | 6.3E-173 |
| **33** | syncrip | 34.06445 | 1.401675463 | 1.6E-171 | 1.5E-169 |
| **34** | calm2b | 33.89594 | 1.535692811 | 2.2E-170 | 2E-168 |
| **35** | tuba1a | 33.82048 | 1.875161648 | 1.5E-167 | 1.4E-165 |
| **36** | rps8a | 33.55668 | 1.094730616 | 5.1E-179 | 5.1E-177 |
| **37** | rpl12 | 33.51262 | 1.084750533 | 6.6E-177 | 6.5E-175 |
| **38** | rpl9 | 33.36365 | 1.241927028 | 1E-172 | 9.4E-171 |
| **39** | anp32a | 33.29186 | 1.73392272 | 1.2E-162 | 1E-160 |
| **40** | rps2 | 33.20897 | 1.105596423 | 4.2E-176 | 4.1E-174 |
| **41** | top2a | 33.0755 | 3.056524515 | 2.3E-157 | 1.9E-155 |
| **42** | ccna2 | 33.01615 | 2.935689688 | 7.2E-157 | 5.8E-155 |
| **43** | rps10 | 32.88968 | 1.076672554 | 1.9E-173 | 1.8E-171 |
| **44** | chd7 | 32.51994 | 2.048763275 | 1.6E-155 | 1.2E-153 |
| **45** | rps3a | 32.49846 | 1.088833332 | 8.4E-169 | 7.6E-167 |
| **46** | eef1g | 32.04942 | 1.135146499 | 5.6E-166 | 4.9E-164 |
| **47** | marcksb | 31.92685 | 1.577039838 | 5.8E-156 | 4.6E-154 |
| **48** | rps9 | 31.85204 | 1.059009075 | 2.1E-164 | 1.9E-162 |
| **49** | rpl21 | 31.67633 | 1.063766241 | 2.1E-162 | 1.8E-160 |
| **50** | rps27.1 | 31.66462 | 1.068880439 | 4.8E-162 | 4.1E-160 |
| **51** | ddx39ab | 31.63532 | 1.542527676 | 2.9E-153 | 2.2E-151 |
| **52** | rps5 | 31.55388 | 1.08401835 | 5.1E-162 | 4.3E-160 |
| **53** | cenpf | 31.54819 | 3.224763155 | 7.3E-147 | 5.1E-145 |
| **54** | tuba8l | 31.5033 | 2.619463682 | 1.7E-147 | 1.2E-145 |
| **55** | atrx | 31.48364 | 1.441904068 | 5.2E-152 | 3.9E-150 |
| **56** | rps12 | 31.38468 | 1.05997622 | 3.3E-161 | 2.8E-159 |
| **57** | rpl11 | 31.38432 | 1.067273259 | 2.5E-160 | 2.1E-158 |
| **58** | rpl8 | 31.23812 | 1.011838436 | 3.1E-159 | 2.6E-157 |
| **59** | rack1 | 31.11732 | 1.068050981 | 3.7E-158 | 3E-156 |
| **60** | si:ch211-156b7.4 | 31.02056 | 2.002809048 | 2.6E-147 | 1.9E-145 |
| **61** | khdrbs1a | 30.96933 | 1.123795033 | 1.8E-151 | 1.4E-149 |
| **62** | rpl10 | 30.92304 | 1.046919823 | 5.4E-156 | 4.3E-154 |
| **63** | rpl7a | 30.8417 | 1.033721924 | 7.3E-157 | 5.9E-155 |
| **64** | rplp2l | 30.82251 | 1.149687529 | 9E-156 | 7.1E-154 |
| **65** | rpl17 | 30.79738 | 1.031118155 | 4.6E-156 | 3.7E-154 |
| **66** | rps23 | 30.77365 | 1.030158401 | 9.2E-156 | 7.3E-154 |
| **67** | rps7 | 30.7561 | 1.01433897 | 1.8E-156 | 1.4E-154 |
| **68** | smc1al | 30.75104 | 1.697334647 | 3.1E-145 | 2.2E-143 |
| **69** | rps4x | 30.67778 | 1.067282557 | 2.4E-154 | 1.8E-152 |
| **70** | rpl23 | 30.57838 | 1.031162262 | 2.3E-154 | 1.8E-152 |
| **71** | cdk1 | 30.51452 | 3.077165365 | 3E-140 | 2E-138 |
| **72** | sumo3a | 30.51121 | 1.500948191 | 1.7E-144 | 1.1E-142 |
| **73** | hnrnpa0b | 30.49336 | 1.282516718 | 2.6E-146 | 1.8E-144 |
| **74** | rps3 | 30.45933 | 1.085661173 | 2.8E-151 | 2.1E-149 |
| **75** | rps19 | 30.39997 | 1.08299017 | 2.3E-151 | 1.7E-149 |
| **76** | nusap1 | 30.39967 | 3.574988604 | 8.2E-139 | 5.4E-137 |
| **77** | rps16 | 30.34506 | 0.99266386 | 1.2E-151 | 9.1E-150 |
| **78** | rpl15 | 30.18792 | 1.100134611 | 3E-150 | 2.2E-148 |
| **79** | ranbp1 | 30.18565 | 1.699102163 | 7.9E-142 | 5.3E-140 |
| **80** | rps6 | 30.05888 | 1.113031864 | 6.2E-149 | 4.5E-147 |
| **81** | rps15a | 29.97981 | 1.014885187 | 5.2E-149 | 3.8E-147 |
| **82** | rps27a | 29.95521 | 1.010507107 | 2.2E-148 | 1.6E-146 |
| **83** | rpl7 | 29.87661 | 1.020480514 | 6.6E-148 | 4.7E-146 |
| **84** | rbbp4 | 29.82339 | 1.833594918 | 6.3E-139 | 4.1E-137 |
| **85** | rpl24 | 29.76094 | 0.985218167 | 3.9E-147 | 2.7E-145 |
| **86** | CABZ01058261.1 | 29.62782 | 2.826435566 | 1.2E-134 | 7.6E-133 |
| **87** | sox11a | 29.47841 | 3.320035934 | 7.7E-133 | 4.7E-131 |
| **88** | rpl32 | 29.3923 | 0.991548121 | 3.7E-145 | 2.6E-143 |
| **89** | rpl18 | 29.27316 | 1.010679126 | 5.8E-143 | 3.9E-141 |
| **90** | rpl10a | 29.15111 | 1.003625154 | 4.7E-142 | 3.2E-140 |
| **91** | si:ch211-137a8.4 | 29.10474 | 1.584949732 | 5.7E-134 | 3.6E-132 |
| **92** | rpl3 | 29.00317 | 1.005699158 | 5.4E-141 | 3.6E-139 |
| **93** | hdac1 | 28.81993 | 1.406098843 | 2.8E-133 | 1.7E-131 |
| **94** | snrpf | 28.79664 | 1.455657721 | 2.2E-133 | 1.4E-131 |
| **95** | nono | 28.72711 | 1.620738864 | 4.4E-131 | 2.7E-129 |
| **96** | rps25 | 28.72005 | 1.065409899 | 3.9E-138 | 2.6E-136 |
| **97** | chaf1a | 28.65783 | 2.064607382 | 1.7E-130 | 1E-128 |
| **98** | ube2c | 28.6253 | 3.317968845 | 1.7E-127 | 9.6E-126 |
| **99** | rpl23a | 28.60292 | 1.047563672 | 2.4E-136 | 1.6E-134 |

**Cluster 7**

| | names | scores | logfoldchanges | pvals | pvals_adj |
|---|---|---|---|---|---|
| **0** | si:ch211-222l21.1 | 41.61283 | 1.77926445 | 1.3E-263 | 1.4E-261 |
| **1** | hmgb2b | 38.623 | 1.602672935 | 3.6E-237 | 3.2E-235 |
| **2** | h2afvb | 37.9501 | 1.413876176 | 3.4E-233 | 3.1E-231 |
| **3** | hmgn6 | 35.88954 | 1.159591436 | 2.4E-207 | 1.8E-205 |
| **4** | hnrnpa0l-1 | 35.50745 | 1.024327993 | 8E-205 | 5.8E-203 |
| **5** | rps19 | 34.65981 | 0.914957047 | 1.4E-212 | 1.1E-210 |
| **6** | tubb5 | 34.61428 | 1.913021445 | 3.5E-191 | 2.3E-189 |
| **7** | eef1g | 34.50315 | 0.94448632 | 1.9E-212 | 1.4E-210 |
| **8** | hmgn2 | 34.45097 | 1.665735841 | 5.8E-202 | 4.1E-200 |
| **9** | cirbpb | 33.95919 | 0.873040974 | 2.1E-196 | 1.4E-194 |
| **10** | rps10 | 33.86999 | 0.886373162 | 3.2E-205 | 2.3E-203 |
| **11** | rpl23 | 33.5192 | 0.878149569 | 1.8E-201 | 1.3E-199 |
| **12** | rplp0 | 33.43109 | 0.853233933 | 6.2E-203 | 4.4E-201 |
| **13** | rps12 | 33.38526 | 0.911770105 | 1E-198 | 6.9E-197 |
| **14** | rplp1 | 33.13024 | 0.839943111 | 6.3E-201 | 4.4E-199 |
| **15** | rpl11 | 33.08396 | 0.883604825 | 3.2E-196 | 2.2E-194 |
| **16** | rpl17 | 33.00816 | 0.854162097 | 6.5E-197 | 4.4E-195 |
| **17** | fabp7a | 32.99862 | 1.321530581 | 8.2E-185 | 5.2E-183 |
| **18** | rps9 | 32.12623 | 0.828430891 | 1.5E-188 | 1E-186 |
| **19** | rps5 | 31.86653 | 0.84949702 | 6.4E-186 | 4.1E-184 |
| **20** | rps3a | 31.68187 | 0.832876205 | 7E-183 | 4.4E-181 |
| **21** | rps27.1 | 31.66276 | 0.856041193 | 4.5E-181 | 2.8E-179 |
| **22** | eef1b2 | 31.55722 | 0.932367086 | 1.9E-175 | 1.1E-173 |
| **23** | rpl15 | 31.44047 | 0.882215798 | 4.5E-180 | 2.8E-178 |
| **24** | rpl21 | 31.41321 | 0.838808417 | 1.9E-179 | 1.2E-177 |
| **25** | rps13 | 31.09816 | 0.845565557 | 5.4E-175 | 3.2E-173 |
| **26** | rpl7a | 30.99184 | 0.810452461 | 1.3E-177 | 7.9E-176 |
| **27** | rpl8 | 30.97138 | 0.794834077 | 1.3E-175 | 7.6E-174 |
| **28** | rplp2l | 30.96316 | 0.920136034 | 4.6E-175 | 2.7E-173 |
| **29** | rps7 | 30.83394 | 0.7879498 | 7E-177 | 4.2E-175 |
| **30** | rpl13 | 30.68253 | 0.80433017 | 5.6E-173 | 3.3E-171 |
| **31** | rpl12 | 30.45439 | 0.809612393 | 1.9E-169 | 1.1E-167 |
| **32** | rpl28 | 30.41 | 0.81254369 | 3.6E-169 | 2.1E-167 |
| **33** | rps4x | 30.4074 | 0.82513994 | 2.4E-170 | 1.4E-168 |
| **34** | rpl10 | 30.38111 | 0.792707264 | 2.6E-170 | 1.5E-168 |
| **35** | rps15a | 30.33371 | 0.827185452 | 4.6E-168 | 2.7E-166 |
| **36** | rpsa | 30.12223 | 0.801590681 | 1.1E-167 | 6.3E-166 |
| **37** | rpl10a | 29.91078 | 0.77975297 | 1E-165 | 5.7E-164 |
| **38** | si:dkey-151g10.6 | 29.90657 | 0.76859659 | 7E-165 | 3.9E-163 |
| **39** | rpl32 | 29.80276 | 0.828407943 | 5.8E-163 | 3.2E-161 |
| **40** | rps8a | 29.78196 | 0.771633089 | 1.3E-166 | 7.6E-165 |
| **41** | rpl24 | 29.74594 | 0.792007267 | 2.1E-162 | 1.2E-160 |
| **42** | rpl18 | 29.6037 | 0.791923046 | 6E-162 | 3.3E-160 |
| **43** | rps25 | 29.56172 | 0.856716514 | 4.1E-160 | 2.2E-158 |
| **44** | rpl3 | 29.30279 | 0.778002977 | 9.8E-160 | 5.3E-158 |
| **45** | rpl35a | 29.29649 | 0.839905143 | 5.6E-156 | 2.9E-154 |
| **46** | rps16 | 29.25809 | 0.781751037 | 1.4E-157 | 7.2E-156 |
| **47** | fabp3 | 29.08586 | 1.149132252 | 5.5E-149 | 2.8E-147 |
| **48** | rps24 | 28.81985 | 0.755632758 | 8.4E-155 | 4.3E-153 |
| **49** | stmn1b | 28.65819 | 1.455160141 | 2.9E-144 | 1.4E-142 |
| **50** | faua | 28.60137 | 0.82081449 | 6.2E-151 | 3.1E-149 |
| **51** | si:ch211-288g17.3 | 28.44066 | 1.324968338 | 6.4E-142 | 3E-140 |
| **52** | rps14 | 28.36089 | 0.728639066 | 1.3E-151 | 6.6E-150 |
| **53** | rps6 | 28.25324 | 0.825858593 | 2.3E-149 | 1.1E-147 |
| **54** | rps27a | 28.15806 | 0.751810312 | 1.6E-148 | 7.8E-147 |
| **55** | sox11b | 27.82893 | 2.157065868 | 1.1E-131 | 4.9E-130 |
| **56** | rpl7 | 27.66368 | 0.753323257 | 4.8E-144 | 2.3E-142 |
| **57** | rps15 | 27.65852 | 0.770883679 | 1.3E-142 | 5.9E-141 |
| **58** | rps2 | 27.62746 | 0.743544221 | 9.6E-146 | 4.6E-144 |
| **59** | rack1 | 27.57765 | 0.769946098 | 1.3E-143 | 6.3E-142 |
| **60** | rpl19 | 27.32542 | 0.712888956 | 1.3E-143 | 6.3E-142 |
| **61** | ppiaa | 27.16442 | 0.858769953 | 1.7E-135 | 7.6E-134 |
| **62** | rps11 | 27.00904 | 0.743216157 | 5.1E-137 | 2.3E-135 |
| **63** | rpl18a | 26.99198 | 0.726323903 | 7.4E-139 | 3.4E-137 |
| **64** | rps3 | 26.98615 | 0.789535046 | 2.1E-136 | 9.2E-135 |
| **65** | hnrnpabb | 26.85308 | 1.010205626 | 8E-130 | 3.4E-128 |
| **66** | rps23 | 26.71899 | 0.722133517 | 1.8E-136 | 8E-135 |
| **67** | cirbpa | 26.68651 | 0.857136786 | 2.1E-131 | 8.9E-130 |
| **68** | rpl27 | 26.49814 | 0.747077525 | 1.1E-132 | 5E-131 |
| **69** | rpl23a | 26.48748 | 0.792256594 | 5.3E-131 | 2.3E-129 |
| **70** | tp53inp2 | 26.20919 | 1.926505923 | 2.7E-120 | 1.1E-118 |
| **71** | khdrbs1a | 26.02629 | 0.786468565 | 3.6E-125 | 1.5E-123 |
| **72** | rpl39 | 25.90498 | 0.765111566 | 1.2E-125 | 4.9E-124 |
| **73** | cnp | 25.84046 | 1.600690603 | 9.2E-119 | 3.6E-117 |
| **74** | rpl9 | 25.76854 | 0.810307562 | 1.7E-125 | 7E-124 |
| **75** | rpl13a | 25.6954 | 0.66696465 | 2.8E-126 | 1.2E-124 |
| **76** | rps26l | 25.57702 | 0.781823635 | 1.2E-123 | 4.8E-122 |
| **77** | ptmab | 25.47652 | 0.821610272 | 1.6E-121 | 6.5E-120 |
| **78** | hnrnpaba | 25.3942 | 0.783416629 | 1.2E-119 | 4.6E-118 |
| **79** | rpl36a | 24.54305 | 0.736450851 | 2.2E-115 | 8.4E-114 |
| **80** | marcksb | 24.38404 | 1.152520537 | 1.8E-109 | 6.5E-108 |
| **81** | rpl5a | 24.31759 | 0.778606951 | 3.3E-113 | 1.2E-111 |
| **82** | snrpb | 24.28328 | 1.00574255 | 1.3E-109 | 4.6E-108 |
| **83** | rps17 | 24.15112 | 0.820433795 | 9.7E-111 | 3.6E-109 |
| **84** | rpl35 | 23.73368 | 0.702388763 | 2.9E-108 | 1E-106 |
| **85** | naca | 23.6569 | 0.694908679 | 1.4E-107 | 5E-106 |
| **86** | rpl30 | 23.61177 | 0.687989235 | 8.5E-108 | 3E-106 |
| **87** | h3f3b.1 | 23.56041 | 0.950644314 | 1.3E-107 | 4.7E-106 |
| **88** | rpl31 | 23.53444 | 0.673663616 | 3.9E-107 | 1.4E-105 |
| **89** | hdac1 | 23.37359 | 1.009631634 | 4.8E-102 | 1.6E-100 |
| **90** | ddx39ab | 23.33385 | 1.070853829 | 1.2E-101 | 3.8E-100 |
| **91** | RPS17 | 23.28409 | 0.667254567 | 3.4E-105 | 1.2E-103 |
| **92** | rpl22 | 23.16479 | 0.731656432 | 7.7E-103 | 2.6E-101 |
| **93** | uba52 | 22.68652 | 0.617799282 | 3.1E-101 | 1E-99 |
| **94** | snrpd1 | 22.41313 | 0.968627453 | 6.54E-96 | 2.04E-94 |
| **95** | rpl34 | 22.39778 | 0.673187137 | 8.59E-98 | 2.75E-96 |
| **96** | serbp1a | 22.22529 | 0.698510051 | 1.76E-95 | 5.43E-94 |
| **97** | si:dkey-56m19.5 | 22.13316 | 1.587211847 | 1.87E-91 | 5.57E-90 |
| **98** | rpl14 | 22.03477 | 0.628465295 | 2.63E-95 | 8.12E-94 |
| **99** | insm1a | 21.95723 | 2.050113201 | 5.94E-90 | 1.75E-88 |

**Cluster 8**

| | names | scores | logfoldchanges | pvals | pvals_adj |
|---|---|---|---|---|---|
| **0** | hmgb2a | 67.12348 | 2.779353619 | 0 | 0 |
| **1** | neurod1 | 59.31221 | 4.457812786 | 0 | 0 |
| **2** | hmgn2 | 56.68037 | 2.804169416 | 0 | 0 |
| **3** | hmga1a | 54.15988 | 2.229558468 | 0 | 0 |
| **4** | otx5 | 53.13665 | 4.293460846 | 3.3E-302 | 4.8E-300 |
| **5** | h3f3b.1 | 52.59372 | 2.427684546 | 0 | 0 |
| **6** | stmn1a | 49.46758 | 3.096511364 | 3.6E-289 | 4.9E-287 |
| **7** | si:ch73-281n10.2 | 49.11773 | 2.244504929 | 6.4E-300 | 9.3E-298 |
| **8** | hmgb2b | 48.6291 | 2.011791945 | 0 | 0 |
| **9** | nr2e3 | 47.91886 | 3.956299543 | 8.9E-270 | 1.1E-267 |
| **10** | pde6gb | 44.3974 | 4.142101765 | 3.1E-247 | 3.4E-245 |
| **11** | h2afvb | 43.6097 | 1.809593678 | 3.1E-269 | 3.8E-267 |
| **12** | h2afx | 40.63955 | 3.02434516 | 4.5E-221 | 4.2E-219 |
| **13** | tuba8l4 | 40.17926 | 1.770045161 | 3.3E-231 | 3.3E-229 |
| **14** | tubb2b | 39.5692 | 2.097799063 | 2.1E-225 | 2E-223 |
| **15** | hmgn6 | 38.96754 | 1.345105648 | 1.7E-226 | 1.6E-224 |
| **16** | stmn1b | 38.70943 | 2.050158739 | 7.3E-220 | 6.8E-218 |
| **17** | faua | 38.42354 | 1.195759892 | 3.8E-231 | 3.7E-229 |
| **18** | arl13a | 38.16461 | 3.777628422 | 6.7E-202 | 5.6E-200 |
| **19** | chaf1a | 37.5683 | 2.378391981 | 2E-203 | 1.7E-201 |
| **20** | crx | 37.20008 | 3.141419172 | 1E-196 | 8E-195 |
| **21** | si:ch211-156b7.4 | 37.1531 | 2.186157703 | 7.2E-202 | 6E-200 |
| **22** | rps27.1 | 36.82118 | 1.130154133 | 6.2E-218 | 5.7E-216 |
| **23** | si:ch211-288g17.3 | 36.57405 | 1.676701427 | 6.3E-205 | 5.3E-203 |
| **24** | anp32e | 35.88087 | 1.735583663 | 2.4E-193 | 1.9E-191 |
| **25** | inhbb | 35.48219 | 3.08743906 | 5.5E-183 | 4E-181 |
| **26** | ptmab | 35.37823 | 1.268137455 | 2.1E-197 | 1.7E-195 |
| **27** | dek | 35.36619 | 2.233664989 | 6.6E-187 | 4.9E-185 |
| **28** | rplp1 | 35.1895 | 1.066068888 | 2.8E-206 | 2.4E-204 |
| **29** | h2afva | 35.14886 | 1.827295303 | 1.3E-185 | 9.5E-184 |
| **30** | cirbpa | 34.75042 | 1.185677767 | 1.9E-194 | 1.5E-192 |
| **31** | si:ch73-28h20.1 | 34.5794 | 4.081810474 | 4.9E-176 | 3.3E-174 |
| **32** | rbbp4 | 34.35822 | 1.939522028 | 1.9E-180 | 1.4E-178 |
| **33** | rps9 | 34.31971 | 1.029520035 | 6.3E-198 | 5.1E-196 |
| **34** | rps5 | 34.1325 | 1.087626338 | 1.3E-194 | 1.1E-192 |
| **35** | rplp2l | 34.03513 | 1.176220059 | 7.4E-193 | 5.8E-191 |
| **36** | rpl21 | 33.5185 | 1.059054494 | 7.4E-188 | 5.6E-186 |
| **37** | rpsa | 33.44976 | 1.076921821 | 1.7E-186 | 1.3E-184 |
| **38** | rps10 | 33.04453 | 1.010643363 | 3.6E-186 | 2.7E-184 |
| **39** | rpl10 | 32.74474 | 1.043875098 | 2.5E-180 | 1.8E-178 |
| **40** | cxxc5a | 32.58 | 2.115901709 | 6.7E-165 | 4.3E-163 |
| **41** | rpl7a | 32.43849 | 1.0223912 | 1.2E-179 | 8.6E-178 |
| **42** | rpl11 | 32.37378 | 1.027661562 | 1.3E-178 | 9.2E-177 |
| **43** | rps23 | 32.34606 | 1.042908907 | 4.2E-177 | 2.9E-175 |
| **44** | rack1 | 32.30142 | 1.076356053 | 3.7E-176 | 2.6E-174 |
| **45** | insm1a | 32.22336 | 2.895099401 | 3.8E-161 | 2.3E-159 |
| **46** | rps7 | 32.1926 | 1.021976233 | 8E-177 | 5.6E-175 |
| **47** | rps3 | 32.12946 | 1.044309974 | 6.3E-175 | 4.3E-173 |
| **48** | atp1a3b | 31.93337 | 3.332612753 | 2.3E-157 | 1.3E-155 |
| **49** | rps19 | 31.84344 | 1.010191679 | 3.3E-174 | 2.2E-172 |
| **50** | rpl18a | 31.7183 | 1.017709851 | 7.5E-172 | 5E-170 |
| **51** | rps4x | 31.4954 | 1.038496614 | 8.1E-170 | 5.4E-168 |
| **52** | ran | 31.45869 | 1.139656544 | 2.7E-166 | 1.8E-164 |
| **53** | rps3a | 31.36823 | 0.995694935 | 2E-169 | 1.3E-167 |
| **54** | rps25 | 31.31455 | 1.067583323 | 2.5E-167 | 1.6E-165 |
| **55** | rpl28 | 31.12911 | 0.963128924 | 1.2E-167 | 7.8E-166 |
| **56** | msi1 | 30.95373 | 1.650294542 | 1.3E-153 | 7.4E-152 |
| **57** | rpl8 | 30.82375 | 0.984676838 | 2.2E-163 | 1.4E-161 |
| **58** | six7 | 30.60105 | 3.94678998 | 2.5E-147 | 1.3E-145 |
| **59** | lbr | 30.58515 | 2.219436884 | 2.5E-150 | 1.4E-148 |
| **60** | rpl24 | 30.45271 | 0.952076495 | 7.2E-161 | 4.4E-159 |
| **61** | rpl15 | 30.40518 | 1.067016363 | 9.8E-160 | 5.9E-158 |
| **62** | rps14 | 30.34067 | 0.921545327 | 3.2E-161 | 2E-159 |
| **63** | rpl23 | 30.28661 | 0.97721374 | 3.8E-160 | 2.3E-158 |
| **64** | rpl18 | 30.26096 | 0.981683016 | 7.9E-159 | 4.7E-157 |
| **65** | rps16 | 30.24792 | 0.96431154 | 2.9E-158 | 1.7E-156 |
| **66** | pcna | 30.1896 | 2.281074047 | 1.5E-148 | 8.5E-147 |
| **67** | rps6 | 30.10794 | 1.080718756 | 9.9E-157 | 5.8E-155 |
| **68** | khdrbs1a | 30.10605 | 1.004435897 | 4.6E-154 | 2.7E-152 |
| **69** | rps12 | 30.09531 | 1.021129608 | 1.6E-157 | 9.5E-156 |
| **70** | rps8a | 29.98163 | 0.976803482 | 1.6E-157 | 9.5E-156 |
| **71** | rrm2-1 | 29.90607 | 2.406019211 | 1.4E-145 | 7.5E-144 |
| **72** | rpl10a | 29.89342 | 0.939995348 | 1E-156 | 6E-155 |
| **73** | rps24 | 29.74668 | 0.938270032 | 8E-155 | 4.6E-153 |
| **74** | rbp4l | 29.58597 | 3.06001687 | 2.1E-142 | 1.1E-140 |
| **75** | rps13 | 29.55441 | 0.949156225 | 1.9E-153 | 1.1E-151 |
| **76** | snrpd1 | 29.49668 | 1.342951179 | 5.4E-147 | 2.9E-145 |
| **77** | thrb | 29.49044 | 3.617175579 | 1.4E-139 | 7.1E-138 |
| **78** | rpl17 | 29.48057 | 0.951195896 | 3.1E-153 | 1.8E-151 |
| **79** | rpl13 | 29.37915 | 0.956986487 | 1.4E-151 | 8.1E-150 |
| **80** | rps15 | 29.22549 | 0.958225787 | 1.5E-149 | 8.1E-148 |
| **81** | rps27a | 28.99521 | 0.94915694 | 7.8E-148 | 4.3E-146 |
| **82** | snrpf | 28.95937 | 1.355659008 | 5.9E-142 | 3.1E-140 |
| **83** | rpl3 | 28.85182 | 0.954410315 | 7.3E-147 | 4E-145 |
| **84** | rpl19 | 28.73604 | 0.925646305 | 1.4E-147 | 7.9E-146 |
| **85** | rpl7 | 28.69171 | 0.960864484 | 3.3E-145 | 1.8E-143 |
| **86** | seta | 28.57745 | 1.258389473 | 4.5E-140 | 2.3E-138 |
| **87** | rps15a | 28.40662 | 0.911839604 | 3.8E-144 | 2E-142 |
| **88** | uba52 | 28.37418 | 0.907005012 | 3.3E-142 | 1.7E-140 |
| **89** | rpl32 | 28.36332 | 0.912043869 | 3.9E-144 | 2.1E-142 |
| **90** | mki67 | 28.3106 | 2.186181068 | 1.3E-134 | 6.2E-133 |
| **91** | rpl30 | 28.28944 | 0.941928029 | 3.8E-141 | 2E-139 |
| **92** | ppiab | 28.28312 | 0.947445273 | 1.4E-137 | 6.8E-136 |
| **93** | si:dkey-151g10.6 | 28.18334 | 0.889485359 | 1.2E-141 | 6.3E-140 |
| **94** | rps2 | 28.0612 | 0.917723715 | 5.3E-142 | 2.8E-140 |
| **95** | eef1g | 27.74969 | 0.959605694 | 5.8E-139 | 3E-137 |
| **96** | slbp | 27.74192 | 2.341868877 | 4.2E-129 | 2E-127 |
| **97** | cct2-1 | 27.73662 | 1.251080871 | 1.7E-132 | 8.2E-131 |
| **98** | mibp | 27.68613 | 2.588896036 | 2.6E-128 | 1.2E-126 |
| **99** | rxrgb | 27.67214 | 3.213609934 | 3.3E-127 | 1.5E-125 |

**Cluster 9**

| | names | scores | logfoldchanges | pvals | pvals_adj |
|---|---|---|---|---|---|
| 0 | pde6gb | 45.31855 | 5.805340767 | 4.6E-136 | 3.8E-134 |
| 1 | h3f3b.1 | 34.27803 | 2.064031601 | 3.8E-114 | 2.5E-112 |
| 2 | tmsb4x | 34.15756 | 1.992908597 | 2.1E-108 | 1.3E-106 |
| 3 | hmgb1b | 28.27168 | 1.809103131 | 2.27E-89 | 1.15E-87 |
| 4 | h2afvb | 27.3001 | 1.631613851 | 1.7E-88 | 8.48E-87 |
| 5 | hmgn2 | 26.91901 | 2.163398504 | 8.13E-87 | 4.01E-85 |
| 6 | prox1a | 26.91223 | 3.606124163 | 1.21E-81 | 5.59E-80 |
| 7 | tubb5 | 24.92814 | 2.35922718 | 1.62E-76 | 7.03E-75 |
| 8 | zeb2a | 22.976 | 2.637861729 | 4.13E-68 | 1.59E-66 |
| 9 | tubb2b | 22.90328 | 1.908594608 | 1.59E-69 | 6.28E-68 |
| 10 | si:ch211-288g17.3 | 22.47307 | 1.657690525 | 4.45E-68 | 1.71E-66 |
| 11 | h3f3d | 22.46573 | 0.92917341 | 7.87E-69 | 3.07E-67 |
| 12 | cotl1 | 22.42771 | 1.616794944 | 3.92E-67 | 1.49E-65 |
| 13 | ywhah | 22.39713 | 2.090929985 | 1.88E-66 | 7.14E-65 |
| 14 | hmgn6 | 22.19003 | 1.086266279 | 3.11E-68 | 1.2E-66 |
| 15 | rem1 | 21.49858 | 8.35446167 | 3.98E-62 | 1.39E-60 |
| 16 | onecut1 | 21.43028 | 5.181397915 | 5.7E-62 | 1.97E-60 |
| 17 | tfap2b | 21.32709 | 5.2685256 | 1.28E-61 | 4.41E-60 |
| 18 | hmgb2b | 21.25424 | 1.528167725 | 3.17E-64 | 1.15E-62 |
| 19 | stmn1a | 21.21375 | 2.578372955 | 5.9E-62 | 2.04E-60 |
| 20 | ptmab | 21.08867 | 1.104574323 | 1.2E-63 | 4.29E-62 |
| 21 | fabp3 | 21.0306 | 1.377563715 | 8.75E-63 | 3.08E-61 |
| 22 | eef1g | 20.96286 | 0.913054764 | 2.32E-65 | 8.59E-64 |
| 23 | rps27.1 | 20.50296 | 0.867426991 | 5.66E-63 | 2E-61 |
| 24 | onecutl | 20.34417 | 7.409417629 | 6.3E-58 | 2.05E-56 |
| 25 | tox | 20.27131 | 2.241892576 | 3.25E-58 | 1.06E-56 |
| 26 | golga7ba | 20.17522 | 2.910443783 | 1.11E-57 | 3.6E-56 |
| 27 | CR361564.1 | 19.95805 | 7.87608242 | 1.67E-56 | 5.28E-55 |
| 28 | ndrg4 | 19.79968 | 2.136945248 | 1.27E-56 | 4.05E-55 |
| 29 | stmn1b | 19.54752 | 1.511133909 | 3.98E-57 | 1.27E-55 |
| 30 | rbfox2 | 19.02843 | 3.179932833 | 2.18E-53 | 6.5E-52 |
| 31 | rpl28 | 18.7783 | 0.775095165 | 7.9E-56 | 2.47E-54 |
| 32 | rps10 | 18.54901 | 0.7647475 | 3.83E-55 | 1.19E-53 |
| 33 | rps9 | 18.3002 | 0.718524575 | 2.08E-54 | 6.31E-53 |
| 34 | rbbp4 | 18.2825 | 1.732675076 | 3.94E-51 | 1.13E-49 |
| 35 | rps5 | 18.07771 | 0.736728191 | 2.07E-53 | 6.19E-52 |
| 36 | rps19 | 17.97738 | 0.791886091 | 2.75E-52 | 8.04E-51 |
| 37 | hnrnpaba | 17.93326 | 0.935697317 | 1.22E-50 | 3.47E-49 |
| 38 | rps12 | 17.93323 | 0.755777478 | 1.66E-52 | 4.9E-51 |
| 39 | rpl10 | 17.88057 | 0.713867128 | 2.38E-52 | 6.95E-51 |
| 40 | rps4x | 17.8334 | 0.733098507 | 3.43E-52 | 9.97E-51 |
| 41 | nme2b.1 | 17.78885 | 0.871106267 | 1E-50 | 2.85E-49 |
| 42 | seta | 17.73796 | 1.204390645 | 1.32E-49 | 3.63E-48 |
| 43 | faua | 17.69118 | 0.793981969 | 6.84E-51 | 1.95E-49 |
| 44 | hmgn7 | 17.56817 | 1.301896214 | 1.67E-48 | 4.46E-47 |
| 45 | rpl7a | 17.42317 | 0.69759804 | 1.26E-50 | 3.58E-49 |
| 46 | si:ch73-1a9.3 | 17.15257 | 1.245741606 | 4.12E-47 | 1.08E-45 |
| 47 | si:ch211-222l21.1 | 17.11261 | 1.312135935 | 1.24E-47 | 3.27E-46 |
| 48 | atrx | 17.06371 | 1.236733079 | 1.17E-46 | 3E-45 |
| 49 | si:ch73-281n10.2 | 17.04289 | 1.513535976 | 1.42E-46 | 3.62E-45 |
| 50 | rpl32 | 16.88686 | 0.716740131 | 6.63E-48 | 1.77E-46 |
| 51 | sumo3a | 16.8608 | 1.308552742 | 8.66E-46 | 2.17E-44 |
| 52 | rpl23 | 16.6083 | 0.672060072 | 4.62E-47 | 1.2E-45 |
| 53 | rplp1 | 16.46432 | 0.633801401 | 7.25E-47 | 1.88E-45 |
| 54 | rack1 | 16.45527 | 0.720064938 | 4.86E-46 | 1.23E-44 |
| 55 | six6b | 15.97176 | 2.492325068 | 6.55E-42 | 1.49E-40 |
| 56 | rpl24 | 15.96949 | 0.700263202 | 9.51E-44 | 2.28E-42 |
| 57 | khdrbs1a | 15.95052 | 0.823039651 | 6.83E-43 | 1.6E-41 |
| 58 | rpl21 | 15.85818 | 0.657526731 | 9.88E-44 | 2.37E-42 |
| 59 | chd4a | 15.79557 | 1.609508276 | 1.7E-41 | 3.85E-40 |
| 60 | hnrnpa0l-1 | 15.71274 | 0.720775008 | 3.24E-42 | 7.49E-41 |
| 61 | rps3a | 15.69729 | 0.670445561 | 6.29E-43 | 1.48E-41 |
| 62 | hmgb2a | 15.68408 | 1.461215496 | 1.37E-41 | 3.11E-40 |
| 63 | rpsa | 15.63609 | 0.666253924 | 1.08E-42 | 2.52E-41 |
| 64 | rpl17 | 15.56202 | 0.653773844 | 1.8E-42 | 4.17E-41 |
| 65 | rplp2l | 15.5487 | 0.759736955 | 3.51E-42 | 8.07E-41 |
| 66 | rpl19 | 15.45128 | 0.632404208 | 3.3E-42 | 7.62E-41 |
| 67 | plekhg4 | 15.42409 | 2.689051151 | 8.05E-40 | 1.75E-38 |
| 68 | cirbpa | 15.32252 | 0.796740949 | 1.09E-40 | 2.42E-39 |
| 69 | h3f3b.1-1 | 15.31708 | 1.084022284 | 5.07E-40 | 1.11E-38 |
| 70 | si:dkey-28b4.7 | 15.11673 | 1.687421083 | 7.11E-39 | 1.5E-37 |
| 71 | rpl18 | 15.04653 | 0.639845014 | 3.16E-40 | 6.96E-39 |
| 72 | h2afx | 15.01055 | 2.10858798 | 2.07E-38 | 4.32E-37 |
| 73 | fkbp1aa | 14.93795 | 1.043140769 | 1.69E-38 | 3.54E-37 |
| 74 | ndufa4l | 14.89999 | 0.926854014 | 1.3E-38 | 2.74E-37 |
| 75 | rpl11 | 14.88396 | 0.668128371 | 1.79E-39 | 3.85E-38 |
| 76 | tmsb | 14.86012 | 1.917511344 | 6.1E-38 | 1.26E-36 |
| 77 | zc4h2 | 14.84985 | 2.016158581 | 8.18E-38 | 1.69E-36 |
| 78 | rps3 | 14.83941 | 0.708161175 | 4.79E-39 | 1.02E-37 |
| 79 | rpl13 | 14.83679 | 0.597650826 | 1.31E-39 | 2.83E-38 |
| 80 | rpl8 | 14.60755 | 0.605774581 | 1.81E-38 | 3.79E-37 |
| 81 | rps8a | 14.55141 | 0.602845669 | 1.89E-38 | 3.96E-37 |
| 82 | rps7 | 14.49462 | 0.593027949 | 3.11E-38 | 6.49E-37 |
| 83 | rpl3 | 14.48732 | 0.63044709 | 6.71E-38 | 1.39E-36 |
| 84 | pclaf | 14.42333 | 2.173114538 | 3.3E-36 | 6.55E-35 |
| 85 | tmeff1b | 14.40351 | 1.49878943 | 3.16E-36 | 6.28E-35 |
| 86 | rps27a | 14.25091 | 0.582784176 | 4.14E-37 | 8.42E-36 |
| 87 | mab21l1 | 14.21964 | 1.896145344 | 1.86E-35 | 3.62E-34 |
| 88 | uba52 | 14.16737 | 0.620055854 | 2.07E-36 | 4.15E-35 |
| 89 | si:ch211-156b7.4 | 14.12852 | 1.598426938 | 2.92E-35 | 5.66E-34 |
| 90 | rps6 | 14.1061 | 0.690129936 | 3.05E-36 | 6.07E-35 |
| 91 | lsm6 | 14.0396 | 1.410925865 | 7.21E-35 | 1.38E-33 |
| 92 | rps14 | 14.0236 | 0.589617789 | 4.36E-36 | 8.62E-35 |
| 93 | pcdh8 | 14.00961 | 4.083306789 | 1.56E-34 | 2.96E-33 |
| 94 | rps25 | 13.90268 | 0.687786162 | 2.52E-35 | 4.89E-34 |
| 95 | chaf1a | 13.8414 | 1.687627554 | 3.73E-34 | 7.02E-33 |
| 96 | rpl10a | 13.80906 | 0.583319247 | 2.93E-35 | 5.66E-34 |
| 97 | rrm2-1 | 13.73925 | 2.040427446 | 1.03E-33 | 1.92E-32 |
| 98 | dalrd3 | 13.65998 | 3.842134714 | 3.21E-33 | 5.91E-32 |
| 99 | rps13 | 13.61701 | 0.589052677 | 1.87E-34 | 3.54E-33 |

**Cluster 10**

| | names | scores | logfoldchanges | pvals | pvals_adj |
|---|---|---|---|---|---|
| **0** | elavl3 | 77.99724 | 6.284736633 | 1.4E-219 | 2.5E-217 |
| **1** | tmsb | 69.54066 | 4.788538456 | 7.9E-207 | 1.3E-204 |
| **2** | tubb5 | 69.38703 | 5.394479275 | 1.3E-205 | 2.2E-203 |
| **3** | stmn1b | 66.6868 | 3.850678444 | 1.9E-213 | 3.3E-211 |
| **4** | gap43 | 63.14544 | 8.218997002 | 1.7E-186 | 2.4E-184 |
| **5** | tuba1c | 54.22678 | 4.22229147 | 4E-171 | 4.9E-169 |
| **6** | mllt11 | 53.4115 | 4.735610962 | 1E-166 | 1.2E-164 |
| **7** | stmn2b | 52.82681 | 7.843675613 | 4.2E-163 | 4.9E-161 |
| **8** | vim | 52.47554 | 4.247157097 | 1.6E-166 | 1.9E-164 |
| **9** | tmsb4x | 47.32957 | 2.277805805 | 2.4E-159 | 2.6E-157 |
| **10** | marcksl1b | 44.28115 | 2.224685192 | 2.5E-157 | 2.7E-155 |
| **11** | rbpms2b | 42.33289 | 3.402827024 | 2.5E-139 | 2.3E-137 |
| **12** | tuba1a | 41.57 | 2.990813017 | 1.8E-138 | 1.6E-136 |
| **13** | uchl1 | 41.34855 | 4.457408428 | 6.6E-135 | 5.8E-133 |
| **14** | fabp3 | 41.18753 | 2.427572489 | 9.7E-141 | 9E-139 |
| **15** | gng3 | 39.73628 | 4.544009209 | 4.7E-129 | 3.8E-127 |
| **16** | rtn1a | 39.29915 | 3.117470741 | 3.3E-129 | 2.7E-127 |
| **17** | tmsb2 | 38.12695 | 5.78056097 | 2.6E-123 | 2E-121 |
| **18** | cnp | 36.78994 | 3.632299185 | 2.2E-120 | 1.7E-118 |
| **19** | rtn1b | 36.76604 | 4.318268776 | 1.4E-119 | 1E-117 |
| **20** | stmn2a | 35.2411 | 7.416443825 | 7.7E-114 | 5.3E-112 |
| **21** | dpysl3 | 35.18645 | 4.678085804 | 3.3E-114 | 2.3E-112 |
| **22** | klf7b | 34.3878 | 2.974934101 | 2.6E-113 | 1.8E-111 |
| **23** | vdac3 | 34.21999 | 2.901899815 | 2.2E-112 | 1.5E-110 |
| **24** | si:dkey-280e21.3 | 33.88601 | 5.158335209 | 8.3E-110 | 5.4E-108 |
| **25** | alcama | 33.70361 | 4.884744167 | 3.2E-109 | 2.1E-107 |
| **26** | nova2 | 33.13497 | 2.707598686 | 1.1E-109 | 7E-108 |
| **27** | ywhaz | 33.05883 | 3.038551569 | 2.9E-108 | 1.9E-106 |
| **28** | rbfox2 | 33.05655 | 4.428223133 | 2.2E-107 | 1.4E-105 |
| **29** | inab | 32.79713 | 7.990090847 | 9E-106 | 5.6E-104 |
| **30** | jpt1b | 32.71454 | 2.790271997 | 2.5E-107 | 1.5E-105 |
| **31** | elavl4 | 32.17347 | 7.083750248 | 9.9E-104 | 5.9E-102 |
| **32** | islr2 | 32.03269 | 6.528103828 | 2.8E-103 | 1.7E-101 |
| **33** | map1b | 31.80873 | 4.402173519 | 4.5E-103 | 2.6E-101 |
| **34** | si:dkey-276j7.1 | 31.67456 | 3.559438229 | 7.8E-103 | 4.6E-101 |
| **35** | vat1 | 31.22507 | 2.382035255 | 4.2E-104 | 2.5E-102 |
| **36** | stmn4l | 30.59625 | 7.518033981 | 2.59E-98 | 1.45E-96 |
| **37** | si:dkey-56m19.5 | 29.84246 | 2.891279936 | 2.82E-97 | 1.56E-95 |
| **38** | rab6bb | 29.55522 | 4.517085552 | 5.59E-95 | 3E-93 |
| **39** | gdi1 | 29.28638 | 3.562043667 | 2.12E-94 | 1.13E-92 |
| **40** | stmn4 | 29.16547 | 8.488958359 | 2.93E-93 | 1.54E-91 |
| **41** | tubb4b | 29.07067 | 1.347495914 | 1.99E-98 | 1.12E-96 |
| **42** | tubb2 | 28.98465 | 7.388160229 | 1.21E-92 | 6.32E-91 |
| **43** | cfl1 | 28.3221 | 1.294718027 | 4.94E-94 | 2.62E-92 |
| **44** | tuba2 | 27.83016 | 5.600058079 | 1.23E-88 | 6.08E-87 |
| **45** | anxa13l | 27.74301 | 6.787528992 | 3.2E-88 | 1.57E-86 |
| **46** | scrt2 | 27.62558 | 4.455796242 | 3.19E-88 | 1.57E-86 |
| **47** | maptb | 27.5245 | 4.891450405 | 1.1E-87 | 5.37E-86 |
| **48** | gpm6aa | 27.41314 | 1.519470811 | 3.64E-91 | 1.86E-89 |
| **49** | tubb2b | 27.31199 | 1.847708821 | 7.06E-92 | 3.64E-90 |
| **50** | tmeff1b | 26.70131 | 2.429315329 | 1.24E-85 | 5.93E-84 |
| **51** | eef1g | 26.67317 | 1.298143864 | 5.72E-91 | 2.92E-89 |
| **52** | zgc:65894 | 26.34632 | 5.259031296 | 3.03E-83 | 1.41E-81 |
| **53** | h2afx1 | 25.94992 | 1.579252958 | 4.28E-85 | 2.02E-83 |
| **54** | marcksb | 25.81384 | 1.759350777 | 4.02E-84 | 1.89E-82 |
| **55** | ywhag2 | 25.34646 | 3.982825279 | 8.79E-80 | 3.93E-78 |
| **56** | ppp1r14ba | 25.03397 | 4.914696217 | 2.45E-78 | 1.07E-76 |
| **57** | kif3cb | 24.97928 | 5.973311424 | 4.8E-78 | 2.09E-76 |
| **58** | dpysl2b | 24.6315 | 3.520289898 | 4.6E-77 | 1.98E-75 |
| **59** | dbi | 24.59581 | 1.687151074 | 1.34E-78 | 5.89E-77 |
| **60** | dpysl5a | 24.47413 | 3.198285341 | 1.41E-76 | 6.04E-75 |
| **61** | ank2b | 24.39385 | 2.932426214 | 1.78E-76 | 7.61E-75 |
| **62** | alcamb | 24.06705 | 4.352358341 | 9.86E-75 | 4.07E-73 |
| **63** | kif1aa | 23.97997 | 3.952137947 | 1.68E-74 | 6.94E-73 |
| **64** | gpm6ab | 23.67161 | 1.628277302 | 1.07E-75 | 4.47E-74 |
| **65** | nsg2 | 23.29612 | 2.445736885 | 1.78E-72 | 7.1E-71 |
| **66** | syt11a | 22.97694 | 2.690350533 | 3.31E-71 | 1.3E-69 |
| **67** | epb41a | 22.92062 | 2.399034262 | 7.89E-71 | 3.05E-69 |
| **68** | zgc:153426 | 22.86145 | 4.541609764 | 4.32E-70 | 1.65E-68 |
| **69** | tuba8l4 | 22.6835 | 1.430788279 | 2.96E-72 | 1.18E-70 |
| **70** | prdx2 | 22.47492 | 1.330635905 | 2.81E-70 | 1.08E-68 |
| **71** | ebf3a-1 | 22.44803 | 7.209118843 | 2.3E-68 | 8.6E-67 |
| **72** | zgc:101840 | 22.26582 | 4.986060619 | 9.67E-68 | 3.58E-66 |
| **73** | ywhah | 22.20234 | 2.360124111 | 4.08E-68 | 1.52E-66 |
| **74** | map7d2b | 22.16343 | 7.872359276 | 3.01E-67 | 1.1E-65 |
| **75** | ywhaqa | 21.70571 | 2.025176525 | 2.79E-66 | 1.01E-64 |
| **76** | csdc2a | 21.65125 | 2.832964182 | 1.01E-65 | 3.62E-64 |
| **77** | si:dkeyp-75h12.5 | 21.29547 | 2.491817474 | 2.06E-64 | 7.2E-63 |
| **78** | fkbp1aa | 21.15021 | 1.322054386 | 4.73E-65 | 1.67E-63 |
| **79** | hmgb1b | 21.08457 | 1.360093117 | 3.41E-65 | 1.21E-63 |
| **80** | si:busm1-57f23.1 | 21.06547 | 4.324352264 | 3.74E-63 | 1.28E-61 |
| **81** | hsp90ab1 | 20.98585 | 1.071817756 | 2.65E-64 | 9.21E-63 |
| **82** | gng5 | 20.91486 | 1.871556759 | 2.65E-63 | 9.1E-62 |
| **83** | dclk1b | 20.90953 | 4.903087139 | 1.88E-62 | 6.38E-61 |
| **84** | add2 | 20.89866 | 5.772411823 | 2.31E-62 | 7.84E-61 |
| **85** | prph | 20.81114 | 8.07297039 | 5.57E-62 | 1.87E-60 |
| **86** | si:ch211-284f22.3 | 20.77369 | 3.973699093 | 5.57E-62 | 1.87E-60 |
| **87** | cd99l2 | 20.75372 | 4.002908707 | 7.09E-62 | 2.38E-60 |
| **88** | anxa5b | 20.66391 | 4.878666401 | 1.75E-61 | 5.81E-60 |
| **89** | si:ch211-195b15.8 | 20.33169 | 3.041670561 | 1.67E-60 | 5.47E-59 |
| **90** | si:dkey-28b4.7 | 20.18958 | 2.106191397 | 3.31E-60 | 1.07E-58 |
| **91** | scg2b | 20.14479 | 6.470963001 | 2.14E-59 | 6.87E-58 |
| **92** | rps19 | 19.99676 | 0.791598558 | 9.78E-64 | 3.37E-62 |
| **93** | tbcb | 19.79526 | 1.86586082 | 1.1E-58 | 3.47E-57 |
| **94** | adcyap1b | 19.61666 | 8.515849113 | 2.72E-57 | 8.32E-56 |
| **95** | coro1cb | 19.55775 | 2.983227015 | 2.43E-57 | 7.46E-56 |
| **96** | zc4h2 | 19.47209 | 2.499441147 | 3.48E-57 | 1.06E-55 |
| **97** | kif5aa | 19.36726 | 4.668578148 | 2.18E-56 | 6.54E-55 |
| **98** | eml1 | 19.28413 | 3.152324915 | 3.41E-56 | 1.02E-54 |
| **99** | sox11b | 19.26969 | 2.320665359 | 1.36E-56 | 4.13E-55 |

**Cluster 11**

| | names | scores | logfoldchanges | pvals | pvals_adj |
|---|---|---|---|---|---|
| **0** | rbp4l | 69.7916 | 5.663568497 | 6.7E-246 | 1.1E-243 |
| **1** | thrb | 68.77347 | 4.534096718 | 1.2E-247 | 1.9E-245 |
| **2** | aanat2 | 67.73853 | 6.9752388 | 7.5E-232 | 1E-229 |
| **3** | rcvrn2 | 62.65492 | 5.942514896 | 5.1E-220 | 6.4E-218 |
| **4** | neurod1 | 51.49628 | 3.872107744 | 6.9E-196 | 7.1E-194 |
| **5** | guk1b | 49.40574 | 6.647085667 | 6.9E-179 | 6.1E-177 |
| **6** | arl13a | 48.37612 | 4.121647835 | 3.2E-181 | 2.8E-179 |
| **7** | atp5mc3b | 46.36296 | 2.320496559 | 5E-180 | 4.5E-178 |
| **8** | ppdpfa | 43.9301 | 4.699728012 | 4.5E-162 | 3.5E-160 |
| **9** | ckmt2a | 43.92736 | 6.408274651 | 5E-161 | 3.9E-159 |
| **10** | si:dkey-72l14.3 | 42.8831 | 6.207275391 | 5.5E-157 | 4.1E-155 |
| **11** | fkbp1b | 39.92556 | 3.357848644 | 3.5E-149 | 2.4E-147 |
| **12** | crx | 39.71903 | 3.784885883 | 4.6E-148 | 3.1E-146 |
| **13** | zgc:109965 | 39.18085 | 4.470720768 | 6.7E-145 | 4.4E-143 |
| **14** | atp2b1b | 38.81934 | 4.906696796 | 7.7E-143 | 5E-141 |
| **15** | atp5f1b | 37.82442 | 1.955410361 | 4.1E-144 | 2.7E-142 |
| **16** | atp5pd | 35.82613 | 2.040931463 | 9.3E-136 | 5.6E-134 |
| **17** | anp32e | 35.72473 | 2.075330734 | 7.2E-137 | 4.4E-135 |
| **18** | sypb | 35.10272 | 4.73572731 | 2.1E-128 | 1.2E-126 |
| **19** | cox4i1 | 34.78309 | 1.695576906 | 5.5E-133 | 3.2E-131 |
| **20** | six7 | 34.2225 | 4.387979984 | 1.7E-125 | 9.2E-124 |
| **21** | elovl4b | 33.96679 | 4.435673714 | 4.8E-124 | 2.6E-122 |
| **22** | cox6a1 | 33.06525 | 1.609958172 | 1.1E-125 | 5.9E-124 |
| **23** | slc25a5 | 32.63343 | 1.301927686 | 2.8E-122 | 1.4E-120 |
| **24** | atp5mc1 | 32.58017 | 1.931897402 | 1.4E-121 | 7.2E-120 |
| **25** | ndufa4l | 32.05808 | 1.575374603 | 9.5E-123 | 5E-121 |
| **26** | gpx4b | 30.99536 | 2.202866554 | 8.4E-114 | 4.1E-112 |
| **27** | snap25b | 30.96257 | 3.156216621 | 2.6E-113 | 1.3E-111 |
| **28** | si:dkey-44g23.5 | 30.96007 | 4.700011253 | 2.4E-111 | 1.1E-109 |
| **29** | si:ch73-28h20.1 | 30.71418 | 3.563786745 | 1.5E-112 | 7.3E-111 |
| **30** | xbp1 | 30.67335 | 1.851337671 | 2.5E-113 | 1.2E-111 |
| **31** | atp6v0cb | 30.52959 | 2.497104406 | 1.6E-111 | 7.5E-110 |
| **32** | h3f3b.1-2 | 30.08783 | 1.688842773 | 4.9E-112 | 2.4E-110 |
| **33** | atp5po | 30.03177 | 1.895007849 | 2.1E-110 | 1E-108 |
| **34** | cox5aa | 30.00272 | 1.83091867 | 1.2E-110 | 5.7E-109 |
| **35** | hmgn6 | 29.99281 | 1.222048879 | 1.1E-116 | 5.3E-115 |
| **36** | COX5B | 29.74447 | 1.836199522 | 2.2E-109 | 1E-107 |
| **37** | slc25a3a | 29.61564 | 5.821687222 | 1.6E-105 | 7.3E-104 |
| **38** | rps27.1 | 29.42719 | 1.127908707 | 2.9E-116 | 1.5E-114 |
| **39** | cox8a | 28.84851 | 1.458173037 | 5.6E-106 | 2.5E-104 |
| **40** | rps5 | 28.80428 | 1.122909427 | 1E-113 | 5E-112 |
| **41** | rps7 | 28.566 | 1.099828839 | 3.6E-112 | 1.7E-110 |
| **42** | rpl19 | 28.1412 | 1.180482626 | 2.5E-108 | 1.2E-106 |
| **43** | rps6 | 28.07899 | 1.131963134 | 1E-109 | 4.8E-108 |
| **44** | cox4i2 | 27.99816 | 2.492438793 | 4.3E-100 | 1.81E-98 |
| **45** | rxrgb | 27.97975 | 3.893286228 | 4.8E-99 | 2E-97 |
| **46** | cox7a2a | 27.63653 | 1.958716869 | 2.6E-99 | 1.1E-97 |
| **47** | atp5fa1 | 27.63401 | 1.587958097 | 6.7E-100 | 2.8E-98 |
| **48** | mt-co2 | 27.30873 | 1.423971176 | 8.61E-98 | 3.54E-96 |
| **49** | rps4x | 27.05352 | 1.099805474 | 1.7E-103 | 7.2E-102 |
| **50** | rs1a | 26.95639 | 4.589650154 | 1.66E-94 | 6.42E-93 |
| **51** | rpl18a | 26.94043 | 1.020688891 | 6.2E-104 | 2.7E-102 |
| **52** | rpl17 | 26.9024 | 1.017889142 | 6.1E-104 | 2.7E-102 |
| **53** | unc119b | 26.49944 | 3.737752438 | 1.59E-92 | 6.05E-91 |
| **54** | ckbb | 26.37506 | 1.418716669 | 7.45E-97 | 3.01E-95 |
| **55** | ndrg1a | 26.36163 | 4.804290771 | 1.07E-91 | 4E-90 |
| **56** | zgc:153441 | 26.18509 | 4.608101368 | 5.78E-91 | 2.15E-89 |
| **57** | tulp1a | 26.04637 | 4.74846077 | 2.63E-90 | 9.66E-89 |
| **58** | rpl7a | 26.03493 | 1.007828712 | 2.9E-99 | 1.21E-97 |
| **59** | selenot1a | 25.78408 | 1.917764544 | 8.79E-91 | 3.26E-89 |
| **60** | mt-nd1 | 25.76087 | 1.508603454 | 5.65E-91 | 2.1E-89 |
| **61** | mdh2 | 25.66134 | 1.892920494 | 4.47E-90 | 1.63E-88 |
| **62** | rpl15 | 25.65539 | 1.094471574 | 3.71E-96 | 1.47E-94 |
| **63** | rpl3 | 25.59778 | 1.012274384 | 4.53E-96 | 1.79E-94 |
| **64** | rps13 | 25.45943 | 0.947067499 | 1.88E-96 | 7.52E-95 |
| **65** | rpl10 | 25.42695 | 0.981639445 | 1.37E-95 | 5.39E-94 |
| **66** | opn6a | 25.41356 | 4.374788761 | 1.35E-87 | 4.76E-86 |
| **67** | tmem244 | 25.32793 | 4.210472584 | 2.79E-87 | 9.79E-86 |
| **68** | rack1 | 25.10625 | 1.089299202 | 1.36E-92 | 5.17E-91 |
| **69** | mt-co3 | 25.00665 | 1.357063293 | 2.78E-87 | 9.77E-86 |
| **70** | sall1a | 24.9954 | 3.489038706 | 4.77E-86 | 1.65E-84 |
| **71** | nr2e3 | 24.94977 | 3.009399652 | 1.85E-86 | 6.44E-85 |
| **72** | faua | 24.79133 | 0.986643851 | 4.82E-92 | 1.83E-90 |
| **73** | mt-atp6 | 24.7508 | 1.446355939 | 3.34E-86 | 1.16E-84 |
| **74** | rpl13 | 24.65279 | 1.021046042 | 1.54E-90 | 5.67E-89 |
| **75** | uba52 | 24.62097 | 0.881376266 | 1.13E-91 | 4.24E-90 |
| **76** | si:dkey-220f10.4 | 24.58719 | 4.560071468 | 6.72E-84 | 2.26E-82 |
| **77** | rpl8 | 24.55198 | 0.955642045 | 1.22E-90 | 4.51E-89 |
| **78** | atp5f1d | 24.37313 | 1.674706936 | 1.44E-84 | 4.88E-83 |
| **79** | atp5if1b | 24.27318 | 2.07263732 | 1.86E-83 | 6.18E-82 |
| **80** | eef1g | 24.20253 | 0.985345185 | 8.41E-90 | 3.05E-88 |
| **81** | arl3l1 | 24.16562 | 5.169780254 | 6.57E-82 | 2.15E-80 |
| **82** | rps19 | 24.13347 | 0.902123153 | 5.34E-90 | 1.95E-88 |
| **83** | ldhbb | 24.12926 | 4.918907166 | 9.2E-82 | 3E-80 |
| **84** | rpl30 | 24.08674 | 1.012762904 | 5.74E-87 | 2E-85 |
| **85** | rpl7 | 23.84151 | 0.960256696 | 7.73E-87 | 2.68E-85 |
| **86** | rps9 | 23.83957 | 0.917826831 | 9.32E-88 | 3.29E-86 |
| **87** | rpl10a | 23.79163 | 0.925878108 | 5.68E-87 | 1.98E-85 |
| **88** | rps8a | 23.76958 | 0.938959181 | 3.36E-87 | 1.17E-85 |
| **89** | atp5pf | 23.61867 | 1.753030896 | 5.21E-81 | 1.67E-79 |
| **90** | naca | 23.55142 | 0.975248992 | 3.37E-84 | 1.14E-82 |
| **91** | rpl28 | 23.45271 | 0.9391312 | 8.06E-85 | 2.75E-83 |
| **92** | inaa | 23.39441 | 6.496417522 | 2.24E-78 | 6.93E-77 |
| **93** | zgc:103625 | 23.17529 | 5.125854969 | 1.67E-77 | 5.09E-76 |
| **94** | gngt2a | 23.17059 | 4.153347015 | 1.15E-77 | 3.53E-76 |
| **95** | rpl32 | 23.1554 | 0.91199559 | 7.12E-84 | 2.39E-82 |
| **96** | slc25a18 | 23.04619 | 2.834618568 | 2.2E-77 | 6.69E-76 |
| **97** | tcima | 22.94127 | 4.042626858 | 1.18E-76 | 3.56E-75 |
| **98** | rpl13a | 22.89417 | 0.899820268 | 2.08E-81 | 6.75E-80 |
| **99** | pcdh10a | 22.84693 | 4.186196804 | 3.49E-76 | 1.05E-74 |

**Cluster 12**

| | names | scores | logfoldchanges | pvals | pvals_adj |
|---|---|---|---|---|---|
| 0 | arl13a | 53.44701 | 4.167642593 | 9.5E-198 | 8.4E-196 |
| 1 | rbp4l | 50.26901 | 5.163287163 | 1.3E-184 | 1E-182 |
| 2 | gngt2a | 46.63506 | 5.043726444 | 3E-170 | 2.1E-168 |
| 3 | ppdpfa | 40.0969 | 4.24877882 | 9.5E-148 | 5.5E-146 |
| 4 | cxxc5a | 38.50266 | 2.775964499 | 1.7E-144 | 9.4E-143 |
| 5 | ckmt2a | 37.29204 | 5.475167274 | 2.3E-136 | 1.2E-134 |
| 6 | zgc:109965 | 36.8304 | 4.480729103 | 8.9E-135 | 4.5E-133 |
| 7 | neurod1 | 36.21203 | 3.369781733 | 7.8E-136 | 4E-134 |
| 8 | atp5mc3b | 35.98587 | 1.439428329 | 6.2E-145 | 3.4E-143 |
| 9 | fkbp1b | 35.94051 | 2.987205029 | 2.1E-133 | 1E-131 |
| 10 | si:ch73-28h20.1 | 35.54596 | 5.054292679 | 8.8E-130 | 4.2E-128 |
| 11 | crx | 34.26119 | 3.467672348 | 7.1E-126 | 3.2E-124 |
| 12 | rpl19 | 33.96902 | 1.001766443 | 2E-152 | 1.2E-150 |
| 13 | anp32e | 31.89721 | 1.759705305 | 7.9E-121 | 3.4E-119 |
| 14 | nr2e3 | 31.62177 | 3.179868698 | 9E-116 | 3.7E-114 |
| 15 | h3f3b.1-2 | 31.41482 | 1.656202912 | 4.4E-118 | 1.8E-116 |
| 16 | zgc:153441 | 31.2162 | 4.905676842 | 7.8E-112 | 3.1E-110 |
| 17 | syt5b | 31.06565 | 4.943226814 | 3.6E-111 | 1.4E-109 |
| 18 | rps5 | 30.94678 | 0.935383141 | 1.2E-134 | 5.9E-133 |
| 19 | cox4i2 | 30.92143 | 2.457738638 | 1E-112 | 4.1E-111 |
| 20 | rps7 | 30.71694 | 0.876183152 | 3.7E-135 | 1.9E-133 |
| 21 | snap25b | 29.92464 | 3.075303555 | 2.8E-108 | 1.1E-106 |
| 22 | tmem244 | 29.14986 | 4.427742958 | 2.3E-103 | 8.3E-102 |
| 23 | rpl7a | 28.83632 | 0.843262672 | 3.1E-123 | 1.3E-121 |
| 24 | rps4x | 28.46353 | 0.889535069 | 3.3E-118 | 1.4E-116 |
| 25 | gpx4b | 28.223 | 2.046368361 | 3.5E-101 | 1.2E-99 |
| 26 | rpl8 | 28.17669 | 0.764190078 | 1.8E-120 | 7.6E-119 |
| 27 | atp2b1b | 27.91111 | 3.994916677 | 2.35E-98 | 7.84E-97 |
| 28 | laptm4b | 27.8376 | 1.921810865 | 7.3E-100 | 2.48E-98 |
| 29 | atp6v0cb | 27.81422 | 2.232342243 | 1.9E-99 | 6.43E-98 |
| 30 | ndrg1a | 27.62278 | 4.71707201 | 9.37E-97 | 3.04E-95 |
| 31 | rpl13 | 27.513 | 0.828255534 | 4.8E-113 | 1.9E-111 |
| 32 | rack1 | 27.1977 | 0.849814892 | 4.1E-111 | 1.6E-109 |
| 33 | rpl17 | 26.57596 | 0.806814134 | 5E-108 | 1.9E-106 |
| 34 | sypb | 26.56752 | 3.859397888 | 1.61E-92 | 5E-91 |
| 35 | rpl15 | 26.52748 | 0.881689548 | 2.3E-106 | 8.3E-105 |
| 36 | rps27.1 | 26.41185 | 0.897524774 | 2.2E-103 | 8E-102 |
| 37 | hmgn6 | 26.19894 | 1.0688411 | 1.14E-97 | 3.76E-96 |
| 38 | rpl18 | 26.04649 | 0.781025887 | 9.6E-105 | 3.4E-103 |
| 39 | rpl10 | 26.02632 | 0.736429036 | 9.3E-107 | 3.4E-105 |
| 40 | opn6a | 25.99137 | 4.398880959 | 1.06E-89 | 3.15E-88 |
| 41 | rps8a | 25.96126 | 0.771639526 | 1E-105 | 3.7E-104 |
| 42 | rpl18a | 25.87892 | 0.797070861 | 2.4E-103 | 8.5E-102 |
| 43 | ckbb | 25.84623 | 1.011829138 | 3.3E-99 | 1.12E-97 |
| 44 | eef1g | 25.27461 | 0.824545205 | 1.9E-100 | 6.4E-99 |
| 45 | rpl7 | 25.19827 | 0.78865844 | 8.8E-99 | 2.95E-97 |
| 46 | tmx3a | 25.13877 | 4.64642477 | 6.63E-86 | 1.91E-84 |
| 47 | ndrg1b | 24.68048 | 3.608685493 | 2.8E-84 | 7.94E-83 |
| 48 | rpl3 | 24.63154 | 0.774188042 | 1.54E-95 | 4.93E-94 |
| 49 | rpl28 | 24.57047 | 0.798832715 | 4.02E-94 | 1.27E-92 |
| 50 | rps6 | 24.33335 | 0.859112561 | 8.79E-93 | 2.74E-91 |
| 51 | rpl11 | 23.90525 | 0.71523881 | 1.81E-93 | 5.67E-92 |
| 52 | rpl10a | 23.88967 | 0.715114713 | 1.99E-92 | 6.16E-91 |
| 53 | selenot1a | 23.56553 | 1.665691376 | 1.22E-80 | 3.32E-79 |
| 54 | aplnra | 23.55503 | 4.454494953 | 6.72E-79 | 1.78E-77 |
| 55 | rps9 | 23.5456 | 0.662600636 | 7.51E-93 | 2.35E-91 |
| 56 | rpl32 | 23.46778 | 0.702766776 | 1.14E-90 | 3.45E-89 |
| 57 | si:dkey-220f10.4 | 23.39208 | 4.439289093 | 3.12E-78 | 8.19E-77 |
| 58 | rpl13a | 23.33629 | 0.704063058 | 1.48E-87 | 4.32E-86 |
| 59 | slc25a3a | 23.30503 | 4.471027851 | 6.24E-78 | 1.64E-76 |
| 60 | ndufa4l | 23.14509 | 1.216362238 | 3.03E-80 | 8.18E-79 |
| 61 | tmsb2 | 23.12352 | 3.023330688 | 6.34E-78 | 1.66E-76 |
| 62 | atp5f1b | 22.79394 | 1.072304487 | 2E-78 | 5.26E-77 |
| 63 | rps10 | 22.68702 | 0.70667851 | 1.72E-85 | 4.96E-84 |
| 64 | fstl5 | 22.6635 | 4.286492825 | 5.45E-75 | 1.37E-73 |
| 65 | tulp1a | 22.5865 | 4.166074753 | 1.02E-74 | 2.55E-73 |
| 66 | jun | 22.32785 | 1.05002296 | 1.07E-81 | 2.94E-80 |
| 67 | slc25a5 | 22.25247 | 0.643797755 | 4.19E-77 | 1.09E-75 |
| 68 | faua | 22.06378 | 0.750687897 | 4.19E-80 | 1.13E-78 |
| 69 | zgc:112294 | 21.98085 | 4.350487232 | 6.03E-72 | 1.46E-70 |
| 70 | ldhbb | 21.93536 | 4.19235611 | 8.53E-72 | 2.06E-70 |
| 71 | rps2 | 21.90655 | 0.706074417 | 8.18E-81 | 2.23E-79 |
| 72 | cldn2 | 21.61791 | 4.262046814 | 2.53E-70 | 5.96E-69 |
| 73 | mt-co2 | 21.54855 | 0.913247168 | 5.62E-72 | 1.36E-70 |
| 74 | daam1a | 21.53014 | 2.755152464 | 3.11E-70 | 7.32E-69 |
| 75 | rps23 | 21.25978 | 0.711747944 | 4.8E-76 | 1.22E-74 |
| 76 | pdcl | 21.21949 | 1.655368805 | 1.1E-69 | 2.57E-68 |
| 77 | mt-co3 | 21.14372 | 0.878739655 | 3.05E-70 | 7.18E-69 |
| 78 | atp5pd | 20.9451 | 1.225665212 | 4.36E-69 | 1.01E-67 |
| 79 | rs1a | 20.81515 | 4.391170979 | 9.59E-67 | 2.16E-65 |
| 80 | ptmab | 20.67576 | 0.784829199 | 3.01E-71 | 7.19E-70 |
| 81 | prdm1a | 20.59002 | 3.912133694 | 9.2E-66 | 2.04E-64 |
| 82 | hmga1b | 20.27947 | 3.729668617 | 2.22E-64 | 4.82E-63 |
| 83 | elovl4b | 20.1759 | 3.143271208 | 3.78E-64 | 8.18E-63 |
| 84 | rcvrn2 | 20.09399 | 3.323274374 | 8.26E-64 | 1.78E-62 |
| 85 | rpgrb | 20.08928 | 2.939602613 | 1.08E-63 | 2.3E-62 |
| 86 | rps3a | 19.78378 | 0.591439784 | 9.08E-70 | 2.12E-68 |
| 87 | rpl21 | 19.62358 | 0.611033022 | 2.94E-68 | 6.75E-67 |
| 88 | rps13 | 19.59334 | 0.660726368 | 6.8E-67 | 1.53E-65 |
| 89 | rps3 | 19.5315 | 0.688178957 | 3.06E-66 | 6.84E-65 |
| 90 | prph2a | 19.50964 | 4.992937565 | 7.77E-61 | 1.62E-59 |
| 91 | rpl30 | 19.46901 | 0.742855251 | 1.27E-64 | 2.78E-63 |
| 92 | unc119b | 19.31224 | 2.952768087 | 3.33E-60 | 6.89E-59 |
| 93 | nptna | 19.24524 | 3.83417654 | 9.47E-60 | 1.94E-58 |
| 94 | rps27a | 19.16818 | 0.580067635 | 8.28E-66 | 1.83E-64 |
| 95 | uba52 | 19.08483 | 0.63183856 | 9.32E-64 | 2E-62 |
| 96 | xbp1 | 18.98545 | 1.097527027 | 7.07E-60 | 1.45E-58 |
| 97 | rpl5b | 18.95905 | 0.868398011 | 3.06E-61 | 6.38E-60 |
| 98 | zgc:103625 | 18.7948 | 4.230988503 | 1.03E-57 | 2.06E-56 |
| 99 | naca | 18.60692 | 0.707758009 | 3.7E-60 | 7.64E-59 |

**Cluster 13**

| | names | scores | logfoldchanges | pvals | pvals_adj |
|---|---|---|---|---|---|
| **0** | elavl3 | 47.59343 | 4.885855198 | 9E-157 | 6.8E-155 |
| **1** | hmgb3a | 39.8751 | 2.369702578 | 1.2E-136 | 7.3E-135 |
| **2** | zc4h2 | 39.0952 | 3.821972609 | 3.5E-130 | 2E-128 |
| **3** | marcksl1b | 36.37799 | 2.3721416 | 9E-127 | 4.9E-125 |
| **4** | si:ch73-1a9.3 | 33.59291 | 1.881837606 | 3E-117 | 1.5E-115 |
| **5** | snap25b | 33.55449 | 4.011558533 | 1E-111 | 4.9E-110 |
| **6** | stx1b | 32.71359 | 5.640387535 | 8.4E-108 | 3.8E-106 |
| **7** | ptmab | 32.12175 | 1.364897013 | 2.4E-116 | 1.2E-114 |
| **8** | nova2 | 31.29289 | 2.577271461 | 3.4E-105 | 1.5E-103 |
| **9** | ywhah | 31.12801 | 2.594067097 | 3.5E-104 | 1.5E-102 |
| **10** | stmn1b | 30.46353 | 2.355851412 | 2.6E-104 | 1.1E-102 |
| **11** | gng3 | 29.19424 | 4.01731348 | 9.38E-96 | 3.62E-94 |
| **12** | h2afx1 | 28.89923 | 1.801385283 | 6.54E-98 | 2.63E-96 |
| **13** | ndrg4 | 28.50275 | 2.811211348 | 6.49E-94 | 2.44E-92 |
| **14** | tkta | 27.96958 | 3.454151392 | 2.65E-91 | 9.68E-90 |
| **15** | ppp1r14c | 27.66105 | 5.04389286 | 1.3E-89 | 4.67E-88 |
| **16** | slc32a1 | 26.97668 | 6.620533466 | 6.97E-87 | 2.42E-85 |
| **17** | hmgn7 | 26.25145 | 1.725021839 | 1.78E-86 | 6.15E-85 |
| **18** | ptmaa | 26.10138 | 1.471295714 | 2.1E-88 | 7.4E-87 |
| **19** | syt1a | 25.45311 | 5.426211357 | 3.55E-81 | 1.15E-79 |
| **20** | gpm6aa | 25.39178 | 1.586851835 | 1.55E-83 | 5.21E-82 |
| **21** | stxbp1a | 24.8139 | 5.629206657 | 1.16E-78 | 3.63E-77 |
| **22** | tuba1c | 24.06178 | 2.418558598 | 6.68E-77 | 2.03E-75 |
| **23** | hmgn6 | 23.91729 | 1.21454072 | 6.45E-79 | 2.02E-77 |
| **24** | rbfox2 | 23.27165 | 3.688537836 | 5.68E-73 | 1.6E-71 |
| **25** | atp6v0cb | 23.19559 | 2.439435244 | 4.42E-73 | 1.26E-71 |
| **26** | h2afy2 | 23.18473 | 2.073511839 | 3.04E-73 | 8.69E-72 |
| **27** | ywhag2 | 23.18036 | 3.996520758 | 1.78E-72 | 4.97E-71 |
| **28** | ccni | 22.97561 | 1.92903471 | 1.7E-72 | 4.77E-71 |
| **29** | mdkb | 22.81258 | 1.929414749 | 1.09E-72 | 3.05E-71 |
| **30** | ppiab | 22.71882 | 1.180059552 | 1.35E-72 | 3.79E-71 |
| **31** | pax6a | 22.39902 | 3.533084869 | 2.18E-69 | 5.91E-68 |
| **32** | atp6v1g1 | 22.28578 | 2.394785166 | 2.37E-69 | 6.44E-68 |
| **33** | h3f3d | 22.00778 | 1.02500689 | 6.89E-70 | 1.89E-68 |
| **34** | kdm6bb | 21.434 | 3.426253557 | 1.4E-65 | 3.64E-64 |
| **35** | ppp1r14ba | 21.39812 | 3.622980595 | 1.72E-65 | 4.45E-64 |
| **36** | jpt1b | 20.91422 | 1.772910357 | 3.22E-64 | 8.14E-63 |
| **37** | atp6v1e1b | 20.75832 | 2.281619787 | 3.67E-63 | 9.09E-62 |
| **38** | apc | 20.70758 | 3.184223175 | 1.14E-62 | 2.79E-61 |
| **39** | pax10 | 20.51826 | 7.094687939 | 1.24E-61 | 2.99E-60 |
| **40** | mab21l1 | 20.42645 | 2.67046833 | 1.17E-61 | 2.82E-60 |
| **41** | rtn1b | 20.4185 | 2.91753459 | 1.25E-61 | 3.01E-60 |
| **42** | calm2b | 20.40911 | 1.887760043 | 5.5E-62 | 1.34E-60 |
| **43** | hsp90ab1 | 20.29822 | 0.873704851 | 6.69E-63 | 1.65E-61 |
| **44** | si:ch73-290k24.5 | 20.2575 | 5.234352112 | 1.22E-60 | 2.9E-59 |
| **45** | h3f3c | 20.25117 | 1.489620805 | 6.44E-62 | 1.57E-60 |
| **46** | tfap2a | 20.22489 | 6.172959328 | 1.82E-60 | 4.29E-59 |
| **47** | si:dkey-280e21.3 | 20.09857 | 3.644492388 | 3.31E-60 | 7.79E-59 |
| **48** | basp1 | 19.91347 | 4.168925762 | 2.62E-59 | 6.07E-58 |
| **49** | sncb | 19.70108 | 4.021833897 | 1.77E-58 | 4.01E-57 |
| **50** | sumo2b | 19.41626 | 1.408165812 | 2.63E-58 | 5.95E-57 |
| **51** | gng2 | 19.25251 | 5.368483543 | 1.44E-56 | 3.17E-55 |
| **52** | elmo1 | 19.24801 | 2.884595394 | 9.01E-57 | 1.99E-55 |
| **53** | rbfox1 | 18.74553 | 5.295084953 | 1.53E-54 | 3.26E-53 |
| **54** | vamp2 | 18.55072 | 2.290856838 | 4.48E-54 | 9.45E-53 |
| **55** | dnajc5aa | 18.41609 | 3.033177376 | 2.39E-53 | 5E-52 |
| **56** | csdc2a | 18.32683 | 2.371032476 | 3.41E-53 | 7.11E-52 |
| **57** | eno2 | 18.22068 | 3.144123316 | 1.51E-52 | 3.11E-51 |
| **58** | si:dkey-81l17.6 | 18.19662 | 3.702046871 | 2.14E-52 | 4.39E-51 |
| **59** | pax6b | 17.9009 | 2.382846832 | 2.1E-51 | 4.23E-50 |
| **60** | atp1a3a | 17.81363 | 3.531244278 | 7.26E-51 | 1.45E-49 |
| **61** | gpm6ab | 17.69015 | 1.382048368 | 2.81E-51 | 5.63E-50 |
| **62** | cspg5a | 17.64552 | 1.714193344 | 9.46E-51 | 1.88E-49 |
| **63** | fscn1a | 17.59551 | 2.13964653 | 3.03E-50 | 6E-49 |
| **64** | rab11bb | 17.48482 | 3.434072495 | 1.5E-49 | 2.92E-48 |
| **65** | ndrg2 | 17.40715 | 2.383869886 | 2.27E-49 | 4.4E-48 |
| **66** | cnrip1a | 17.22821 | 2.926807642 | 1.48E-48 | 2.83E-47 |
| **67** | tuba2 | 17.21075 | 4.067759991 | 2.11E-48 | 4.01E-47 |
| **68** | tmsb4x | 17.08338 | 1.030745506 | 8.39E-49 | 1.61E-47 |
| **69** | zgc:100920 | 16.91031 | 4.654830456 | 3.84E-47 | 7.18E-46 |
| **70** | scrt2 | 16.76306 | 3.020741463 | 9.39E-47 | 1.75E-45 |
| **71** | tiam1a | 16.74403 | 4.688462257 | 1.79E-46 | 3.32E-45 |
| **72** | gad2 | 16.70621 | 6.308121681 | 2.82E-46 | 5.19E-45 |
| **73** | fez1 | 16.56593 | 1.960237145 | 4.57E-46 | 8.4E-45 |
| **74** | hp1bp3 | 16.21523 | 1.792403579 | 1.18E-44 | 2.11E-43 |
| **75** | nrxn1a | 16.14522 | 4.830283642 | 4.61E-44 | 8.16E-43 |
| **76** | map1aa | 16.12482 | 2.401688337 | 3.64E-44 | 6.46E-43 |
| **77** | vdac3 | 16.0427 | 1.638497233 | 5.18E-44 | 9.15E-43 |
| **78** | hist2h2l | 16.0084 | 1.765642881 | 7.22E-44 | 1.27E-42 |
| **79** | gpr85 | 15.94105 | 2.557376146 | 2.21E-43 | 3.86E-42 |
| **80** | nsfa | 15.93045 | 3.221651077 | 2.77E-43 | 4.82E-42 |
| **81** | vdac1 | 15.88041 | 1.517517805 | 2.51E-43 | 4.39E-42 |
| **82** | st8sia5 | 15.80181 | 4.730092049 | 1.08E-42 | 1.87E-41 |
| **83** | cdc42 | 15.73744 | 1.584969997 | 1E-42 | 1.74E-41 |
| **84** | calm3a | 15.70304 | 1.509706497 | 1.27E-42 | 2.19E-41 |
| **85** | itm2ca | 15.58153 | 3.611167192 | 7.48E-42 | 1.27E-40 |
| **86** | fkbp1aa | 15.57737 | 1.156489015 | 2.57E-42 | 4.4E-41 |
| **87** | si:ch73-119p20.1 | 15.39997 | 4.765375614 | 4.38E-41 | 7.32E-40 |
| **88** | foxg1b | 15.35191 | 2.092312098 | 4.35E-41 | 7.28E-40 |
| **89** | si:dkey-276j7.1 | 15.35012 | 1.675403476 | 2.99E-41 | 5.02E-40 |
| **90** | ywhaqb | 15.33956 | 1.09424603 | 2.43E-41 | 4.09E-40 |
| **91** | sv2a | 15.30951 | 5.490233421 | 1.05E-40 | 1.74E-39 |
| **92** | fabp3 | 15.28771 | 1.174806237 | 2.47E-41 | 4.15E-40 |
| **93** | marcksl1a | 15.25371 | 1.04894352 | 3.07E-41 | 5.15E-40 |
| **94** | calm2a | 15.20846 | 1.650955558 | 1.42E-40 | 2.34E-39 |
| **95** | rnasekb | 15.11282 | 1.556983352 | 3.16E-40 | 5.16E-39 |
| **96** | hmgn3 | 15.08429 | 1.610709548 | 3.66E-40 | 5.96E-39 |
| **97** | cbx1b | 15.01925 | 2.906497717 | 1.2E-39 | 1.92E-38 |
| **98** | h3f3b.1-1 | 15.01093 | 1.021741509 | 3.14E-40 | 5.13E-39 |
| **99** | fam49al | 14.97274 | 2.882889032 | 1.8E-39 | 2.87E-38 |

**Cluster 14**

| | names | scores | logfoldchanges | pvals | pvals_adj |
|---|---|---|---|---|---|
| 0 | gng13b | 45.72647 | 6.802714348 | 4.25E-42 | 3.88E-41 |
| 1 | mdkb | 38.88173 | 4.460173607 | 3.58E-39 | 3.12E-38 |
| 2 | calm2b | 28.64004 | 3.10985136 | 8.11E-33 | 6.38E-32 |
| 3 | rbfox1 | 28.00863 | 7.15584898 | 7.55E-32 | 5.85E-31 |
| 4 | scrt2 | 24.92979 | 5.27226162 | 1.35E-29 | 1E-28 |
| 5 | tiam1a | 24.80428 | 6.726092815 | 2.03E-29 | 1.5E-28 |
| 6 | gnb1a | 24.47663 | 3.227253675 | 2.31E-29 | 1.71E-28 |
| 7 | ppp1r14c | 23.27609 | 5.525703907 | 3.32E-28 | 2.39E-27 |
| 8 | olfm1b | 23.01315 | 5.576354504 | 5.73E-28 | 4.1E-27 |
| 9 | ndrg4 | 22.51121 | 3.05749321 | 8.46E-28 | 6.03E-27 |
| 10 | pvalb6 | 21.84878 | 9.853527069 | 6.42E-27 | 4.51E-26 |
| 11 | ptmaa | 20.86428 | 1.855203867 | 6.96E-27 | 4.89E-26 |
| 12 | stx1b | 20.52764 | 5.374100685 | 9.1E-26 | 6.25E-25 |
| 13 | sox2 | 20.08649 | 3.241815329 | 1.74E-25 | 1.19E-24 |
| 14 | nrxn1a | 19.6996 | 6.766798019 | 6.04E-25 | 4.07E-24 |
| 15 | CR855337.1 | 19.32253 | 6.077991009 | 1.38E-24 | 9.24E-24 |
| 16 | rcan3 | 18.9845 | 6.211945534 | 3E-24 | 1.99E-23 |
| 17 | marcksl1b | 18.0022 | 2.097160578 | 1.04E-23 | 6.82E-23 |
| 18 | snap25b | 17.46151 | 3.774785757 | 8.77E-23 | 5.64E-22 |
| 19 | fkbp1b | 16.53998 | 3.229223251 | 8.54E-22 | 5.37E-21 |
| 20 | stmn1b | 16.17747 | 2.007080555 | 9.33E-22 | 5.86E-21 |
| 21 | syt1a | 15.97197 | 5.825585842 | 4.47E-21 | 2.77E-20 |
| 22 | ppdpfb | 15.76896 | 2.26936388 | 5.23E-21 | 3.23E-20 |
| 23 | elavl3 | 15.47612 | 3.887323618 | 1.37E-20 | 8.36E-20 |
| 24 | pcdh19 | 14.70898 | 5.537436008 | 1.3E-19 | 7.77E-19 |
| 25 | si:dkey-35i13.1 | 14.35941 | 7.969618797 | 3.46E-19 | 2.04E-18 |
| 26 | si:ch73-119p20.1 | 13.96951 | 5.74709034 | 1.01E-18 | 5.89E-18 |
| 27 | tmem178b | 13.47456 | 4.701290607 | 4.06E-18 | 2.34E-17 |
| 28 | tkta | 13.37233 | 3.268822908 | 5.06E-18 | 2.91E-17 |
| 29 | slc18a3a | 13.14749 | 8.63232708 | 1.09E-17 | 6.2E-17 |
| 30 | calm3a | 12.85786 | 2.206824303 | 2.12E-17 | 1.2E-16 |
| 31 | si:dkey-280e21.3 | 12.76008 | 4.117484093 | 3.21E-17 | 1.81E-16 |
| 32 | mpp6b | 12.7557 | 3.339702368 | 3.2E-17 | 1.81E-16 |
| 33 | atp6v0cb | 12.62214 | 2.675063848 | 4.51E-17 | 2.53E-16 |
| 34 | tnc | 12.59912 | 3.754549742 | 5.15E-17 | 2.89E-16 |
| 35 | tfap2a | 12.58726 | 5.362655163 | 5.55E-17 | 3.11E-16 |
| 36 | palm1b | 12.42765 | 4.401007175 | 8.94E-17 | 4.97E-16 |
| 37 | slc32a1 | 12.21835 | 5.592228413 | 1.7E-16 | 9.39E-16 |
| 38 | isl1 | 11.40429 | 5.900708675 | 2.12E-15 | 1.13E-14 |
| 39 | cdh18a | 11.33207 | 4.791256905 | 2.65E-15 | 1.4E-14 |
| 40 | gnao1b | 11.25566 | 5.799740791 | 3.4E-15 | 1.8E-14 |
| 41 | st8sia5 | 11.23325 | 5.268324375 | 3.63E-15 | 1.92E-14 |
| 42 | ism1 | 11.17648 | 5.510960102 | 4.38E-15 | 2.3E-14 |
| 43 | gpm6aa | 10.93974 | 1.344212174 | 6.85E-15 | 3.57E-14 |
| 44 | vamp2 | 10.87052 | 2.421702147 | 1.06E-14 | 5.49E-14 |
| 45 | mfge8b | 10.78611 | 5.282325745 | 1.52E-14 | 7.84E-14 |
| 46 | pcbp3 | 10.68497 | 5.640929222 | 2.11E-14 | 1.08E-13 |
| 47 | nova2 | 10.68154 | 2.111906767 | 1.87E-14 | 9.58E-14 |
| 48 | gldn | 10.64083 | 6.705518723 | 2.45E-14 | 1.25E-13 |
| 49 | zgc:92912 | 10.52051 | 6.79508543 | 3.62E-14 | 1.84E-13 |
| 50 | stxbp1a | 10.50695 | 4.687234402 | 3.73E-14 | 1.89E-13 |
| 51 | ctnna2 | 10.33344 | 3.887794495 | 6.55E-14 | 3.29E-13 |
| 52 | h2afy2 | 10.27837 | 1.863886237 | 7.08E-14 | 3.54E-13 |
| 53 | ywhag2 | 10.23504 | 3.325377226 | 8.79E-14 | 4.38E-13 |
| 54 | gng3 | 10.04401 | 2.85466671 | 1.61E-13 | 7.93E-13 |
| 55 | hmx4 | 9.906478 | 3.542741299 | 2.68E-13 | 1.31E-12 |
| 56 | kif1aa | 9.864943 | 3.86991787 | 3.09E-13 | 1.51E-12 |
| 57 | ulk2 | 9.744619 | 3.450796127 | 4.6E-13 | 2.23E-12 |
| 58 | myt1b | 9.740718 | 4.443648338 | 4.7E-13 | 2.28E-12 |
| 59 | si:dkeyp-117h8.2 | 9.73509 | 4.334377289 | 4.8E-13 | 2.33E-12 |
| 60 | foxn3 | 9.665981 | 4.806186199 | 6.08E-13 | 2.94E-12 |
| 61 | atp2b3a | 9.658111 | 6.577282429 | 6.29E-13 | 3.03E-12 |
| 62 | rnasekb | 9.606361 | 1.709561229 | 6.64E-13 | 3.2E-12 |
| 63 | gnai2b | 9.586996 | 3.220162868 | 7.73E-13 | 3.72E-12 |
| 64 | rgs11 | 9.583486 | 6.444658279 | 8.08E-13 | 3.89E-12 |
| 65 | zgc:101840 | 9.544594 | 4.54749918 | 9.12E-13 | 4.37E-12 |
| 66 | si:dkey-81l17.6 | 9.395407 | 3.981873035 | 1.5E-12 | 7.14E-12 |
| 67 | atp6v1e1b | 9.331401 | 2.304917336 | 1.8E-12 | 8.5E-12 |
| 68 | rab3c | 9.250629 | 5.717923164 | 2.49E-12 | 1.17E-11 |
| 69 | eno2 | 9.198894 | 2.995109081 | 2.88E-12 | 1.35E-11 |
| 70 | ywhah | 9.113813 | 1.685589314 | 3.54E-12 | 1.65E-11 |
| 71 | lrrtm2 | 8.945631 | 5.791002274 | 7.08E-12 | 3.27E-11 |
| 72 | apof | 8.906734 | 7.841936588 | 8.12E-12 | 3.73E-11 |
| 73 | grm6b | 8.88651 | 6.39332819 | 8.69E-12 | 4E-11 |
| 74 | zfhx4 | 8.799727 | 3.47836566 | 1.15E-11 | 5.25E-11 |
| 75 | pcdh7b | 8.759848 | 4.50142765 | 1.34E-11 | 6.1E-11 |
| 76 | tfap2b | 8.524959 | 4.162265301 | 2.99E-11 | 1.35E-10 |
| 77 | si:ch211-242b18.1 | 8.523545 | 4.035508633 | 3.03E-11 | 1.36E-10 |
| 78 | khdrbs1b | 8.509533 | 2.889555931 | 3.14E-11 | 1.41E-10 |
| 79 | rassf4 | 8.509409 | 7.211473942 | 3.21E-11 | 1.44E-10 |
| 80 | atp1a3a | 8.389304 | 3.403484583 | 4.78E-11 | 2.13E-10 |
| 81 | si:dkey-7j14.5 | 8.371306 | 4.974750042 | 5.17E-11 | 2.3E-10 |
| 82 | tmem59l | 8.299499 | 3.087040901 | 6.53E-11 | 2.89E-10 |
| 83 | rhocb | 8.290924 | 2.350177288 | 6.65E-11 | 2.94E-10 |
| 84 | zc4h2 | 8.25281 | 2.310898542 | 7.51E-11 | 3.32E-10 |
| 85 | ctnnbip1 | 8.197758 | 2.358332396 | 9.22E-11 | 4.04E-10 |
| 86 | nhsl2 | 8.10646 | 3.857314825 | 1.3E-10 | 5.67E-10 |
| 87 | rgs9bp | 8.101274 | 9.685951233 | 1.34E-10 | 5.84E-10 |
| 88 | slc6a1b | 8.053 | 5.351060867 | 1.58E-10 | 6.86E-10 |
| 89 | pcp4l1 | 8.019696 | 5.120006561 | 1.77E-10 | 7.68E-10 |
| 90 | gad1b | 7.989145 | 6.296114445 | 1.98E-10 | 8.58E-10 |
| 91 | sox13 | 7.907565 | 3.139322281 | 2.61E-10 | 1.12E-09 |
| 92 | atpv0e2 | 7.879141 | 1.909164071 | 2.79E-10 | 1.2E-09 |
| 93 | calm1b | 7.632143 | 2.284138918 | 6.81E-10 | 2.87E-09 |
| 94 | mef2d | 7.547438 | 2.751391172 | 9.3E-10 | 3.89E-09 |
| 95 | nptnb | 7.408628 | 4.454635143 | 1.54E-09 | 6.37E-09 |
| 96 | gad2 | 7.342015 | 4.798160553 | 1.95E-09 | 8.01E-09 |
| 97 | tpd52l1 | 7.319592 | 3.633606195 | 2.1E-09 | 8.62E-09 |
| 98 | gpr85 | 7.270275 | 2.846331596 | 2.5E-09 | 1.02E-08 |
| 99 | mid1ip1l | 7.25369 | 2.444539547 | 2.63E-09 | 1.07E-08 |

**Cluster 15**

| | names | scores | logfoldchanges | pvals | pvals_adj |
|---|---|---|---|---|---|
| 0 | ptmab | 11.40276 | 1.48768127 | 2.82E-14 | 1.42E-13 |
| 1 | samsn1a | 11.34755 | 7.161873817 | 4.44E-14 | 2.23E-13 |
| 2 | nova2 | 11.23821 | 2.838591814 | 5.56E-14 | 2.78E-13 |
| 3 | rs1a | 10.40148 | 5.327133179 | 6.04E-13 | 2.91E-12 |
| 4 | scrt2 | 10.1512 | 4.016829491 | 1.21E-12 | 5.79E-12 |
| 5 | si:ch73-1a9.3 | 9.9276 | 1.661365151 | 2.04E-12 | 9.62E-12 |
| 6 | gnb3a | 9.224526 | 8.706531525 | 1.88E-11 | 8.58E-11 |
| 7 | mdkb | 8.979985 | 1.9459337 | 3.52E-11 | 1.59E-10 |
| 8 | marcksl1b | 8.880391 | 1.31265378 | 4.23E-11 | 1.9E-10 |
| 9 | gng13b | 8.705363 | 5.436635494 | 8.92E-11 | 3.95E-10 |
| 10 | tmem178b | 8.529777 | 4.66374445 | 1.52E-10 | 6.68E-10 |
| 11 | hmgn6 | 8.497252 | 1.350292802 | 1.52E-10 | 6.66E-10 |
| 12 | crx | 8.285783 | 2.663530827 | 3.12E-10 | 1.35E-09 |
| 13 | vsx1 | 7.944711 | 7.747238159 | 9.39E-10 | 3.97E-09 |
| 14 | myt1b | 7.572746 | 4.16663599 | 3E-09 | 1.24E-08 |
| 15 | neurod4 | 7.407536 | 2.815642834 | 4.97E-09 | 2.03E-08 |
| 16 | prox1a | 7.071976 | 2.751247168 | 1.47E-08 | 5.87E-08 |
| 17 | golga7ba | 6.983426 | 2.397141218 | 1.93E-08 | 7.67E-08 |
| 18 | nsg2 | 6.878228 | 2.516883135 | 2.73E-08 | 1.08E-07 |
| 19 | h3f3c | 6.7116 | 1.350272298 | 4.52E-08 | 1.76E-07 |
| 20 | ddah2 | 6.698836 | 2.308382034 | 4.86E-08 | 1.89E-07 |
| 21 | ndrg4 | 6.697749 | 1.982662439 | 4.83E-08 | 1.88E-07 |
| 22 | pdcl | 6.658487 | 1.699535966 | 5.47E-08 | 2.12E-07 |
| 23 | atp1a3a | 6.653254 | 3.195421457 | 5.66E-08 | 2.19E-07 |
| 24 | ppp1r14c | 6.585759 | 3.484622717 | 7.05E-08 | 2.73E-07 |
| 25 | syt5b | 6.347886 | 3.987569571 | 1.53E-07 | 5.79E-07 |
| 26 | rorab | 6.300248 | 3.410335302 | 1.78E-07 | 6.74E-07 |
| 27 | scg3 | 6.198381 | 2.738587856 | 2.46E-07 | 9.21E-07 |
| 28 | smarce1 | 6.154704 | 1.374952912 | 2.78E-07 | 1.04E-06 |
| 29 | lhx4 | 6.120314 | 4.274254322 | 3.19E-07 | 1.19E-06 |
| 30 | eml1 | 6.114842 | 2.521653652 | 3.22E-07 | 1.2E-06 |
| 31 | otx5 | 6.047013 | 2.319761515 | 3.98E-07 | 1.47E-06 |
| 32 | gpm6aa | 6.028522 | 1.104940295 | 4.14E-07 | 1.52E-06 |
| 33 | atp1b2a | 5.843028 | 2.165402174 | 7.78E-07 | 2.83E-06 |
| 34 | mpp6b | 5.742248 | 2.704507351 | 1.08E-06 | 3.89E-06 |
| 35 | zc4h2 | 5.720346 | 2.033675671 | 1.15E-06 | 4.13E-06 |
| 36 | ppp1r1b | 5.569129 | 4.717473984 | 1.9E-06 | 6.73E-06 |
| 37 | elovl4a | 5.536537 | 2.340507269 | 2.1E-06 | 7.41E-06 |
| 38 | hnrnpa0a | 5.476362 | 1.125402689 | 2.5E-06 | 8.82E-06 |
| 39 | vamp1 | 5.439999 | 6.211827278 | 2.89E-06 | 1.01E-05 |
| 40 | sypb | 5.40134 | 2.617710114 | 3.24E-06 | 1.13E-05 |
| 41 | mid1ip1b | 5.35729 | 0.844224215 | 3.64E-06 | 1.26E-05 |
| 42 | gfra1a | 5.3465 | 3.23440671 | 3.89E-06 | 1.35E-05 |
| 43 | pcp4l1 | 5.155189 | 4.338974953 | 7.2E-06 | 2.46E-05 |
| 44 | h3f3b.1-2 | 5.141631 | 0.962968528 | 7.33E-06 | 2.5E-05 |
| 45 | fam107b | 5.023374 | 2.349857092 | 1.09E-05 | 3.68E-05 |
| 46 | bcl2l10 | 4.945974 | 1.558780789 | 1.39E-05 | 4.63E-05 |
| 47 | h3f3b.1-1 | 4.919767 | 0.868024945 | 1.49E-05 | 4.95E-05 |
| 48 | ndrg1b | 4.905591 | 2.516608238 | 1.59E-05 | 5.28E-05 |
| 49 | sox4a-1 | 4.899141 | 2.36222291 | 1.62E-05 | 5.39E-05 |
| 50 | ckbb | 4.874777 | 0.909866691 | 1.71E-05 | 5.69E-05 |
| 51 | celf3a | 4.851412 | 2.779972315 | 1.89E-05 | 6.26E-05 |
| 52 | si:ch211-260e23.9 | 4.798407 | 2.211287498 | 2.24E-05 | 7.31E-05 |
| 53 | isl1 | 4.767258 | 4.190191269 | 2.47E-05 | 8.09E-05 |
| 54 | jpt1b | 4.678518 | 0.93409586 | 3.21E-05 | 0.000104 |
| 55 | stx3a | 4.674256 | 2.516481876 | 3.31E-05 | 0.000108 |
| 56 | sv2bb | 4.625063 | 4.513708115 | 3.88E-05 | 0.000124 |
| 57 | snap25b | 4.59484 | 1.978399158 | 4.23E-05 | 0.000136 |
| 58 | si:dkey-253i9.4 | 4.580679 | 3.151838541 | 4.45E-05 | 0.000142 |
| 59 | hmgb1b | 4.519246 | 0.773526788 | 5.26E-05 | 0.000168 |
| 60 | lmo4b | 4.490333 | 3.536035776 | 5.9E-05 | 0.000188 |
| 61 | mbd3b | 4.440171 | 2.648721218 | 6.89E-05 | 0.000217 |
| 62 | ubl3b | 4.43759 | 2.39005971 | 6.94E-05 | 0.000218 |
| 63 | h2afx1 | 4.414063 | 0.966691911 | 7.39E-05 | 0.000232 |
| 64 | sox12 | 4.368863 | 3.882088423 | 8.61E-05 | 0.000269 |
| 65 | chn1 | 4.362575 | 2.21430254 | 8.75E-05 | 0.000274 |
| 66 | grb2b | 4.324765 | 1.536647797 | 9.8E-05 | 0.000306 |
| 67 | tnnt1 | 4.290765 | 6.435362339 | 0.00011 | 0.000338 |
| 68 | zgc:77784 | 4.252775 | 4.691111565 | 0.000123 | 0.000379 |
| 69 | hnrnpaba | 4.244096 | 0.570516288 | 0.000124 | 0.00038 |
| 70 | ptmaa | 4.234839 | 0.689352453 | 0.000127 | 0.000389 |
| 71 | igsf21b | 4.203497 | 4.630031109 | 0.000143 | 0.000438 |
| 72 | ndrg3b | 4.196335 | 1.725932717 | 0.000146 | 0.000446 |
| 73 | hmgb3a | 4.191482 | 0.899790466 | 0.000147 | 0.00045 |
| 74 | si:dkey-56m19.5 | 4.142285 | 1.28576386 | 0.000171 | 0.000523 |
| 75 | tp53inp1 | 4.141752 | 1.42013967 | 0.000172 | 0.000525 |
| 76 | ctbp2a | 4.138351 | 2.247342825 | 0.000174 | 0.000532 |
| 77 | pcdh19 | 4.138004 | 3.2908144 | 0.000175 | 0.000533 |
| 78 | itm2ca | 4.115719 | 2.781083822 | 0.000187 | 0.000563 |
| 79 | sesn1 | 4.09314 | 1.534160018 | 0.000199 | 0.000599 |
| 80 | clstn3 | 4.08592 | 3.937090874 | 0.000205 | 0.000616 |
| 81 | ucp2 | 4.075579 | 1.194929481 | 0.00021 | 0.000632 |
| 82 | si:ch211-140m22.7 | 4.022749 | 1.520209432 | 0.000247 | 0.000739 |
| 83 | jagn1a | 4.017736 | 3.121504307 | 0.000252 | 0.000753 |
| 84 | selenow1 | 3.996627 | 0.988461673 | 0.000266 | 0.000796 |
| 85 | dab1b | 3.968604 | 3.615131855 | 0.000292 | 0.000871 |
| 86 | h2afva | 3.961214 | 0.857348919 | 0.000295 | 0.000881 |
| 87 | khdrbs1a | 3.930096 | 0.570990384 | 0.000323 | 0.000948 |
| 88 | bhlhe23 | 3.922093 | 5.273126602 | 0.000336 | 0.000986 |
| 89 | ube2e1 | 3.86386 | 1.190570354 | 0.000397 | 0.001162 |
| 90 | oaz1b | 3.858248 | 0.776799142 | 0.000403 | 0.001178 |
| 91 | ssbp3b | 3.842726 | 1.742057443 | 0.000424 | 0.001238 |
| 92 | si:dkey-28b4.7 | 3.83905 | 1.349460602 | 0.000428 | 0.00125 |
| 93 | ppp1r9bb | 3.836051 | 4.274051666 | 0.000434 | 0.001266 |
| 94 | si:ch211-242b18.1 | 3.791735 | 2.402193546 | 0.000494 | 0.001437 |
| 95 | snx8b | 3.790955 | 7.411890984 | 0.000496 | 0.001443 |
| 96 | cpe | 3.789239 | 2.022012949 | 0.000497 | 0.001446 |
| 97 | cd9a | 3.784642 | 2.481589556 | 0.000504 | 0.001466 |
| 98 | calm3a | 3.760697 | 1.569121122 | 0.000542 | 0.001545 |
| 99 | si:ch211-222l21.1 | 3.760572 | 0.78755343 | 0.000533 | 0.001523 |
