## Supplementary_file_2 for "Single cell RNA sequencing unravels the transcriptional network underlying zebrafish retina regeneration"

**Differentially expressed genes between Non-reactive Müller glia (1) and Non-reactive Müller glia (2) clusters**

| | group | names | scores | logfoldchanges | pvals | pvals_adj |
|---|---|---|---|---|---|---|
| 0 | Non-reactive Müller glia (1) | inhbaa | 37.94407 | 3.546254158 | 7.9E-170 | 5.5E-166 |
| 1 | Non-reactive Müller glia (1) | bambia | 32.67387 | 4.581230164 | 5.7E-193 | 7.9E-189 |
| 2 | Non-reactive Müller glia (1) | gpm6ab | 31.56866 | 2.196494102 | 8.8E-117 | 1.7E-113 |
| 3 | Non-reactive Müller glia (1) | tbx2a | 31.52823 | 5.262084484 | 2.9E-189 | 2.7E-185 |
| 4 | Non-reactive Müller glia (1) | crabp2b | 30.49274 | 3.225553989 | 2.5E-129 | 7E-126 |
| 5 | Non-reactive Müller glia (1) | pdgfrl | 28.89457 | 4.525546551 | 1.1E-151 | 5.2E-148 |
| 6 | Non-reactive Müller glia (1) | lman1 | 28.74099 | 4.747096062 | 1.3E-160 | 7E-157 |
| 7 | Non-reactive Müller glia (1) | icn | 27.91342 | 2.361161947 | 1.3E-97 | 2.26E-94 |
| 8 | Non-reactive Müller glia (1) | hspb8 | 27.43563 | 3.687711716 | 4.5E-126 | 1E-122 |
| 9 | Non-reactive Müller glia (1) | cyp26c1 | 27.23067 | 7.758151054 | 7.3E-149 | 2.9E-145 |
| 10 | Non-reactive Müller glia (1) | efnb2a | 27.03174 | 4.671700478 | 4.7E-138 | 1.4E-134 |
| 11 | Non-reactive Müller glia (1) | prss35 | 26.8124 | 5.635470867 | 3.9E-144 | 1.3E-140 |
| 12 | Non-reactive Müller glia (1) | aqp1a.1 | 26.73771 | 1.800045252 | 4.14E-92 | 6.03E-89 |
| 13 | Non-reactive Müller glia (1) | sncgb | 23.87616 | 3.018136978 | 7.34E-97 | 1.2E-93 |
| 14 | Non-reactive Müller glia (1) | si:dkey-222f2.1 | 23.37817 | 3.701656818 | 7.4E-104 | 1.4E-100 |
| 15 | Non-reactive Müller glia (1) | ndrg4 | 22.72685 | 2.267275572 | 4.5E-81 | 6.24E-78 |
| 16 | Non-reactive Müller glia (1) | atp1b4 | 21.74528 | 1.651861191 | 4.84E-75 | 6.38E-72 |
| 17 | Non-reactive Müller glia (1) | CR383676.1 | 20.0594 | 0.50164026 | 1.4E-65 | 1.62E-62 |
| 18 | Non-reactive Müller glia (1) | zgc:165604 | 19.54804 | 1.182065964 | 7.19E-61 | 7.66E-58 |
| 19 | Non-reactive Müller glia (1) | fstb | 19.43203 | 3.406087399 | 9.9E-75 | 1.25E-71 |
| 20 | Non-reactive Müller glia (1) | myl6 | 18.76356 | 2.052682638 | 4.71E-62 | 5.22E-59 |
| 21 | Non-reactive Müller glia (1) | ctsla | 18.67946 | 1.334076166 | 1.13E-57 | 1.04E-54 |
| 22 | Non-reactive Müller glia (1) | fgf24 | 18.24792 | 2.191919565 | 2.05E-58 | 1.96E-55 |
| 23 | Non-reactive Müller glia (1) | txn | 18.07088 | 3.711958408 | 1.15E-68 | 1.38E-65 |
| 24 | Non-reactive Müller glia (1) | myl9a | 17.80181 | 2.873800278 | 1.2E-60 | 1.23E-57 |
| 25 | Non-reactive Müller glia (1) | spock3 | 17.48943 | 1.053786159 | 2.74E-52 | 2.17E-49 |
| 26 | Non-reactive Müller glia (1) | uchl1 | 17.40828 | 2.283057451 | 2.13E-55 | 1.84E-52 |
| 27 | Non-reactive Müller glia (1) | atp1b1a | 17.10874 | 1.693768382 | 2.22E-51 | 1.62E-48 |
| 28 | Non-reactive Müller glia (1) | syt11a | 16.73093 | 2.431065798 | 1.41E-53 | 1.18E-50 |
| 29 | Non-reactive Müller glia (1) | tbx2b | 16.37122 | 4.085027695 | 2.29E-56 | 2.04E-53 |
| 30 | Non-reactive Müller glia (1) | gpm6aa | 16.28531 | 1.206821322 | 1.26E-46 | 7.72E-44 |
| 31 | Non-reactive Müller glia (1) | slco1c1 | 16.11231 | 2.497951508 | 1.37E-49 | 9.5E-47 |
| 32 | Non-reactive Müller glia (1) | CR936442.1 | 16.08533 | 2.014392614 | 1.75E-47 | 1.18E-44 |
| 33 | Non-reactive Müller glia (1) | gapdhs | 15.97316 | 0.750086546 | 1.51E-45 | 9.12E-43 |
| 34 | Non-reactive Müller glia (1) | si:ch211-237l4.6 | 15.72556 | 3.772512913 | 1.66E-51 | 1.24E-48 |
| 35 | Non-reactive Müller glia (1) | serpinb1 | 15.5371 | 2.348143816 | 6.67E-47 | 4.3E-44 |
| 36 | Non-reactive Müller glia (1) | wfdc2 | 15.52593 | 1.45033884 | 5.97E-44 | 3.45E-41 |
| 37 | Non-reactive Müller glia (1) | c1qtnf12 | 15.51172 | 5.112378597 | 2.51E-52 | 2.05E-49 |
| 38 | Non-reactive Müller glia (1) | tcima | 15.4024 | 2.973572016 | 2.65E-50 | 1.88E-47 |
| 39 | Non-reactive Müller glia (1) | cavin1b | 15.34797 | 5.277170658 | 1.66E-51 | 1.24E-48 |
| 40 | Non-reactive Müller glia (1) | fabp7a | 15.11265 | 1.322055101 | 6.44E-42 | 3.57E-39 |
| 41 | Non-reactive Müller glia (1) | cadm4 | 14.82402 | 1.476937056 | 7.87E-41 | 4.11E-38 |
| 42 | Non-reactive Müller glia (1) | rasgef1bb | 14.75837 | 1.830831051 | 7.32E-42 | 3.97E-39 |
| 43 | Non-reactive Müller glia (1) | zgc:77112 | 14.5573 | 25.50667953 | 1.16E-46 | 7.32E-44 |
| 44 | Non-reactive Müller glia (1) | ckbb | 14.55208 | 0.709458292 | 1.05E-39 | 5.18E-37 |
| 45 | Non-reactive Müller glia (1) | sc5d | 14.4588 | 1.554155469 | 3.29E-40 | 1.66E-37 |
| 46 | Non-reactive Müller glia (1) | si:ch211-207i1.2 | 14.44481 | 2.294787169 | 8.25E-42 | 4.39E-39 |
| 47 | Non-reactive Müller glia (1) | elovl1b | 14.40426 | 3.347476721 | 4.6E-44 | 2.71E-41 |
| 48 | Non-reactive Müller glia (1) | hmgcra | 14.38713 | 1.501916409 | 1.35E-39 | 6.45E-37 |
| 49 | Non-reactive Müller glia (1) | dbn1 | 14.24266 | 2.300209045 | 1.19E-40 | 6.09E-38 |
| 50 | Non-reactive Müller glia (1) | cyp26a1 | 14.20365 | 2.129999638 | 1.3E-39 | 6.31E-37 |
| 51 | Non-reactive Müller glia (1) | camk1db | 14.13887 | 1.693755865 | 1.92E-38 | 8.73E-36 |
| 52 | Non-reactive Müller glia (1) | ucmab | 14.02546 | 25.69620132 | 1.59E-43 | 8.96E-41 |
| 53 | Non-reactive Müller glia (1) | si:dkey-56f14.7 | 13.818 | 1.158595204 | 2.97E-36 | 1.21E-33 |
| 54 | Non-reactive Müller glia (1) | clu | 13.7356 | 2.478725433 | 3.69E-38 | 1.6E-35 |
| 55 | Non-reactive Müller glia (1) | mmp2 | 13.58922 | 3.158926725 | 4.99E-39 | 2.3E-36 |
| 56 | Non-reactive Müller glia (1) | qsox1 | 13.52142 | 2.430235147 | 1.99E-37 | 8.34E-35 |
| 57 | Non-reactive Müller glia (1) | lepb | 13.42554 | 3.682810307 | 2.81E-39 | 1.32E-36 |
| 58 | Non-reactive Müller glia (1) | tbx4 | 13.13469 | 4.719081879 | 3.24E-38 | 1.42E-35 |
| 59 | Non-reactive Müller glia (1) | foxg1d | 13.11193 | 25.21665001 | 2.2E-38 | 9.84E-36 |
| 60 | Non-reactive Müller glia (1) | AL954322.2 | 13.0667 | 3.372153521 | 3.08E-36 | 1.23E-33 |
| 61 | Non-reactive Müller glia (1) | gsna | 13.05477 | 2.201474905 | 8.18E-35 | 3.06E-32 |
| 62 | Non-reactive Müller glia (1) | lgals2a | 13.01243 | 5.081232548 | 7.4E-38 | 3.15E-35 |
| 63 | Non-reactive Müller glia (1) | tbx3a | 12.92272 | 3.67233181 | 2.24E-36 | 9.27E-34 |
| 64 | Non-reactive Müller glia (1) | sema3fa | 12.90078 | 1.486633539 | 6.08E-33 | 2.13E-30 |
| 65 | Non-reactive Müller glia (1) | hdac5 | 12.87001 | 3.191675663 | 2.14E-35 | 8.34E-33 |
| 66 | Non-reactive Müller glia (1) | cahz | 12.84609 | 0.660078168 | 7.78E-33 | 2.69E-30 |
| 67 | Non-reactive Müller glia (1) | mrap2a | 12.62787 | 24.92942238 | 8.76E-36 | 3.46E-33 |
| 68 | Non-reactive Müller glia (1) | msmo1 | 12.62267 | 1.18071568 | 1.12E-31 | 3.61E-29 |
| 69 | Non-reactive Müller glia (1) | eno1a | 12.50841 | 0.595672131 | 6.43E-31 | 2.02E-28 |
| 70 | Non-reactive Müller glia (1) | stm | 12.48074 | 26.77745628 | 5.19E-35 | 2E-32 |
| 71 | Non-reactive Müller glia (1) | s100u | 12.41863 | 25.10116196 | 1.09E-34 | 4.04E-32 |
| 72 | Non-reactive Müller glia (1) | hspbp1 | 12.40152 | 2.173260689 | 3.61E-32 | 1.22E-29 |
| 73 | Non-reactive Müller glia (1) | hsd17b12b | 12.34388 | 1.791157246 | 3.57E-31 | 1.14E-28 |
| 74 | Non-reactive Müller glia (1) | map4k2 | 12.31199 | 3.055575371 | 9.93E-33 | 3.4E-30 |
| 75 | Non-reactive Müller glia (1) | rhoab | 12.29561 | 1.0069803 | 4.85E-30 | 1.43E-27 |
| 76 | Non-reactive Müller glia (1) | ccdc85a | 12.20654 | 1.922718763 | 1.1E-30 | 3.36E-28 |
| 77 | Non-reactive Müller glia (1) | clcf1 | 12.19888 | 25.50375175 | 1.49E-33 | 5.42E-31 |
| 78 | Non-reactive Müller glia (1) | igsf9ba | 12.18975 | 1.403206944 | 5.23E-30 | 1.52E-27 |
| 79 | Non-reactive Müller glia (1) | anxa2a | 12.17908 | 4.538850784 | 1.81E-33 | 6.5E-31 |
| 80 | Non-reactive Müller glia (1) | si:ch211-79k12.1 | 12.10606 | 24.98733521 | 4.42E-33 | 1.57E-30 |
| 81 | Non-reactive Müller glia (1) | UBB | 12.04449 | 1.197988391 | 5.21E-29 | 1.42E-26 |
| 82 | Non-reactive Müller glia (1) | lmo7a | 11.9385 | 2.868408918 | 1.62E-30 | 4.82E-28 |
| 83 | Non-reactive Müller glia (1) | cyp51 | 11.84347 | 1.309940219 | 1.31E-28 | 3.4E-26 |
| 84 | Non-reactive Müller glia (1) | chst15 | 11.83229 | 1.787382841 | 5.68E-29 | 1.53E-26 |
| 85 | Non-reactive Müller glia (1) | idh1 | 11.73837 | 1.821365118 | 1.33E-28 | 3.4E-26 |
| 86 | Non-reactive Müller glia (1) | cxcl14 | 11.67784 | 2.674033642 | 1.29E-29 | 3.63E-27 |
| 87 | Non-reactive Müller glia (1) | rhbg | 11.65844 | 0.776906848 | 2.69E-27 | 6.48E-25 |
| 88 | Non-reactive Müller glia (1) | calcr | 11.65833 | 24.70619011 | 7.63E-31 | 2.37E-28 |
| 89 | Non-reactive Müller glia (1) | tnfsf12 | 11.6406 | 24.77231407 | 9.32E-31 | 2.87E-28 |
| 90 | Non-reactive Müller glia (1) | zic2b | 11.58307 | 2.881770611 | 1.16E-29 | 3.31E-27 |
| 91 | Non-reactive Müller glia (1) | si:zfos-943e10.1 | 11.55017 | 1.356756806 | 1.45E-27 | 3.52E-25 |
| 92 | Non-reactive Müller glia (1) | hbegfa | 11.53205 | 1.847887278 | 5.12E-28 | 1.26E-25 |
| 93 | Non-reactive Müller glia (1) | mlip | 11.47244 | 24.66289902 | 6.14E-30 | 1.77E-27 |
| 94 | Non-reactive Müller glia (1) | kctd12.2 | 11.41919 | 2.337012529 | 4.67E-28 | 1.18E-25 |
| 95 | Non-reactive Müller glia (1) | si:ch73-31d8.2 | 11.40491 | 1.436942935 | 7.13E-27 | 1.67E-24 |
| 96 | Non-reactive Müller glia (1) | ppdpfb | 11.4041 | 0.890761256 | 2.22E-26 | 4.95E-24 |
| 97 | Non-reactive Müller glia (1) | nupr1a | 11.38511 | 1.673977971 | 3.88E-27 | 9.18E-25 |
| 98 | Non-reactive Müller glia (1) | cd9a | 11.3705 | 24.68388748 | 1.9E-29 | 5.33E-27 |
| 99 | Non-reactive Müller glia (1) | dbi | 11.35781 | 0.97066009 | 1.85E-26 | 4.23E-24 |

| | group | names | scores | logfoldchanges | pvals | pvals_adj |
|---|---|---|---|---|---|---|
| 100 | Non-reactive Müller glia (2) | apoeb | 43.90213 | 1.884748697 | 7.6E-209 | 2.1E-204 |
| 101 | Non-reactive Müller glia (2) | chrdl2 | 37.53057 | 4.166360855 | 6.4E-128 | 1.6E-124 |
| 102 | Non-reactive Müller glia (2) | rdh10a | 35.07629 | 3.738123655 | 2.8E-120 | 5.9E-117 |
| 103 | Non-reactive Müller glia (2) | aldh1a3 | 28.35645 | 3.093188524 | 2.06E-96 | 3.17E-93 |
| 104 | Non-reactive Müller glia (2) | mstnb | 19.48931 | 2.769267321 | 1.1E-58 | 1.09E-55 |
| 105 | Non-reactive Müller glia (2) | sncga | 16.32416 | 1.095173955 | 2.13E-47 | 1.4E-44 |
| 106 | Non-reactive Müller glia (2) | stc2a | 13.51636 | 1.607256413 | 5.67E-35 | 2.15E-32 |
| 107 | Non-reactive Müller glia (2) | dhrs3a | 13.08953 | 4.178340435 | 4.16E-32 | 1.39E-29 |
| 108 | Non-reactive Müller glia (2) | smoc1 | 13.02685 | 7.674436092 | 8.79E-32 | 2.86E-29 |
| 109 | Non-reactive Müller glia (2) | si:dkey-164f24.2 | 12.8342 | 1.102459073 | 5.13E-32 | 1.69E-29 |
| 110 | Non-reactive Müller glia (2) | nr2f5 | 12.69152 | 4.196836948 | 1.39E-30 | 4.18E-28 |
| 111 | Non-reactive Müller glia (2) | si:ch211-222l21.1 | 10.97201 | 1.698405981 | 1.74E-24 | 3.48E-22 |
| 112 | Non-reactive Müller glia (2) | rgmb | 10.92153 | 2.732622147 | 3.85E-24 | 7.41E-22 |
| 113 | Non-reactive Müller glia (2) | gnai2b | 10.88741 | 1.384487748 | 2.81E-24 | 5.44E-22 |
| 114 | Non-reactive Müller glia (2) | ackr3b | 10.57193 | 4.320560932 | 7.98E-23 | 1.41E-20 |
| 115 | Non-reactive Müller glia (2) | rgmd | 10.23703 | 2.516204357 | 9.92E-22 | 1.61E-19 |
| 116 | Non-reactive Müller glia (2) | cyp1d1 | 9.944285 | 3.030125618 | 1.05E-20 | 1.53E-18 |
| 117 | Non-reactive Müller glia (2) | hspb1 | 9.882424 | 1.128902912 | 4.71E-21 | 7.08E-19 |
| 118 | Non-reactive Müller glia (2) | mt2 | 9.792654 | 0.823626876 | 1.24E-20 | 1.76E-18 |
| 119 | Non-reactive Müller glia (2) | elmod1 | 8.538007 | 4.446221352 | 4.39E-16 | 4.01E-14 |
| 120 | Non-reactive Müller glia (2) | fosb | 8.491188 | 0.434918135 | 2.65E-16 | 2.5E-14 |
| 121 | Non-reactive Müller glia (2) | tnfaip2a | 8.352043 | 1.937019706 | 1.42E-15 | 1.22E-13 |
| 122 | Non-reactive Müller glia (2) | bmpr1ba | 8.216331 | 1.899815202 | 3.65E-15 | 2.92E-13 |
| 123 | Non-reactive Müller glia (2) | egr1 | 8.163019 | 0.793830395 | 4.14E-15 | 3.28E-13 |
| 124 | Non-reactive Müller glia (2) | atp1b3a | 8.140251 | 0.503036618 | 4.29E-15 | 3.4E-13 |
| 125 | Non-reactive Müller glia (2) | cldn7a | 8.026628 | 4.161813259 | 1.58E-14 | 1.15E-12 |
| 126 | Non-reactive Müller glia (2) | ptgs2a | 7.635228 | 1.153928518 | 1.76E-13 | 1.16E-11 |
| 127 | Non-reactive Müller glia (2) | sparc | 7.574094 | 0.374150574 | 2.02E-13 | 1.32E-11 |
| 128 | Non-reactive Müller glia (2) | leo1 | 7.426731 | 0.889044166 | 7.11E-13 | 4.24E-11 |
| 129 | Non-reactive Müller glia (2) | pnp6 | 7.362914 | 1.012173533 | 1.08E-12 | 6.33E-11 |
| 130 | Non-reactive Müller glia (2) | ednraa | 7.147677 | 1.258870363 | 4.55E-12 | 2.42E-10 |
| 131 | Non-reactive Müller glia (2) | ephb2b | 7.121871 | 1.141223073 | 5.27E-12 | 2.77E-10 |
| 132 | Non-reactive Müller glia (2) | brinp3b | 7.115801 | 3.568098307 | 6.37E-12 | 3.32E-10 |
| 133 | Non-reactive Müller glia (2) | pip5k1bb | 6.962028 | 1.683396935 | 1.56E-11 | 7.71E-10 |
| 134 | Non-reactive Müller glia (2) | bmp7b | 6.868118 | 1.976043463 | 2.85E-11 | 1.36E-09 |
| 135 | Non-reactive Müller glia (2) | bmpr1bb | 6.73305 | 4.386248589 | 6.94E-11 | 3.18E-09 |
| 136 | Non-reactive Müller glia (2) | im:7152348 | 6.627189 | 0.705562592 | 1.08E-10 | 4.78E-09 |
| 137 | Non-reactive Müller glia (2) | sox3 | 6.623856 | 1.806533337 | 1.25E-10 | 5.44E-09 |
| 138 | Non-reactive Müller glia (2) | igfbp1a | 6.617581 | 0.780394793 | 1.13E-10 | 5E-09 |
| 139 | Non-reactive Müller glia (2) | CU467822.1 | 6.530359 | 0.394079208 | 1.88E-10 | 7.92E-09 |
| 140 | Non-reactive Müller glia (2) | six6a | 6.162166 | 2.477043152 | 1.94E-09 | 7.1E-08 |
| 141 | Non-reactive Müller glia (2) | cib2 | 6.148203 | 1.364170671 | 2.01E-09 | 7.33E-08 |
| 142 | Non-reactive Müller glia (2) | ndrg3a | 6.075061 | 0.556735098 | 2.75E-09 | 9.78E-08 |
| 143 | Non-reactive Müller glia (2) | pik3r3b | 6.064154 | 0.773879051 | 3.1E-09 | 1.09E-07 |
| 144 | Non-reactive Müller glia (2) | ponzr5 | 5.963062 | 2.685591936 | 6E-09 | 2.01E-07 |
| 145 | Non-reactive Müller glia (2) | irx5a | 5.777402 | 0.979610622 | 1.57E-08 | 4.87E-07 |
| 146 | Non-reactive Müller glia (2) | pcyt1bb | 5.747045 | 2.039861917 | 1.94E-08 | 5.92E-07 |
| 147 | Non-reactive Müller glia (2) | mt-co3 | 5.717512 | 0.266081601 | 1.97E-08 | 5.98E-07 |
| 148 | Non-reactive Müller glia (2) | slc16a9a | 5.494378 | 2.257709026 | 7.49E-08 | 2.05E-06 |
| 149 | Non-reactive Müller glia (2) | BX663503.3 | 5.427116 | 1.085737824 | 1.02E-07 | 2.71E-06 |
| 150 | Non-reactive Müller glia (2) | tob1a | 5.382612 | 0.387688249 | 1.21E-07 | 3.14E-06 |
| 151 | Non-reactive Müller glia (2) | foxi2 | 5.289058 | 3.5141325 | 2.18E-07 | 5.43E-06 |
| 152 | Non-reactive Müller glia (2) | dhrs13a.1 | 5.249379 | 0.997201025 | 2.55E-07 | 6.29E-06 |
| 153 | Non-reactive Müller glia (2) | abat | 5.124996 | 0.678323746 | 4.68E-07 | 1.09E-05 |
| 154 | Non-reactive Müller glia (2) | arpp21 | 5.071553 | 1.379363537 | 6.24E-07 | 1.4E-05 |
| 155 | Non-reactive Müller glia (2) | hdac4 | 5.067826 | 2.217663288 | 6.45E-07 | 1.44E-05 |
| 156 | Non-reactive Müller glia (2) | slc38a2 | 5.007825 | 1.5596807 | 8.59E-07 | 1.89E-05 |
| 157 | Non-reactive Müller glia (2) | CR354540.1 | 4.994704 | 1.773470879 | 9.17E-07 | 2E-05 |
| 158 | Non-reactive Müller glia (2) | vax1 | 4.981232 | 4.07425642 | 1E-06 | 2.15E-05 |
| 159 | Non-reactive Müller glia (2) | esr2a | 4.951549 | 3.530527353 | 1.15E-06 | 2.43E-05 |
| 160 | Non-reactive Müller glia (2) | serpinf1 | 4.927566 | 3.571918011 | 1.29E-06 | 2.71E-05 |
| 161 | Non-reactive Müller glia (2) | rrh | 4.925615 | 0.660763323 | 1.24E-06 | 2.62E-05 |
| 162 | Non-reactive Müller glia (2) | zgc:195173 | 4.882537 | 0.389116883 | 1.43E-06 | 2.99E-05 |
| 163 | Non-reactive Müller glia (2) | rcan3 | 4.87946 | 1.607069612 | 1.59E-06 | 3.28E-05 |
| 164 | Non-reactive Müller glia (2) | vegfaa | 4.649835 | 0.446214408 | 4.45E-06 | 8.29E-05 |
| 165 | Non-reactive Müller glia (2) | ch25h | 4.637689 | 2.428310871 | 4.96E-06 | 9.14E-05 |
| 166 | Non-reactive Müller glia (2) | cntnap5b | 4.634336 | 3.551418543 | 5.07E-06 | 9.32E-05 |
| 167 | Non-reactive Müller glia (2) | dusp2 | 4.613782 | 0.51304549 | 5.29E-06 | 9.64E-05 |
| 168 | Non-reactive Müller glia (2) | f3b | 4.606153 | 0.420251071 | 5.41E-06 | 9.84E-05 |
| 169 | Non-reactive Müller glia (2) | mycn | 4.597778 | 1.000431299 | 5.82E-06 | 0.000105 |
| 170 | Non-reactive Müller glia (2) | cav1 | 4.592935 | 0.461518347 | 5.84E-06 | 0.000106 |
| 171 | Non-reactive Müller glia (2) | ip6k2b | 4.542986 | 0.566893041 | 7.37E-06 | 0.000132 |
| 172 | Non-reactive Müller glia (2) | mt-co1 | 4.494689 | 0.257468611 | 8.99E-06 | 0.000154 |
| 173 | Non-reactive Müller glia (2) | rlbp1a | 4.31893 | 0.194846377 | 1.92E-05 | 0.000306 |
| 174 | Non-reactive Müller glia (2) | si:ch211-145b13.5 | 4.237026 | 1.026896715 | 2.85E-05 | 0.00043 |
| 175 | Non-reactive Müller glia (2) | atp2b3b | 4.216623 | 2.794859648 | 3.16E-05 | 0.000471 |
| 176 | Non-reactive Müller glia (2) | foxg1b | 4.211659 | 0.636181951 | 3.15E-05 | 0.000471 |
| 177 | Non-reactive Müller glia (2) | si:dkey-76d14.2 | 4.09479 | 0.947381318 | 5.17E-05 | 0.000722 |
| 178 | Non-reactive Müller glia (2) | hmgb2a | 4.073831 | 0.452234268 | 5.55E-05 | 0.00077 |
| 179 | Non-reactive Müller glia (2) | si:dkey-217l24.1 | 4.07129 | 1.555879354 | 5.73E-05 | 0.000794 |
| 180 | Non-reactive Müller glia (2) | bgnb | 4.055054 | 0.343163759 | 5.95E-05 | 0.000821 |
| 181 | Non-reactive Müller glia (2) | si:ch73-141c7.1 | 3.993604 | 1.651939869 | 7.88E-05 | 0.001035 |
| 182 | Non-reactive Müller glia (2) | fosab | 3.939539 | 0.197727963 | 9.23E-05 | 0.001193 |
| 183 | Non-reactive Müller glia (2) | si:ch73-46j18.5 | 3.937312 | 0.285565376 | 9.63E-05 | 0.001238 |
| 184 | Non-reactive Müller glia (2) | si:dkey-112a7.4 | 3.933747 | 0.551066101 | 9.86E-05 | 0.001264 |
| 185 | Non-reactive Müller glia (2) | NPFFR2 | 3.91013 | 1.037614346 | 0.000109 | 0.001361 |
| 186 | Non-reactive Müller glia (2) | iqca1 | 3.893461 | 3.934196949 | 0.000119 | 0.001466 |
| 187 | Non-reactive Müller glia (2) | diras1a | 3.871813 | 1.346796513 | 0.000128 | 0.001552 |
| 188 | Non-reactive Müller glia (2) | marcksl1a | 3.849621 | 0.363260418 | 0.000137 | 0.001654 |
| 189 | Non-reactive Müller glia (2) | slc3a2a | 3.825608 | 0.431124777 | 0.000151 | 0.001802 |
| 190 | Non-reactive Müller glia (2) | sik2b | 3.822247 | 0.30115059 | 0.000152 | 0.001814 |
| 191 | Non-reactive Müller glia (2) | cyp2p6 | 3.816729 | 0.986664295 | 0.000158 | 0.001874 |
| 192 | Non-reactive Müller glia (2) | si:ch211-105c13.3 | 3.791507 | 3.397749901 | 0.000176 | 0.002065 |
| 193 | Non-reactive Müller glia (2) | aspa | 3.764535 | 0.447333992 | 0.000192 | 0.002192 |
| 194 | Non-reactive Müller glia (2) | aldh1a2 | 3.729247 | 0.51929605 | 0.000218 | 0.002459 |
| 195 | Non-reactive Müller glia (2) | rxfp3.3b | 3.721139 | 6.246320248 | 0.000232 | 0.002594 |
| 196 | Non-reactive Müller glia (2) | nfasca | 3.671763 | 0.156083569 | 0.00027 | 0.002978 |
| 197 | Non-reactive Müller glia (2) | si:ch211-286o17.1 | 3.655677 | 2.171005011 | 0.000295 | 0.003226 |
| 198 | Non-reactive Müller glia (2) | sh3gl2a | 3.641486 | 1.672080874 | 0.000311 | 0.003309 |
| 199 | Non-reactive Müller glia (2) | aldh1l1 | 3.626436 | 1.54876101 | 0.000328 | 0.003472 |
