## Supplementary material for "Single cell RNA sequencing unravels the transcriptional network underlying zebrafish retina regeneration": Suppelementary_file_3

**Differentially expressed genes between Reactive Müller glia (2) and Reactive Müller glia (1) clusters**

| | group | names | scores | logfoldchanges | pvals | pvals_adj |
|---|---|---|---|---|---|---|
| 0 | Reactive Müller glia (2) | ckbb | 50.84286 | 3.734843969 | 0 | 0 |
| 1 | Reactive Müller glia (2) | acbd7 | 43.06075 | 3.538524628 | 1.7E-245 | 4.2E-242 |
| 2 | Reactive Müller glia (2) | fabp7a | 35.67048 | 2.07262969 | 9.1E-206 | 1.5E-202 |
| 3 | Reactive Müller glia (2) | gpm6aa | 31.36484 | 2.181623697 | 2.6E-167 | 2.5E-164 |
| 4 | Reactive Müller glia (2) | fxyd6l | 29.00227 | 2.877330303 | 1.2E-141 | 7.2E-139 |
| 5 | Reactive Müller glia (2) | mdka | 24.50395 | 1.458361506 | 8.2E-113 | 3.3E-110 |
| 6 | Reactive Müller glia (2) | col18a1a | 22.96548 | 2.439180136 | 2.68E-95 | 7.57E-93 |
| 7 | Reactive Müller glia (2) | crabp1a | 22.51464 | 1.35724318 | 1.82E-97 | 5.42E-95 |
| 8 | Reactive Müller glia (2) | CU467822.1 | 21.8247 | 1.936510324 | 6.76E-91 | 1.77E-88 |
| 9 | Reactive Müller glia (2) | f3a | 21.46131 | 1.343417048 | 1.94E-89 | 5.02E-87 |
| 10 | Reactive Müller glia (2) | pros1 | 21.12595 | 2.928258896 | 3.8E-81 | 8.15E-79 |
| 11 | Reactive Müller glia (2) | klf7b | 20.6625 | 2.281086683 | 1.22E-80 | 2.53E-78 |
| 12 | Reactive Müller glia (2) | zgc:165604 | 20.61534 | 1.651979804 | 9.21E-83 | 2.06E-80 |
| 13 | Reactive Müller glia (2) | marcksl1a | 20.12578 | 1.25527668 | 2.13E-80 | 4.35E-78 |
| 14 | Reactive Müller glia (2) | si:dkey-238o13.4 | 19.63945 | 2.279247999 | 1.82E-74 | 3.32E-72 |
| 15 | Reactive Müller glia (2) | mdkb | 18.74052 | 1.774334431 | 1.94E-69 | 2.97E-67 |
| 16 | Reactive Müller glia (2) | gfap | 17.7655 | 1.350174308 | 6.77E-64 | 8.89E-62 |
| 17 | Reactive Müller glia (2) | slc1a2b | 17.73886 | 2.154527426 | 5.26E-63 | 6.75E-61 |
| 18 | Reactive Müller glia (2) | zgc:165461 | 16.79933 | 7.665141106 | 1.78E-53 | 1.75E-51 |
| 19 | Reactive Müller glia (2) | si:ch211-251b21.1 | 16.71493 | 1.814291835 | 5.26E-56 | 5.46E-54 |
| 20 | Reactive Müller glia (2) | rlbp1a | 16.04219 | 2.598093271 | 1.15E-51 | 1.06E-49 |
| 21 | Reactive Müller glia (2) | myl9b | 15.97128 | 1.315761447 | 5.22E-53 | 5.02E-51 |
| 22 | Reactive Müller glia (2) | clcf1 | 15.64738 | 1.35037744 | 6.09E-51 | 5.46E-49 |
| 23 | Reactive Müller glia (2) | cotl1 | 15.10173 | 1.2638762 | 5.06E-48 | 4.09E-46 |
| 24 | Reactive Müller glia (2) | eml1 | 15.04819 | 2.15019083 | 1.52E-46 | 1.17E-44 |
| 25 | Reactive Müller glia (2) | crlf1a | 14.53827 | 1.168107033 | 7.25E-45 | 5.26E-43 |
| 26 | Reactive Müller glia (2) | gstp1 | 14.52729 | 1.762205362 | 2.39E-44 | 1.7E-42 |
| 27 | Reactive Müller glia (2) | fads2 | 14.16714 | 2.553449392 | 1.62E-41 | 1.04E-39 |
| 28 | Reactive Müller glia (2) | smad3a | 14.14918 | 1.93445015 | 9.53E-42 | 6.14E-40 |
| 29 | Reactive Müller glia (2) | si:ch73-352p4.8 | 14.01927 | 3.987175465 | 5.25E-40 | 3.11E-38 |
| 30 | Reactive Müller glia (2) | celf2 | 13.93621 | 2.079809904 | 1.3E-40 | 8.01E-39 |
| 31 | Reactive Müller glia (2) | plekhg2 | 13.88625 | 3.167042732 | 1.14E-39 | 6.66E-38 |
| 32 | Reactive Müller glia (2) | zgc:109949 | 13.74708 | 1.53017807 | 3.77E-40 | 2.25E-38 |
| 33 | Reactive Müller glia (2) | tjp2b | 13.68649 | 1.429957986 | 8.18E-40 | 4.82E-38 |
| 34 | Reactive Müller glia (2) | igfbp5b | 13.58487 | 1.896944046 | 3.83E-39 | 2.19E-37 |
| 35 | Reactive Müller glia (2) | sall1b | 13.51111 | 1.979957819 | 1.55E-38 | 8.61E-37 |
| 36 | Reactive Müller glia (2) | actb1 | 13.47996 | 0.658644021 | 2.9E-39 | 1.67E-37 |
| 37 | Reactive Müller glia (2) | itm2ba | 13.42153 | 0.942289829 | 6.73E-39 | 3.79E-37 |
| 38 | Reactive Müller glia (2) | ncam1a | 13.34969 | 1.571761489 | 5.38E-38 | 2.93E-36 |
| 39 | Reactive Müller glia (2) | ccni | 13.30744 | 1.353316069 | 6.68E-38 | 3.61E-36 |
| 40 | Reactive Müller glia (2) | eml2 | 13.29075 | 1.394143581 | 7.7E-38 | 4.15E-36 |
| 41 | Reactive Müller glia (2) | mgst3b | 13.17489 | 1.27916646 | 2.38E-37 | 1.25E-35 |
| 42 | Reactive Müller glia (2) | ywhag2 | 13.15285 | 2.172872066 | 1.56E-36 | 7.94E-35 |
| 43 | Reactive Müller glia (2) | rhbg | 12.90405 | 1.388804674 | 4.69E-36 | 2.36E-34 |
| 44 | Reactive Müller glia (2) | zgc:153867 | 12.84802 | 0.862836361 | 5.7E-36 | 2.85E-34 |
| 45 | Reactive Müller glia (2) | ISCU (1 of many) | 12.84055 | 1.269833088 | 1.35E-35 | 6.65E-34 |
| 46 | Reactive Müller glia (2) | il11b | 12.73038 | 2.29036808 | 1.83E-34 | 8.57E-33 |
| 47 | Reactive Müller glia (2) | igsf9ba | 12.72013 | 2.775602579 | 2.88E-34 | 1.34E-32 |
| 48 | Reactive Müller glia (2) | hmgn6 | 12.68202 | 0.950532019 | 4.49E-35 | 2.16E-33 |
| 49 | Reactive Müller glia (2) | zgc:86709 | 12.64639 | 2.087529898 | 3.24E-34 | 1.5E-32 |
| 50 | Reactive Müller glia (2) | arhgef4 | 12.62399 | 2.433920383 | 7.06E-34 | 3.21E-32 |
| 51 | Reactive Müller glia (2) | fabp3 | 12.48091 | 0.832520843 | 5.42E-34 | 2.48E-32 |
| 52 | Reactive Müller glia (2) | gpm6ab | 12.39938 | 1.428881168 | 1.58E-33 | 7.03E-32 |
| 53 | Reactive Müller glia (2) | cnn3a | 12.33085 | 1.018712282 | 3.16E-33 | 1.39E-31 |
| 54 | Reactive Müller glia (2) | si:dkey-28b4.7 | 12.19304 | 1.937535167 | 4.02E-32 | 1.69E-30 |
| 55 | Reactive Müller glia (2) | cdo1 | 12.15533 | 1.729387641 | 3.26E-32 | 1.38E-30 |
| 56 | Reactive Müller glia (2) | hivep2a | 12.14949 | 2.019537449 | 7.98E-32 | 3.31E-30 |
| 57 | Reactive Müller glia (2) | cbsb | 11.97805 | 1.878661871 | 4.08E-31 | 1.64E-29 |
| 58 | Reactive Müller glia (2) | kitlgb | 11.93795 | 2.118535757 | 9.32E-31 | 3.66E-29 |
| 59 | Reactive Müller glia (2) | akap12b | 11.93602 | 1.003729224 | 2.36E-31 | 9.6E-30 |
| 60 | Reactive Müller glia (2) | tp53inp1 | 11.93179 | 1.784459949 | 6.12E-31 | 2.43E-29 |
| 61 | Reactive Müller glia (2) | SMIM18 | 11.87499 | 3.085141182 | 3.34E-30 | 1.27E-28 |
| 62 | Reactive Müller glia (2) | flna | 11.83472 | 0.958186984 | 5.79E-31 | 2.3E-29 |
| 63 | Reactive Müller glia (2) | slc1a3b | 11.8222 | 5.071202278 | 1.19E-29 | 4.39E-28 |
| 64 | Reactive Müller glia (2) | efhd1 | 11.74102 | 1.481784344 | 2.67E-30 | 1.02E-28 |
| 65 | Reactive Müller glia (2) | elmsan1b | 11.69203 | 1.127770901 | 3.91E-30 | 1.48E-28 |
| 66 | Reactive Müller glia (2) | cygb2 | 11.67312 | 6.660600662 | 6.48E-29 | 2.29E-27 |
| 67 | Reactive Müller glia (2) | ttyh3b | 11.57314 | 2.33053875 | 3.88E-29 | 1.39E-27 |
| 68 | Reactive Müller glia (2) | s1pr1 | 11.56586 | 1.060813904 | 1.36E-29 | 5.04E-28 |
| 69 | Reactive Müller glia (2) | h3f3c | 11.54668 | 1.392056823 | 2.24E-29 | 8.17E-28 |
| 70 | Reactive Müller glia (2) | appa | 11.48259 | 0.975471556 | 3.25E-29 | 1.17E-27 |
| 71 | Reactive Müller glia (2) | FP102018.1 | 11.47569 | 2.04701972 | 8.78E-29 | 3.08E-27 |
| 72 | Reactive Müller glia (2) | dusp5 | 11.42468 | 1.124017239 | 5.28E-29 | 1.88E-27 |
| 73 | Reactive Müller glia (2) | col2a1a | 11.24889 | 1.526601434 | 5.94E-28 | 2E-26 |
| 74 | Reactive Müller glia (2) | atf5b | 11.19322 | 1.664454937 | 1.24E-27 | 4.11E-26 |
| 75 | Reactive Müller glia (2) | slc7a3a | 11.05061 | 1.27078557 | 3.44E-27 | 1.12E-25 |
| 76 | Reactive Müller glia (2) | mvp | 10.92912 | 0.836193025 | 9.07E-27 | 2.9E-25 |
| 77 | Reactive Müller glia (2) | zgc:162780 | 10.90802 | 1.421466231 | 1.78E-26 | 5.64E-25 |
| 78 | Reactive Müller glia (2) | zgc:100829 | 10.8524 | 1.64732337 | 3.62E-26 | 1.13E-24 |
| 79 | Reactive Müller glia (2) | ndrg4 | 10.70748 | 2.193386316 | 1.97E-25 | 5.96E-24 |
| 80 | Reactive Müller glia (2) | ndrg2 | 10.61652 | 2.331968546 | 5.66E-25 | 1.68E-23 |
| 81 | Reactive Müller glia (2) | acaca | 10.59206 | 1.666230202 | 5.17E-25 | 1.54E-23 |
| 82 | Reactive Müller glia (2) | swap70b | 10.28239 | 1.954082012 | 1.17E-23 | 3.21E-22 |
| 83 | Reactive Müller glia (2) | enpp6 | 10.26681 | 1.98515594 | 1.42E-23 | 3.88E-22 |
| 84 | Reactive Müller glia (2) | ephb2b | 10.16133 | 1.490687251 | 2.78E-23 | 7.49E-22 |
| 85 | Reactive Müller glia (2) | sparc | 10.15438 | 0.888876855 | 1.84E-23 | 4.99E-22 |
| 86 | Reactive Müller glia (2) | myl9a | 10.09955 | 1.868021727 | 6.68E-23 | 1.77E-21 |
| 87 | Reactive Müller glia (2) | chn1 | 10.07059 | 1.783767462 | 8.85E-23 | 2.33E-21 |
| 88 | Reactive Müller glia (2) | CR855337.1 | 10.0666 | 2.21694684 | 1.01E-22 | 2.67E-21 |
| 89 | Reactive Müller glia (2) | myo1ea | 10.06554 | 1.225029945 | 5.59E-23 | 1.48E-21 |
| 90 | Reactive Müller glia (2) | opcml | 9.885158 | 1.766567349 | 4E-22 | 1.02E-20 |
| 91 | Reactive Müller glia (2) | lepb | 9.792505 | 1.081987619 | 6.51E-22 | 1.64E-20 |
| 92 | Reactive Müller glia (2) | anxa4 | 9.78717 | 1.52517879 | 9.41E-22 | 2.35E-20 |
| 93 | Reactive Müller glia (2) | zic2b | 9.774028 | 0.915013194 | 6.3E-22 | 1.59E-20 |
| 94 | Reactive Müller glia (2) | dbn1 | 9.659241 | 0.945812345 | 2.22E-21 | 5.39E-20 |
| 95 | Reactive Müller glia (2) | nupr1b | 9.644771 | 2.353170395 | 4.3E-21 | 1.02E-19 |
| 96 | Reactive Müller glia (2) | lrrc4.1 | 9.626767 | 3.760903358 | 8.46E-21 | 1.97E-19 |
| 97 | Reactive Müller glia (2) | zgc:110182 | 9.54512 | 2.229369164 | 1.16E-20 | 2.68E-19 |
| 98 | Reactive Müller glia (2) | ca14 | 9.416307 | 1.495685816 | 2.42E-20 | 5.53E-19 |
| 99 | Reactive Müller glia (2) | appb | 9.374798 | 1.406201243 | 3.37E-20 | 7.65E-19 |

| | group | names | scores | logfoldchanges | pvals | pvals_adj |
|---|---|---|---|---|---|---|
| 100 | Reactive Müller glia (1) | hsp90aa1.2 | 82.35464 | 7.280426502 | 0 | 0 |
| 101 | Reactive Müller glia (1) | hsp70.3 | 66.28247 | 7.832823277 | 0 | 0 |
| 102 | Reactive Müller glia (1) | her15.1-1 | 60.54194 | 5.888670444 | 0 | 0 |
| 103 | Reactive Müller glia (1) | fosab | 55.58332 | 4.124118328 | 0 | 0 |
| 104 | Reactive Müller glia (1) | hsp70l | 49.03855 | 8.664011002 | 5.6E-263 | 2.2E-259 |
| 105 | Reactive Müller glia (1) | stm | 49.01005 | 10.58326435 | 1E-258 | 3.6E-255 |
| 106 | Reactive Müller glia (1) | id1 | 42.62602 | 4.362968922 | 6.9E-264 | 3.2E-260 |
| 107 | Reactive Müller glia (1) | hsp70.2 | 42.19383 | 7.396927834 | 2.7E-221 | 5.8E-218 |
| 108 | Reactive Müller glia (1) | her6 | 41.85655 | 4.55499506 | 5.6E-250 | 1.7E-246 |
| 109 | Reactive Müller glia (1) | si:ch211-222l21.1 | 41.21204 | 2.949662924 | 2.8E-246 | 7.7E-243 |
| 110 | Reactive Müller glia (1) | her15.1 | 39.92948 | 5.519450665 | 9.9E-216 | 2E-212 |
| 111 | Reactive Müller glia (1) | ubb | 38.23594 | 2.612923861 | 2E-227 | 4.7E-224 |
| 112 | Reactive Müller glia (1) | dnajb1b | 37.08304 | 4.597066879 | 4E-197 | 6.1E-194 |
| 113 | Reactive Müller glia (1) | jdp2b | 36.60299 | 3.446919441 | 6.4E-212 | 1.2E-208 |
| 114 | Reactive Müller glia (1) | cxcl18b | 35.82386 | 3.723559141 | 4.9E-207 | 8.5E-204 |
| 115 | Reactive Müller glia (1) | hsp70.1 | 34.68315 | 7.875595093 | 4.2E-170 | 4.1E-167 |
| 116 | Reactive Müller glia (1) | jun | 34.63765 | 2.626043081 | 1.1E-190 | 1.6E-187 |
| 117 | Reactive Müller glia (1) | tubb2b | 34.19307 | 3.273971796 | 5.8E-188 | 7.7E-185 |
| 118 | Reactive Müller glia (1) | si:ch73-281n10.2 | 33.63424 | 2.815275908 | 3.6E-188 | 5E-185 |
| 119 | Reactive Müller glia (1) | pcna | 33.39236 | 3.55332756 | 2.5E-184 | 3.2E-181 |
| 120 | Reactive Müller glia (1) | her12 | 33.0239 | 4.520014763 | 5.7E-173 | 6.4E-170 |
| 121 | Reactive Müller glia (1) | tuba8l4 | 32.91037 | 2.326611519 | 5.5E-182 | 6.7E-179 |
| 122 | Reactive Müller glia (1) | UBB | 32.78486 | 2.321974039 | 5.9E-181 | 6.8E-178 |
| 123 | Reactive Müller glia (1) | her4.2-1 | 32.63203 | 5.523843765 | 3.8E-161 | 2.7E-158 |
| 124 | Reactive Müller glia (1) | her4.2 | 32.58655 | 4.103734016 | 1.5E-171 | 1.5E-168 |
| 125 | Reactive Müller glia (1) | tubb4b | 32.01009 | 1.639407158 | 6.7E-172 | 7.2E-169 |
| 126 | Reactive Müller glia (1) | cirbpa | 31.76522 | 1.853838801 | 4.8E-167 | 4.4E-164 |
| 127 | Reactive Müller glia (1) | her4.1 | 31.74485 | 4.691707611 | 5.5E-163 | 4.2E-160 |
| 128 | Reactive Müller glia (1) | foxj1a | 31.5159 | 3.284177542 | 2.3E-165 | 2E-162 |
| 129 | Reactive Müller glia (1) | cebpb | 31.4061 | 3.698709488 | 2.5E-161 | 1.9E-158 |
| 130 | Reactive Müller glia (1) | junbb | 31.1249 | 2.515970707 | 2E-166 | 1.8E-163 |
| 131 | Reactive Müller glia (1) | si:ch73-335l21.4 | 31.02646 | 2.666006804 | 3.9E-158 | 2.7E-155 |
| 132 | Reactive Müller glia (1) | junba | 30.98528 | 2.227041006 | 1.4E-163 | 1.1E-160 |
| 133 | Reactive Müller glia (1) | socs3a | 30.86547 | 3.608341694 | 3.3E-161 | 2.4E-158 |
| 134 | Reactive Müller glia (1) | hmgb2a | 30.8064 | 2.7140553 | 3.6E-164 | 3E-161 |
| 135 | Reactive Müller glia (1) | hnrnpa1b | 30.73948 | 2.371783972 | 1.5E-163 | 1.2E-160 |
| 136 | Reactive Müller glia (1) | hmgb2b | 30.21128 | 2.393172264 | 6.7E-154 | 4.5E-151 |
| 137 | Reactive Müller glia (1) | arrdc3b | 29.89188 | 3.526479006 | 8.9E-151 | 5.8E-148 |
| 138 | Reactive Müller glia (1) | cdkn1a | 29.63683 | 4.312912941 | 4E-145 | 2.4E-142 |
| 139 | Reactive Müller glia (1) | tomm20a | 29.20352 | 2.382113695 | 1.6E-150 | 1E-147 |
| 140 | Reactive Müller glia (1) | hmga1a | 28.95779 | 2.122936964 | 2.9E-147 | 1.9E-144 |
| 141 | Reactive Müller glia (1) | tma7 | 28.69193 | 2.332645416 | 1.6E-146 | 9.7E-144 |
| 142 | Reactive Müller glia (1) | hspa5 | 28.6831 | 2.147303104 | 3.9E-144 | 2.3E-141 |
| 143 | Reactive Müller glia (1) | ascl1a | 28.26748 | 5.355374336 | 9.5E-131 | 5E-128 |
| 144 | Reactive Müller glia (1) | h2afvb | 27.82815 | 2.05033493 | 1.5E-133 | 8E-131 |
| 145 | Reactive Müller glia (1) | jund | 27.55827 | 2.139711618 | 4.2E-137 | 2.4E-134 |
| 146 | Reactive Müller glia (1) | si:ch211-156b7.4 | 27.55638 | 2.757825375 | 6.6E-137 | 3.7E-134 |
| 147 | Reactive Müller glia (1) | gadd45bb | 27.0352 | 3.035989761 | 4.5E-130 | 2.3E-127 |
| 148 | Reactive Müller glia (1) | tuba8l | 26.9602 | 2.795085669 | 2.1E-129 | 1E-126 |
| 149 | Reactive Müller glia (1) | nrarpa | 26.88822 | 2.839094162 | 1.8E-130 | 9E-128 |
| 150 | Reactive Müller glia (1) | nap1l1 | 26.84321 | 2.251416922 | 2E-131 | 1E-128 |
| 151 | Reactive Müller glia (1) | klf11b | 26.57303 | 3.741070986 | 6.2E-123 | 2.9E-120 |
| 152 | Reactive Müller glia (1) | rhov | 26.558 | 4.548403263 | 9.8E-120 | 4.5E-117 |
| 153 | Reactive Müller glia (1) | hspb1 | 26.17954 | 2.979319334 | 1.3E-123 | 5.9E-121 |
| 154 | Reactive Müller glia (1) | sox4a-1 | 26.11847 | 2.454756975 | 1.5E-125 | 7.1E-123 |
| 155 | Reactive Müller glia (1) | tuba1a | 25.96273 | 2.253772259 | 2.2E-124 | 1E-121 |
| 156 | Reactive Müller glia (1) | pim2 | 25.94703 | 4.804347515 | 1.8E-114 | 7.3E-112 |
| 157 | Reactive Müller glia (1) | khdrbs1a | 25.53657 | 1.588988662 | 3.1E-118 | 1.3E-115 |
| 158 | Reactive Müller glia (1) | anp32b | 25.48372 | 2.679930925 | 4.2E-119 | 1.9E-116 |
| 159 | Reactive Müller glia (1) | rrm2-1 | 25.3829 | 4.111489773 | 2.7E-113 | 1.1E-110 |
| 160 | Reactive Müller glia (1) | hsp90b1 | 25.09717 | 2.30643487 | 2.4E-117 | 1E-114 |
| 161 | Reactive Müller glia (1) | snrpe | 25.04205 | 2.148802519 | 3.9E-117 | 1.7E-114 |
| 162 | Reactive Müller glia (1) | atf3 | 24.81995 | 2.228286505 | 2.5E-115 | 1E-112 |
| 163 | Reactive Müller glia (1) | cirbpb | 24.78739 | 1.340621114 | 3.6E-111 | 1.4E-108 |
| 164 | Reactive Müller glia (1) | rpa2 | 24.6018 | 3.002773762 | 7.7E-112 | 3E-109 |
| 165 | Reactive Müller glia (1) | ran | 24.45253 | 1.411974311 | 4.9E-111 | 1.9E-108 |
| 166 | Reactive Müller glia (1) | nutf2l | 24.3949 | 3.412918568 | 1.1E-108 | 4.1E-106 |
| 167 | Reactive Müller glia (1) | dut | 23.97212 | 3.487446547 | 8.8E-105 | 3.2E-102 |
| 168 | Reactive Müller glia (1) | pclaf | 23.63413 | 4.183244228 | 3.1E-100 | 9.99E-98 |
| 169 | Reactive Müller glia (1) | mt-atp6 | 23.63085 | 1.458390713 | 5.7E-105 | 2.1E-102 |
| 170 | Reactive Müller glia (1) | TXN | 23.55486 | 1.474241495 | 1.2E-104 | 4.3E-102 |
| 171 | Reactive Müller glia (1) | ivns1abpb | 23.53427 | 4.192891121 | 5.2E-100 | 1.64E-97 |
| 172 | Reactive Müller glia (1) | ier2b | 23.34049 | 2.06543541 | 5.4E-104 | 2E-101 |
| 173 | Reactive Müller glia (1) | phlda2 | 23.32513 | 2.696411371 | 1.1E-103 | 3.9E-101 |
| 174 | Reactive Müller glia (1) | mcl1a | 23.31013 | 1.716981888 | 9.6E-104 | 3.4E-101 |
| 175 | Reactive Müller glia (1) | odc1 | 23.30475 | 1.990248203 | 1E-103 | 3.6E-101 |
| 176 | Reactive Müller glia (1) | ube2e2 | 23.27251 | 2.631728411 | 1.4E-101 | 4.7E-99 |
| 177 | Reactive Müller glia (1) | hspa4a | 23.24728 | 2.490544319 | 6.9E-102 | 2.3E-99 |
| 178 | Reactive Müller glia (1) | mycb | 23.18145 | 2.375152826 | 2.3E-102 | 7.7E-100 |
| 179 | Reactive Müller glia (1) | hspa8 | 22.97676 | 1.025163531 | 1.4E-99 | 4.21E-97 |
| 180 | Reactive Müller glia (1) | pfn2 | 22.96859 | 1.79169929 | 4.8E-101 | 1.6E-98 |
| 181 | Reactive Müller glia (1) | rrm1 | 22.96025 | 3.213908911 | 1.89E-98 | 5.69E-96 |
| 182 | Reactive Müller glia (1) | hmgn2 | 22.89884 | 1.936519623 | 1E-99 | 3.19E-97 |
| 183 | Reactive Müller glia (1) | ier5 | 22.86311 | 1.700139523 | 2.1E-100 | 6.99E-98 |
| 184 | Reactive Müller glia (1) | rpa3 | 22.81489 | 2.850908041 | 9.3E-99 | 2.83E-96 |
| 185 | Reactive Müller glia (1) | ncl | 22.77591 | 1.516149879 | 2.3E-99 | 7.06E-97 |
| 186 | Reactive Müller glia (1) | banf1 | 22.64566 | 2.745243549 | 3.24E-97 | 9.56E-95 |
| 187 | Reactive Müller glia (1) | fen1 | 22.64449 | 3.099192142 | 1.39E-96 | 4.06E-94 |
| 188 | Reactive Müller glia (1) | rbbp4 | 22.39164 | 2.448310137 | 3.76E-96 | 1.09E-93 |
| 189 | Reactive Müller glia (1) | cdca7a | 22.35177 | 4.124370575 | 7.94E-92 | 2.11E-89 |
| 190 | Reactive Müller glia (1) | socs3b | 22.27313 | 2.275952578 | 6.6E-96 | 1.89E-93 |
| 191 | Reactive Müller glia (1) | zfand2a | 22.06434 | 2.736085653 | 2.26E-92 | 6.13E-90 |
| 192 | Reactive Müller glia (1) | seta | 22.02438 | 1.771619797 | 5.23E-94 | 1.46E-91 |
| 193 | Reactive Müller glia (1) | chaf1a | 22.01164 | 2.596083641 | 2.86E-93 | 7.84E-91 |
| 194 | Reactive Müller glia (1) | hnrnpa0b | 21.99581 | 1.414581776 | 1.25E-93 | 3.45E-91 |
| 195 | Reactive Müller glia (1) | kpnb3 | 21.78032 | 2.509880304 | 2.19E-91 | 5.79E-89 |
| 196 | Reactive Müller glia (1) | her9 | 21.71943 | 2.040117264 | 7.8E-92 | 2.1E-89 |
| 197 | Reactive Müller glia (1) | CABZ01080568.1 | 21.61807 | 5.59968853 | 7.58E-85 | 1.8E-82 |
| 198 | Reactive Müller glia (1) | hells | 21.575 | 3.470985413 | 4.78E-88 | 1.2E-85 |
| 199 | Reactive Müller glia (1) | mcm5 | 21.51358 | 3.517920494 | 2.25E-87 | 5.55E-85 |
