## Supplementary_file_4 for "Single cell RNA sequencing unravels the transcriptional network underlying zebrafish retina regeneration"

**Differentially expressed genes between Reactive Müller glia (1) and Müller glia_Progenitors 1 clusters**

| | group | names | scores | logfoldchanges | pvals | pvals_adj |
|---|---|---|---|---|---|---|
| 0 | Reactive Müller glia (1) | hsp90aa1.2 | 78.96281 | 6.156524181 | 0 | 0 |
| 1 | Reactive Müller glia (1) | hsp70.3 | 67.37897 | 7.598965645 | 0 | 0 |
| 2 | Reactive Müller glia (1) | fosab | 60.32014 | 3.864514589 | 0 | 0 |
| 3 | Reactive Müller glia (1) | cxcl18b | 54.41115 | 6.127927303 | 2.3E-308 | 7.1E-305 |
| 4 | Reactive Müller glia (1) | si:ch73-335l21.4 | 54.25473 | 3.799617767 | 0 | 0 |
| 5 | Reactive Müller glia (1) | apoeb | 50.15687 | 4.11577177 | 0 | 0 |
| 6 | Reactive Müller glia (1) | hsp70l | 49.6275 | 9.198058128 | 3.9E-262 | 1.1E-258 |
| 7 | Reactive Müller glia (1) | stm | 49.15161 | 11.48877525 | 6.8E-259 | 1.7E-255 |
| 8 | Reactive Müller glia (1) | ubb | 48.80151 | 2.940044165 | 0 | 0 |
| 9 | Reactive Müller glia (1) | hspa5 | 48.58926 | 3.129906178 | 0 | 0 |
| 10 | Reactive Müller glia (1) | fosl1a | 47.78948 | 3.239794254 | 0 | 0 |
| 11 | Reactive Müller glia (1) | sox4a-1 | 43.41177 | 4.170948982 | 9.7E-249 | 2.2E-245 |
| 12 | Reactive Müller glia (1) | hsp70.2 | 42.73847 | 7.744447708 | 2.4E-221 | 3.6E-218 |
| 13 | Reactive Müller glia (1) | jdp2b | 40.42208 | 3.47694993 | 1.3E-235 | 2.2E-232 |
| 14 | Reactive Müller glia (1) | jund | 39.82693 | 2.880287886 | 1E-235 | 1.9E-232 |
| 15 | Reactive Müller glia (1) | jun | 38.33742 | 2.379800797 | 1.1E-244 | 2.2E-241 |
| 16 | Reactive Müller glia (1) | her6 | 37.62339 | 3.614906311 | 6.7E-220 | 9.8E-217 |
| 17 | Reactive Müller glia (1) | mmp9 | 37.06501 | 3.245724678 | 9E-214 | 1.2E-210 |
| 18 | Reactive Müller glia (1) | hspb1 | 37.02354 | 5.577278137 | 8E-188 | 8.8E-185 |
| 19 | Reactive Müller glia (1) | dnajb1b | 36.90395 | 4.200685978 | 1.2E-191 | 1.5E-188 |
| 20 | Reactive Müller glia (1) | socs3a | 35.88629 | 4.152430058 | 2.5E-189 | 3E-186 |
| 21 | Reactive Müller glia (1) | hsp70.1 | 34.8815 | 8.202496529 | 1.6E-170 | 1.2E-167 |
| 22 | Reactive Müller glia (1) | mcl1a | 34.03123 | 2.240658045 | 8E-189 | 9.2E-186 |
| 23 | Reactive Müller glia (1) | cebpb | 33.66378 | 3.929008961 | 1.4E-172 | 1.2E-169 |
| 24 | Reactive Müller glia (1) | zgc:158343 | 33.49094 | 2.860630989 | 6.7E-183 | 6.8E-180 |
| 25 | Reactive Müller glia (1) | rtn4a | 33.158 | 1.535396218 | 5.7E-195 | 7.5E-192 |
| 26 | Reactive Müller glia (1) | UBB | 33.15166 | 2.019450665 | 1.1E-185 | 1.2E-182 |
| 27 | Reactive Müller glia (1) | krt8 | 33.09382 | 2.681904554 | 1.5E-179 | 1.4E-176 |
| 28 | Reactive Müller glia (1) | mych | 32.59881 | 2.416787386 | 1.6E-182 | 1.6E-179 |
| 29 | Reactive Müller glia (1) | phlda2 | 32.40475 | 3.69194746 | 4.9E-163 | 3.3E-160 |
| 30 | Reactive Müller glia (1) | ier2b | 32.13038 | 2.724096537 | 3.2E-171 | 2.5E-168 |
| 31 | Reactive Müller glia (1) | sgk1 | 31.57028 | 2.193031549 | 8.9E-173 | 7.7E-170 |
| 32 | Reactive Müller glia (1) | junba | 31.5586 | 2.038266897 | 4.4E-180 | 4.2E-177 |
| 33 | Reactive Müller glia (1) | atp1b1a | 31.22713 | 2.143111944 | 3.2E-169 | 2.4E-166 |
| 34 | Reactive Müller glia (1) | mycb | 30.67942 | 3.167697191 | 3.2E-154 | 2E-151 |
| 35 | Reactive Müller glia (1) | junbb | 30.32054 | 2.221120358 | 4.2E-167 | 2.9E-164 |
| 36 | Reactive Müller glia (1) | s100a10b | 29.84934 | 4.043631554 | 3.1E-142 | 1.6E-139 |
| 37 | Reactive Müller glia (1) | gadd45bb | 29.53768 | 3.263278246 | 2.5E-146 | 1.5E-143 |
| 38 | Reactive Müller glia (1) | foxj1a | 29.46652 | 2.682247877 | 3.4E-148 | 2E-145 |
| 39 | Reactive Müller glia (1) | her15.1-1 | 29.25406 | 2.39308548 | 2.1E-158 | 1.4E-155 |
| 40 | Reactive Müller glia (1) | hsp90b1 | 29.07632 | 2.432119846 | 3.1E-146 | 1.7E-143 |
| 41 | Reactive Müller glia (1) | spry4 | 28.70327 | 2.905027628 | 1.1E-140 | 5.7E-138 |
| 42 | Reactive Müller glia (1) | klf6a | 28.34971 | 2.063533545 | 3E-146 | 1.7E-143 |
| 43 | Reactive Müller glia (1) | angptl4 | 27.8325 | 1.892593265 | 1.5E-142 | 8E-140 |
| 44 | Reactive Müller glia (1) | atp1a1b | 27.82399 | 2.743518829 | 7.5E-133 | 3.6E-130 |
| 45 | Reactive Müller glia (1) | arrdc3b | 27.66318 | 2.762813091 | 5E-133 | 2.4E-130 |
| 46 | Reactive Müller glia (1) | cdkn1a | 27.33998 | 3.319270849 | 1.2E-128 | 5.4E-126 |
| 47 | Reactive Müller glia (1) | nocta | 26.73047 | 2.115593433 | 2.4E-129 | 1.1E-126 |
| 48 | Reactive Müller glia (1) | zfand2a | 26.63722 | 3.617676735 | 6.3E-120 | 2.4E-117 |
| 49 | Reactive Müller glia (1) | txn | 26.63174 | 2.057679653 | 1.1E-131 | 5E-129 |
| 50 | Reactive Müller glia (1) | socs3b | 26.47211 | 2.58660078 | 3E-127 | 1.3E-124 |
| 51 | Reactive Müller glia (1) | eno1a | 26.30534 | 1.911908746 | 3.2E-128 | 1.4E-125 |
| 52 | Reactive Müller glia (1) | hbegfa | 26.1929 | 2.144310951 | 6.4E-127 | 2.8E-124 |
| 53 | Reactive Müller glia (1) | cd99 | 26.18384 | 3.468750477 | 1.2E-117 | 4.6E-115 |
| 54 | Reactive Müller glia (1) | krt18a.1 | 26.10007 | 4.836473465 | 2E-114 | 7.5E-112 |
| 55 | Reactive Müller glia (1) | klf11b | 26.01771 | 3.258551121 | 1.6E-117 | 6.2E-115 |
| 56 | Reactive Müller glia (1) | fosb | 25.87888 | 2.240445614 | 4.9E-123 | 2E-120 |
| 57 | Reactive Müller glia (1) | irs2b | 25.44503 | 3.12472558 | 7.3E-114 | 2.7E-111 |
| 58 | Reactive Müller glia (1) | atf3 | 25.40961 | 1.901364088 | 2E-120 | 8.3E-118 |
| 59 | Reactive Müller glia (1) | rhov | 25.17624 | 3.698608398 | 1.3E-110 | 4.7E-108 |
| 60 | Reactive Müller glia (1) | brd2a | 25.13917 | 1.458679438 | 4.2E-120 | 1.7E-117 |
| 61 | Reactive Müller glia (1) | gapdhs | 24.98846 | 1.26728642 | 3.2E-120 | 1.3E-117 |
| 62 | Reactive Müller glia (1) | cd63 | 24.68965 | 1.63531208 | 1.7E-114 | 6.4E-112 |
| 63 | Reactive Müller glia (1) | hspb8 | 24.66699 | 3.826961517 | 4.6E-106 | 1.5E-103 |
| 64 | Reactive Müller glia (1) | STMP1 | 24.27877 | 1.955182076 | 4.8E-110 | 1.7E-107 |
| 65 | Reactive Müller glia (1) | mCherry | 24.09944 | 2.188792229 | 2E-107 | 6.7E-105 |
| 66 | Reactive Müller glia (1) | mcl1b | 23.96723 | 2.348562956 | 6.6E-106 | 2.2E-103 |
| 67 | Reactive Müller glia (1) | pim3 | 23.68014 | 1.853738189 | 4.6E-107 | 1.5E-104 |
| 68 | Reactive Müller glia (1) | CR383676.1 | 23.60384 | 0.715097487 | 1E-107 | 3.5E-105 |
| 69 | Reactive Müller glia (1) | ascl1a | 23.57243 | 3.16264081 | 2.3E-101 | 7.4E-99 |
| 70 | Reactive Müller glia (1) | pdgfrb | 22.91039 | 3.372953415 | 3.34E-95 | 9.54E-93 |
| 71 | Reactive Müller glia (1) | dusp1 | 22.77743 | 2.231638432 | 9.59E-98 | 2.8E-95 |
| 72 | Reactive Müller glia (1) | tob1b | 22.76532 | 2.983662605 | 2.55E-94 | 7.21E-92 |
| 73 | Reactive Müller glia (1) | marcksl1b | 22.72348 | 1.133144617 | 3.1E-101 | 9.8E-99 |
| 74 | Reactive Müller glia (1) | pgam1a | 22.70318 | 2.195799828 | 3.14E-97 | 9.07E-95 |
| 75 | Reactive Müller glia (1) | c7b | 22.6899 | 3.574649334 | 1.65E-93 | 4.63E-91 |
| 76 | Reactive Müller glia (1) | rasgef1ba | 22.67658 | 1.824628711 | 1.91E-98 | 5.74E-96 |
| 77 | Reactive Müller glia (1) | si:dkey-7j14.6 | 22.38213 | 1.169266582 | 3.5E-99 | 1.08E-96 |
| 78 | Reactive Müller glia (1) | tomm20a | 22.37193 | 1.532642841 | 6.38E-98 | 1.9E-95 |
| 79 | Reactive Müller glia (1) | CABZ01080568.1 | 22.34726 | 7.764894009 | 1.13E-88 | 2.81E-86 |
| 80 | Reactive Müller glia (1) | tcima | 22.30956 | 3.769705772 | 2.57E-90 | 6.76E-88 |
| 81 | Reactive Müller glia (1) | anxa11a | 22.27368 | 2.720237017 | 5.81E-92 | 1.59E-89 |
| 82 | Reactive Müller glia (1) | fkbp5 | 22.25666 | 3.941752434 | 7.01E-90 | 1.83E-87 |
| 83 | Reactive Müller glia (1) | adgrg1-1 | 22.09657 | 3.025897741 | 1.53E-90 | 4.09E-88 |
| 84 | Reactive Müller glia (1) | alkbh3-1 | 22.04471 | 3.283715487 | 1.58E-89 | 4.04E-87 |
| 85 | Reactive Müller glia (1) | glula | 21.9769 | 2.318344831 | 3.56E-91 | 9.68E-89 |
| 86 | Reactive Müller glia (1) | pim2 | 21.8734 | 2.943208694 | 5.32E-89 | 1.33E-86 |
| 87 | Reactive Müller glia (1) | prdx6 | 21.48656 | 1.887133479 | 1.88E-89 | 4.78E-87 |
| 88 | Reactive Müller glia (1) | cdh2 | 21.46231 | 1.062956452 | 2.52E-92 | 6.99E-90 |
| 89 | Reactive Müller glia (1) | rgs5b | 21.31656 | 5.72307539 | 1.2E-82 | 2.69E-80 |
| 90 | Reactive Müller glia (1) | oaz1a | 21.25925 | 1.020347834 | 1.48E-90 | 3.98E-88 |
| 91 | Reactive Müller glia (1) | dab2ipb | 21.25077 | 2.017103434 | 2.01E-87 | 4.93E-85 |
| 92 | Reactive Müller glia (1) | pgk1 | 21.23523 | 1.918073297 | 2.04E-87 | 4.95E-85 |
| 93 | Reactive Müller glia (1) | tuft1a | 21.14848 | 2.924800873 | 4.52E-84 | 1.05E-81 |
| 94 | Reactive Müller glia (1) | ppp1r15a | 20.97945 | 2.838791609 | 6.81E-83 | 1.56E-80 |
| 95 | Reactive Müller glia (1) | hspa4a | 20.92429 | 1.903325438 | 1.69E-84 | 3.96E-82 |
| 96 | Reactive Müller glia (1) | rtn3 | 20.85957 | 1.949776292 | 1.01E-84 | 2.39E-82 |
| 97 | Reactive Müller glia (1) | bsg | 20.82198 | 1.313603282 | 2.63E-86 | 6.27E-84 |
| 98 | Reactive Müller glia (1) | MYO1D | 20.70807 | 3.313413382 | 1.72E-80 | 3.74E-78 |
| 99 | Reactive Müller glia (1) | rpz5 | 20.3744 | 4.091419697 | 1.48E-77 | 3.02E-75 |

| | group | names | scores | logfoldchanges | pvals | pvals_adj |
|---|---|---|---|---|---|---|
| 100 | Progenitors 1 | hmgn6 | 39.57415 | 2.13189435 | 4E-247 | 8.5E-244 |
| 101 | Progenitors 1 | ckbb | 36.08776 | 2.636459112 | 4.3E-225 | 7.1E-222 |
| 102 | Progenitors 1 | h3f3b.1-1 | 30.95333 | 2.0084548 | 5E-175 | 4.5E-172 |
| 103 | Progenitors 1 | fabp7a | 30.82239 | 1.702252626 | 1.7E-168 | 1.2E-165 |
| 104 | Progenitors 1 | gpm6aa | 30.8102 | 2.006668091 | 2.2E-172 | 1.8E-169 |
| 105 | Progenitors 1 | h3f3c | 30.08421 | 2.430395603 | 3.2E-168 | 2.3E-165 |
| 106 | Progenitors 1 | hmgn2 | 29.66204 | 1.723821163 | 4.4E-155 | 2.8E-152 |
| 107 | Progenitors 1 | stmn1a | 28.7132 | 2.58965683 | 3.1E-155 | 2E-152 |
| 108 | Progenitors 1 | acbd7 | 28.15553 | 2.370023727 | 2.5E-150 | 1.5E-147 |
| 109 | Progenitors 1 | fabp3 | 27.85391 | 1.387690187 | 5.6E-143 | 3.1E-140 |
| 110 | Progenitors 1 | si:dkey-238o13.4 | 27.16729 | 2.512182474 | 2.1E-141 | 1.1E-138 |
| 111 | Progenitors 1 | si:ch73-1a9.3 | 27.11039 | 1.595444441 | 2.3E-138 | 1.1E-135 |
| 112 | Progenitors 1 | hmgb1b | 25.72023 | 1.373405933 | 1.2E-124 | 4.9E-122 |
| 113 | Progenitors 1 | GFP | 25.22922 | 3.153699398 | 6.5E-124 | 2.7E-121 |
| 114 | Progenitors 1 | cadm3 | 23.78748 | 1.586513638 | 1.7E-111 | 6.3E-109 |
| 115 | Progenitors 1 | h3f3b.1 | 23.68885 | 1.421848297 | 4.9E-109 | 1.7E-106 |
| 116 | Progenitors 1 | crabp1a | 22.68151 | 1.223906994 | 4.1E-101 | 1.29E-98 |
| 117 | Progenitors 1 | si:ch211-288g17.3 | 22.52918 | 1.24346602 | 4.2E-100 | 1.3E-97 |
| 118 | Progenitors 1 | mdkb | 22.33049 | 1.663776517 | 1.5E-99 | 4.75E-97 |
| 119 | Progenitors 1 | hmgb1a | 22.25559 | 1.198459744 | 8.82E-98 | 2.6E-95 |
| 120 | Progenitors 1 | hmgb2a | 21.26914 | 1.317965388 | 1.45E-89 | 3.75E-87 |
| 121 | Progenitors 1 | tdh | 20.94719 | 1.739386201 | 3.13E-89 | 7.87E-87 |
| 122 | Progenitors 1 | nr2e1 | 20.64315 | 2.29147315 | 6.27E-87 | 1.51E-84 |
| 123 | Progenitors 1 | ptmab | 20.28705 | 0.83880198 | 7.24E-83 | 1.64E-80 |
| 124 | Progenitors 1 | tspan7 | 20.16095 | 1.419094563 | 3.57E-83 | 8.25E-81 |
| 125 | Progenitors 1 | tmsb | 19.9558 | 1.716845274 | 8.5E-82 | 1.87E-79 |
| 126 | Progenitors 1 | cbsb | 19.62659 | 2.234595299 | 2.18E-79 | 4.64E-77 |
| 127 | Progenitors 1 | marcksb | 19.60996 | 1.083263874 | 1.88E-78 | 3.95E-76 |
| 128 | Progenitors 1 | hmgb2b | 19.49424 | 0.92587024 | 5.55E-77 | 1.12E-74 |
| 129 | Progenitors 1 | dek | 19.21368 | 1.444202542 | 3.06E-76 | 6.06E-74 |
| 130 | Progenitors 1 | CU467822.1 | 18.43581 | 1.455706596 | 1.32E-70 | 2.35E-68 |
| 131 | Progenitors 1 | tuba1c | 17.86642 | 1.410547018 | 6.21E-67 | 1.02E-64 |
| 132 | Progenitors 1 | sox9b | 17.68629 | 1.859355927 | 9.07E-66 | 1.48E-63 |
| 133 | Progenitors 1 | si:dkey-28b4.7 | 17.62914 | 2.130151272 | 2.48E-65 | 3.98E-63 |
| 134 | Progenitors 1 | adh5 | 17.57185 | 1.542553425 | 5.9E-65 | 9.38E-63 |
| 135 | Progenitors 1 | rcor2 | 17.48 | 3.061095953 | 7.46E-64 | 1.18E-61 |
| 136 | Progenitors 1 | ankrd12 | 17.23396 | 1.494663835 | 1.01E-62 | 1.56E-60 |
| 137 | Progenitors 1 | sesn2 | 17.20127 | 1.955556989 | 1.7E-62 | 2.59E-60 |
| 138 | Progenitors 1 | crabp2a | 17.02717 | 1.986029267 | 2.32E-61 | 3.47E-59 |
| 139 | Progenitors 1 | psat1 | 16.95135 | 1.092074513 | 2.4E-60 | 3.54E-58 |
| 140 | Progenitors 1 | smc1al | 16.65961 | 1.145083785 | 1.25E-58 | 1.76E-56 |
| 141 | Progenitors 1 | klf7b | 16.64874 | 1.638218045 | 7.06E-59 | 1.01E-56 |
| 142 | Progenitors 1 | hmgb3a | 16.52694 | 1.268060207 | 5.77E-58 | 7.95E-56 |
| 143 | Progenitors 1 | marcksa | 16.49779 | 1.599452138 | 6.94E-58 | 9.47E-56 |
| 144 | Progenitors 1 | apex1 | 16.48928 | 1.374166369 | 9.17E-58 | 1.24E-55 |
| 145 | Progenitors 1 | h2afvb | 16.45271 | 0.72024411 | 5.42E-57 | 7.26E-55 |
| 146 | Progenitors 1 | rcc2 | 16.30293 | 1.532214999 | 1.16E-56 | 1.55E-54 |
| 147 | Progenitors 1 | sod1 | 16.26657 | 1.188361645 | 2.97E-56 | 3.88E-54 |
| 148 | Progenitors 1 | slc7a3a | 16.21495 | 1.498661041 | 4.52E-56 | 5.88E-54 |
| 149 | Progenitors 1 | rpl12 | 16.15476 | 0.574337721 | 8.85E-55 | 1.12E-52 |
| 150 | Progenitors 1 | celf2 | 16.04774 | 1.949820876 | 5.25E-55 | 6.7E-53 |
| 151 | Progenitors 1 | top2b | 15.60833 | 1.320457339 | 3.1E-52 | 3.71E-50 |
| 152 | Progenitors 1 | tp53inp1 | 15.2565 | 1.763965011 | 3.68E-50 | 4.32E-48 |
| 153 | Progenitors 1 | rab39ba | 15.23841 | 3.177596807 | 1.37E-49 | 1.54E-47 |
| 154 | Progenitors 1 | h1f0 | 15.2238 | 1.353938341 | 9.21E-50 | 1.06E-47 |
| 155 | Progenitors 1 | ptprz1a | 15.19141 | 2.72170639 | 1.79E-49 | 2.02E-47 |
| 156 | Progenitors 1 | tp53inp2 | 15.0532 | 1.484325528 | 6.24E-49 | 6.91E-47 |
| 157 | Progenitors 1 | nova2 | 15.0495 | 1.107542992 | 9.37E-49 | 1.03E-46 |
| 158 | Progenitors 1 | seta | 14.97295 | 0.802234411 | 4.56E-48 | 4.95E-46 |
| 159 | Progenitors 1 | ubtf | 14.81291 | 1.303162575 | 1.9E-47 | 1.98E-45 |
| 160 | Progenitors 1 | prdx2 | 14.73677 | 0.80364126 | 9.61E-47 | 9.85E-45 |
| 161 | Progenitors 1 | fads2 | 14.67992 | 2.178925991 | 1.14E-46 | 1.16E-44 |
| 162 | Progenitors 1 | si:ch211-137a8.4 | 14.45589 | 1.096811056 | 3.15E-45 | 3.15E-43 |
| 163 | Progenitors 1 | lbr | 14.42498 | 1.25320518 | 3.42E-45 | 3.41E-43 |
| 164 | Progenitors 1 | pou3f1 | 14.19874 | 4.809452057 | 3.99E-43 | 3.81E-41 |
| 165 | Progenitors 1 | creb1a | 14.10757 | 1.335472941 | 2.18E-43 | 2.1E-41 |
| 166 | Progenitors 1 | atf5b | 14.02652 | 1.596842647 | 5.81E-43 | 5.51E-41 |
| 167 | Progenitors 1 | smarca4a | 13.69398 | 1.074635983 | 5.26E-41 | 4.75E-39 |
| 168 | Progenitors 1 | sept4a | 13.54579 | 1.694412708 | 2.72E-40 | 2.41E-38 |
| 169 | Progenitors 1 | nucks1a | 13.25495 | 0.945909262 | 1.43E-38 | 1.21E-36 |
| 170 | Progenitors 1 | phc2a | 13.16774 | 1.356100798 | 3.13E-38 | 2.62E-36 |
| 171 | Progenitors 1 | acin1a | 13.07517 | 0.839772761 | 1.7E-37 | 1.4E-35 |
| 172 | Progenitors 1 | nfyba | 12.89905 | 1.451509953 | 8.1E-37 | 6.54E-35 |
| 173 | Progenitors 1 | atrx | 12.86515 | 0.781366765 | 1.97E-36 | 1.56E-34 |
| 174 | Progenitors 1 | si:ch73-215f7.1 | 12.86413 | 2.224107027 | 1.46E-36 | 1.17E-34 |
| 175 | Progenitors 1 | tnrc6c2 | 12.84728 | 2.218010187 | 1.74E-36 | 1.39E-34 |
| 176 | Progenitors 1 | snrpd1 | 12.8338 | 0.702584565 | 3.64E-36 | 2.83E-34 |
| 177 | Progenitors 1 | si:ch73-46j18.5 | 12.8279 | 0.922019362 | 2.6E-36 | 2.04E-34 |
| 178 | Progenitors 1 | hmga1a | 12.82499 | 0.682489455 | 2.61E-36 | 2.05E-34 |
| 179 | Progenitors 1 | baz2ba | 12.82041 | 1.115893126 | 2.43E-36 | 1.92E-34 |
| 180 | Progenitors 1 | gpm6ab | 12.79813 | 1.261794567 | 3.36E-36 | 2.62E-34 |
| 181 | Progenitors 1 | slc6a15 | 12.78617 | 4.334433556 | 8.89E-36 | 6.86E-34 |
| 182 | Progenitors 1 | csdc2a | 12.52681 | 1.596227884 | 6.94E-35 | 5.15E-33 |
| 183 | Progenitors 1 | anp32a | 12.512 | 0.839897811 | 1.07E-34 | 7.87E-33 |
| 184 | Progenitors 1 | cebpg | 12.50603 | 1.050130963 | 9.83E-35 | 7.22E-33 |
| 185 | Progenitors 1 | si:dkey-108k21.10 | 12.50057 | 1.631147027 | 9.59E-35 | 7.06E-33 |
| 186 | Progenitors 1 | polb | 12.32132 | 1.647771955 | 7.72E-34 | 5.54E-32 |
| 187 | Progenitors 1 | hdac1 | 12.27381 | 0.751041532 | 1.95E-33 | 1.38E-31 |
| 188 | Progenitors 1 | ddx39ab | 12.24022 | 0.691922486 | 2.98E-33 | 2.09E-31 |
| 189 | Progenitors 1 | h3f3d | 12.20215 | 0.499156833 | 5.01E-33 | 3.48E-31 |
| 190 | Progenitors 1 | znf1035 | 12.1861 | 1.370621681 | 3.76E-33 | 2.63E-31 |
| 191 | Progenitors 1 | slc1a4 | 12.17181 | 1.805157423 | 4.41E-33 | 3.08E-31 |
| 192 | Progenitors 1 | actl6a | 12.15832 | 0.940167129 | 5.95E-33 | 4.11E-31 |
| 193 | Progenitors 1 | selenow2a | 12.13408 | 1.186491489 | 7.04E-33 | 4.82E-31 |
| 194 | Progenitors 1 | klf7a | 12.12547 | 2.643581629 | 9.99E-33 | 6.82E-31 |
| 195 | Progenitors 1 | zgc:153867 | 12.12113 | 0.730895638 | 1.14E-32 | 7.76E-31 |
| 196 | Progenitors 1 | pacsin1b | 12.09484 | 2.469278336 | 1.3E-32 | 8.79E-31 |
| 197 | Progenitors 1 | ap1s2 | 11.9897 | 1.187903285 | 3.57E-32 | 2.38E-30 |
| 198 | Progenitors 1 | nudt21 | 11.98962 | 1.14147234 | 3.74E-32 | 2.49E-30 |
| 199 | Progenitors 1 | ncam1a | 11.92678 | 1.204928756 | 7.43E-32 | 4.91E-30 |
