## Supplementary_file_5 for "Single cell RNA sequencing unravels the transcriptional network underlying zebrafish retina regeneration"

**Differentially expressed genes between Reactive Müller glia (2) and Müller glia_Progenitors 1 clusters**

| | group | names | scores | logfoldchanges | pvals | pvals_adj |
|---|---|---|---|---|---|---|
| 0 | Reactive Müller glia (2) | marcksl1a | 28.45223 | 1.566915274 | 3.2E-141 | 1.7E-138 |
| 1 | Reactive Müller glia (2) | slc1a2b | 27.86047 | 4.331185341 | 5.7E-119 | 2E-116 |
| 2 | Reactive Müller glia (2) | fxyd6l | 27.35475 | 2.334259748 | 3.3E-125 | 1.3E-122 |
| 3 | Reactive Müller glia (2) | zgc:165604 | 26.48078 | 2.034055948 | 9.8E-122 | 3.6E-119 |
| 4 | Reactive Müller glia (2) | crlf1a | 26.27136 | 2.145042896 | 3.5E-124 | 1.3E-121 |
| 5 | Reactive Müller glia (2) | dusp5 | 25.35421 | 2.675037622 | 1.6E-109 | 4.4E-107 |
| 6 | Reactive Müller glia (2) | sparc | 24.78819 | 2.37652421 | 3.9E-106 | 1.1E-103 |
| 7 | Reactive Müller glia (2) | f3a | 24.01298 | 1.541461587 | 1.9E-110 | 5.4E-108 |
| 8 | Reactive Müller glia (2) | hbegfa | 23.4558 | 2.121440887 | 5.5E-100 | 1.36E-97 |
| 9 | Reactive Müller glia (2) | cnn2 | 23.029 | 1.925940514 | 3.17E-97 | 7.38E-95 |
| 10 | Reactive Müller glia (2) | mdka | 22.80873 | 1.14225626 | 2.24E-98 | 5.43E-96 |
| 11 | Reactive Müller glia (2) | cnn3a | 22.71172 | 1.950399995 | 1.3E-93 | 2.81E-91 |
| 12 | Reactive Müller glia (2) | CR383676.1 | 22.37306 | 0.70974195 | 1.19E-94 | 2.66E-92 |
| 13 | Reactive Müller glia (2) | krt8 | 22.35656 | 2.220809937 | 3.63E-90 | 7.07E-88 |
| 14 | Reactive Müller glia (2) | mvp | 21.91733 | 1.678019047 | 3.08E-89 | 5.77E-87 |
| 15 | Reactive Müller glia (2) | selenop | 21.82425 | 3.010911703 | 7.67E-84 | 1.28E-81 |
| 16 | Reactive Müller glia (2) | txn | 21.76608 | 1.946496964 | 1.2E-88 | 2.24E-86 |
| 17 | Reactive Müller glia (2) | si:ch1073-303k11.2 | 21.71447 | 1.689989328 | 1.04E-89 | 1.99E-87 |
| 18 | Reactive Müller glia (2) | gfap | 21.41233 | 1.51608026 | 2.38E-86 | 4.12E-84 |
| 19 | Reactive Müller glia (2) | icn | 21.24722 | 3.998539448 | 6.96E-79 | 1.02E-76 |
| 20 | Reactive Müller glia (2) | atp1b1a | 21.22201 | 1.900651455 | 2.81E-83 | 4.61E-81 |
| 21 | Reactive Müller glia (2) | sgk1 | 21.03592 | 1.685407877 | 4.81E-84 | 8.07E-82 |
| 22 | Reactive Müller glia (2) | rhbg | 21.01989 | 2.394856691 | 6.02E-81 | 9.2E-79 |
| 23 | Reactive Müller glia (2) | col18a1a | 20.84691 | 1.998223186 | 8.44E-81 | 1.28E-78 |
| 24 | Reactive Müller glia (2) | myl6 | 20.33518 | 1.38188076 | 4.48E-81 | 6.89E-79 |
| 25 | Reactive Müller glia (2) | c7b | 20.18061 | 3.561120272 | 1.75E-73 | 2.25E-71 |
| 26 | Reactive Müller glia (2) | clcf1 | 20.05112 | 1.709903002 | 1.36E-77 | 1.92E-75 |
| 27 | Reactive Müller glia (2) | alkbh3-1 | 19.66386 | 3.396778822 | 1.85E-70 | 2.12E-68 |
| 28 | Reactive Müller glia (2) | rlbp1a | 19.22552 | 3.395123959 | 5.27E-68 | 5.7E-66 |
| 29 | Reactive Müller glia (2) | cd99 | 19.18551 | 2.994940758 | 3.83E-68 | 4.16E-66 |
| 30 | Reactive Müller glia (2) | grb10b | 18.90202 | 1.879710555 | 1.33E-68 | 1.48E-66 |
| 31 | Reactive Müller glia (2) | rtn4a | 18.87692 | 1.03217411 | 2.59E-71 | 3.05E-69 |
| 32 | Reactive Müller glia (2) | anxa13 | 18.84404 | 1.886629581 | 2.43E-68 | 2.67E-66 |
| 33 | Reactive Müller glia (2) | apoeb | 18.75928 | 2.621199131 | 3.58E-66 | 3.61E-64 |
| 34 | Reactive Müller glia (2) | aplp2 | 18.69975 | 2.014311552 | 8.7E-67 | 9.03E-65 |
| 35 | Reactive Müller glia (2) | gstp1 | 18.67665 | 2.208734035 | 2.92E-66 | 2.96E-64 |
| 36 | Reactive Müller glia (2) | mych | 18.4314 | 1.633396864 | 2.51E-67 | 2.64E-65 |
| 37 | Reactive Müller glia (2) | cdo1 | 18.30334 | 3.025168657 | 4.74E-63 | 4.42E-61 |
| 38 | Reactive Müller glia (2) | pros1 | 18.17524 | 2.132115126 | 1.74E-63 | 1.63E-61 |
| 39 | Reactive Müller glia (2) | myl9b | 18.01382 | 1.368223786 | 9.92E-65 | 9.64E-63 |
| 40 | Reactive Müller glia (2) | spock3 | 17.80174 | 3.213432789 | 5.26E-60 | 4.58E-58 |
| 41 | Reactive Müller glia (2) | acbd7 | 17.78956 | 1.1685009 | 1.76E-64 | 1.69E-62 |
| 42 | Reactive Müller glia (2) | ckbb | 17.64818 | 1.098384976 | 5.58E-64 | 5.28E-62 |
| 43 | Reactive Müller glia (2) | eml2 | 17.47475 | 1.844926476 | 6.09E-60 | 5.29E-58 |
| 44 | Reactive Müller glia (2) | efhd1 | 17.42809 | 2.39108777 | 5.79E-59 | 4.9E-57 |
| 45 | Reactive Müller glia (2) | flna | 17.09739 | 1.316082716 | 2.74E-59 | 2.35E-57 |
| 46 | Reactive Müller glia (2) | zfand5a | 16.94324 | 1.677537322 | 4.14E-57 | 3.38E-55 |
| 47 | Reactive Müller glia (2) | tob1b | 16.61486 | 2.226042986 | 2.02E-54 | 1.55E-52 |
| 48 | Reactive Müller glia (2) | vmp1 | 16.4526 | 1.245399594 | 1.85E-56 | 1.47E-54 |
| 49 | Reactive Müller glia (2) | bzw1b | 16.2472 | 1.006033659 | 3.21E-54 | 2.44E-52 |
| 50 | Reactive Müller glia (2) | lepb | 16.06698 | 1.834574223 | 2.6E-52 | 1.9E-50 |
| 51 | Reactive Müller glia (2) | irs2b | 16.03065 | 2.343751192 | 3.67E-51 | 2.6E-49 |
| 52 | Reactive Müller glia (2) | kitlgb | 15.9755 | 3.882472038 | 9.47E-50 | 6.4E-48 |
| 53 | Reactive Müller glia (2) | errfi1a | 15.63933 | 1.828522801 | 1.24E-49 | 8.27E-48 |
| 54 | Reactive Müller glia (2) | zic2b | 15.62195 | 1.328985333 | 3.32E-50 | 2.28E-48 |
| 55 | Reactive Müller glia (2) | fosl1a | 15.59316 | 1.633912325 | 7.88E-50 | 5.33E-48 |
| 56 | Reactive Müller glia (2) | epas1b | 15.58087 | 2.754811764 | 2.97E-48 | 1.88E-46 |
| 57 | Reactive Müller glia (2) | s100a10b | 15.4264 | 2.781081438 | 2.69E-47 | 1.63E-45 |
| 58 | Reactive Müller glia (2) | il11b | 15.40082 | 3.162389517 | 4.5E-47 | 2.7E-45 |
| 59 | Reactive Müller glia (2) | ca14 | 15.34694 | 3.267858028 | 1.04E-46 | 6.15E-45 |
| 60 | Reactive Müller glia (2) | vim | 15.3415 | 1.479634047 | 1.15E-48 | 7.4E-47 |
| 61 | Reactive Müller glia (2) | klf6a | 15.33034 | 1.333794951 | 1.02E-48 | 6.64E-47 |
| 62 | Reactive Müller glia (2) | cadm4 | 15.29171 | 1.475498557 | 3.98E-48 | 2.5E-46 |
| 63 | Reactive Müller glia (2) | ppdpfb | 15.23914 | 0.702867806 | 3.86E-49 | 2.52E-47 |
| 64 | Reactive Müller glia (2) | elovl1b | 15.10914 | 1.50200212 | 4.57E-47 | 2.73E-45 |
| 65 | Reactive Müller glia (2) | ywhag2 | 14.81845 | 2.544952631 | 2.39E-44 | 1.28E-42 |
| 66 | Reactive Müller glia (2) | col2a1a | 14.76239 | 2.119177818 | 1.91E-44 | 1.03E-42 |
| 67 | Reactive Müller glia (2) | igfbp5b | 14.7455 | 1.948260307 | 1.42E-44 | 7.73E-43 |
| 68 | Reactive Müller glia (2) | hsd11b2 | 14.58801 | 3.330842733 | 7.79E-43 | 3.96E-41 |
| 69 | Reactive Müller glia (2) | pygl | 14.56145 | 2.348546505 | 4.53E-43 | 2.3E-41 |
| 70 | Reactive Müller glia (2) | chmp4ba | 14.19309 | 1.743988395 | 1.03E-41 | 5E-40 |
| 71 | Reactive Müller glia (2) | sall1b | 14.10458 | 1.989616275 | 3.58E-41 | 1.7E-39 |
| 72 | Reactive Müller glia (2) | dhrs13l1 | 14.09953 | 3.333694935 | 2.45E-40 | 1.13E-38 |
| 73 | Reactive Müller glia (2) | wfdc2 | 14.05942 | 3.250406742 | 3.65E-40 | 1.67E-38 |
| 74 | Reactive Müller glia (2) | si:dkey-7j14.6 | 13.72148 | 0.804624915 | 2.25E-40 | 1.04E-38 |
| 75 | Reactive Müller glia (2) | dab2ipb | 13.60768 | 1.51547873 | 4.83E-39 | 2.13E-37 |
| 76 | Reactive Müller glia (2) | spry4 | 13.53919 | 1.860926628 | 2.3E-38 | 9.92E-37 |
| 77 | Reactive Müller glia (2) | tnfrsf21 | 13.41025 | 1.682223201 | 6.89E-38 | 2.9E-36 |
| 78 | Reactive Müller glia (2) | akap12b | 13.40001 | 1.028597713 | 3.26E-38 | 1.39E-36 |
| 79 | Reactive Müller glia (2) | cotl1 | 13.07852 | 0.965141952 | 7.34E-37 | 2.98E-35 |
| 80 | Reactive Müller glia (2) | zgc:162780 | 13.02275 | 1.687155247 | 6.8E-36 | 2.65E-34 |
| 81 | Reactive Müller glia (2) | midn | 13.00825 | 0.977434278 | 1.33E-36 | 5.37E-35 |
| 82 | Reactive Müller glia (2) | ctsla | 13.00383 | 0.776681185 | 2.63E-36 | 1.04E-34 |
| 83 | Reactive Müller glia (2) | ecm1b | 12.93636 | 3.175004721 | 7.54E-35 | 2.83E-33 |
| 84 | Reactive Müller glia (2) | zgc:165461 | 12.84795 | 2.513814688 | 1.32E-34 | 4.93E-33 |
| 85 | Reactive Müller glia (2) | ahnak | 12.78009 | 3.018579006 | 3.57E-34 | 1.31E-32 |
| 86 | Reactive Müller glia (2) | ptgdsb.2 | 12.69268 | 3.252966642 | 1.41E-33 | 5.07E-32 |
| 87 | Reactive Müller glia (2) | gsna | 12.68134 | 2.104523659 | 4.62E-34 | 1.69E-32 |
| 88 | Reactive Müller glia (2) | irf2 | 12.64866 | 1.739259839 | 4.03E-34 | 1.48E-32 |
| 89 | Reactive Müller glia (2) | lygl1 | 12.64654 | 2.5468297 | 1.2E-33 | 4.34E-32 |
| 90 | Reactive Müller glia (2) | igsf9ba | 12.58847 | 2.519938707 | 2.06E-33 | 7.4E-32 |
| 91 | Reactive Müller glia (2) | fam129bb | 12.57495 | 1.397484899 | 6.31E-34 | 2.29E-32 |
| 92 | Reactive Müller glia (2) | myl9a | 12.53734 | 2.643761158 | 3.7E-33 | 1.31E-31 |
| 93 | Reactive Müller glia (2) | cdh2 | 12.519 | 0.663452744 | 2.37E-34 | 8.79E-33 |
| 94 | Reactive Müller glia (2) | cd82a | 12.50103 | 0.811309636 | 4.55E-34 | 1.67E-32 |
| 95 | Reactive Müller glia (2) | inhbaa | 12.4466 | 2.094907999 | 5.46E-33 | 1.92E-31 |
| 96 | Reactive Müller glia (2) | zgc:174888 | 12.35362 | 3.04491663 | 3.45E-32 | 1.19E-30 |
| 97 | Reactive Müller glia (2) | nocta | 12.29703 | 1.160972834 | 8.28E-33 | 2.89E-31 |
| 98 | Reactive Müller glia (2) | nfkbiaa | 12.17441 | 2.496165752 | 1.43E-31 | 4.77E-30 |
| 99 | Reactive Müller glia (2) | slc12a4 | 12.134 | 1.549150229 | 9.04E-32 | 3.05E-30 |

| | group | names | scores | logfoldchanges | pvals | pvals_adj |
|---|---|---|---|---|---|---|
| **100** | Progenitors 1 | hmgb2a | 59.88311 | 4.032020569 | 0 | 0 |
| **101** | Progenitors 1 | hmgn2 | 53.28411 | 3.660340786 | 0 | 7.1E-308 |
| **102** | Progenitors 1 | hmgb2b | 48.89518 | 3.319042683 | 4.3E-280 | 2.4E-276 |
| **103** | Progenitors 1 | h3f3b.1 | 46.57986 | 2.922519445 | 4.4E-305 | 3.1E-301 |
| **104** | Progenitors 1 | tubb2b | 45.28305 | 3.77976346 | 0 | 0 |
| **105** | Progenitors 1 | si:ch211-222l21.1 | 44.287 | 3.044552565 | 9.9E-273 | 3.4E-269 |
| **106** | Progenitors 1 | hmga1a | 42.31578 | 2.805426359 | 8.4E-263 | 2.6E-259 |
| **107** | Progenitors 1 | si:ch73-281n10.2 | 42.1779 | 3.147583485 | 6.9E-276 | 3.2E-272 |
| **108** | Progenitors 1 | h2afvb | 42.16171 | 2.7705791 | 5.3E-232 | 1.1E-228 |
| **109** | Progenitors 1 | pcna | 41.41436 | 3.794421434 | 6.9E-273 | 2.7E-269 |
| **110** | Progenitors 1 | seta | 39.36641 | 2.573854208 | 1.9E-237 | 4.5E-234 |
| **111** | Progenitors 1 | si:ch211-288g17.3 | 39.03107 | 2.786085606 | 1.9E-225 | 3.6E-222 |
| **112** | Progenitors 1 | si:ch211-156b7.4 | 38.83523 | 3.283548355 | 2.8E-247 | 7.1E-244 |
| **113** | Progenitors 1 | dek | 38.75076 | 3.699343443 | 2.3E-248 | 6.4E-245 |
| **114** | Progenitors 1 | hmgb1b | 37.81065 | 2.306586742 | 1.5E-214 | 2.4E-211 |
| **115** | Progenitors 1 | hmgb1a | 37.6093 | 2.51217103 | 1E-212 | 1.5E-209 |
| **116** | Progenitors 1 | stmn1a | 36.73283 | 3.838213921 | 1E-226 | 2E-223 |
| **117** | Progenitors 1 | mki67 | 36.10389 | 4.584386349 | 5.6E-216 | 9.7E-213 |
| **118** | Progenitors 1 | cirbpa | 35.11591 | 1.893434405 | 3.1E-187 | 4E-184 |
| **119** | Progenitors 1 | rrm1 | 33.81368 | 3.679839134 | 9.2E-200 | 1.3E-196 |
| **120** | Progenitors 1 | rbbp4 | 33.75729 | 2.964738607 | 1.1E-199 | 1.5E-196 |
| **121** | Progenitors 1 | ptmab | 32.60604 | 1.622532725 | 1.8E-168 | 1.7E-165 |
| **122** | Progenitors 1 | lbr | 32.29927 | 3.835795403 | 7.5E-184 | 9.4E-181 |
| **123** | Progenitors 1 | rrm2-1 | 32.17185 | 4.305764198 | 4.3E-181 | 5E-178 |
| **124** | Progenitors 1 | cbx3a | 32.10963 | 2.287667274 | 5E-180 | 5.4E-177 |
| **125** | Progenitors 1 | snrpd1 | 31.98677 | 2.024102688 | 1.2E-171 | 1.2E-168 |
| **126** | Progenitors 1 | stmn1b | 31.88383 | 2.816717625 | 1.1E-180 | 1.2E-177 |
| **127** | Progenitors 1 | khdrbs1a | 31.86275 | 1.816366792 | 5.6E-163 | 4.6E-160 |
| **128** | Progenitors 1 | rpa2 | 31.80771 | 3.262814283 | 1E-181 | 1.3E-178 |
| **129** | Progenitors 1 | hnrnpaba | 31.73182 | 1.634930372 | 1.1E-165 | 9.9E-163 |
| **130** | Progenitors 1 | ran | 31.64779 | 1.646396875 | 3.9E-163 | 3.3E-160 |
| **131** | Progenitors 1 | anp32b | 31.6227 | 2.78754425 | 1.8E-179 | 1.9E-176 |
| **132** | Progenitors 1 | chaf1a | 31.33116 | 2.990514278 | 6.2E-177 | 6.1E-174 |
| **133** | Progenitors 1 | snrpe | 30.55038 | 2.304834366 | 3.2E-165 | 2.7E-162 |
| **134** | Progenitors 1 | pclaf | 30.1753 | 4.34430027 | 4E-162 | 3.1E-159 |
| **135** | Progenitors 1 | snrpb | 30.16121 | 1.93356967 | 3E-153 | 2.1E-150 |
| **136** | Progenitors 1 | hnrnpa0b | 29.80031 | 1.67334938 | 1.2E-152 | 8E-150 |
| **137** | Progenitors 1 | cirbpb | 29.76588 | 1.492200613 | 5.3E-143 | 3E-140 |
| **138** | Progenitors 1 | hnrnpabb | 29.56882 | 1.672339916 | 1.2E-149 | 7.5E-147 |
| **139** | Progenitors 1 | tuba8l4 | 29.51319 | 1.999989867 | 1.8E-156 | 1.3E-153 |
| **140** | Progenitors 1 | slbp | 29.4879 | 4.300786972 | 3.1E-156 | 2.3E-153 |
| **141** | Progenitors 1 | banf1 | 29.38215 | 3.000735044 | 2.5E-159 | 1.9E-156 |
| **142** | Progenitors 1 | ahcy | 28.97365 | 2.334974051 | 1.2E-152 | 8E-150 |
| **143** | Progenitors 1 | sumo3a | 28.87215 | 2.109611034 | 9.4E-150 | 6.2E-147 |
| **144** | Progenitors 1 | snrpf | 28.60287 | 1.968836784 | 8.7E-145 | 5E-142 |
| **145** | Progenitors 1 | tuba1a | 28.25925 | 2.261639595 | 9.2E-146 | 5.8E-143 |
| **146** | Progenitors 1 | hnrnpa1b | 28.16135 | 2.1130054 | 8.1E-145 | 4.8E-142 |
| **147** | Progenitors 1 | CABZ01005379.1 | 28.06565 | 4.081107616 | 2.7E-145 | 1.6E-142 |
| **148** | Progenitors 1 | rpa3 | 27.86267 | 3.011048555 | 9.5E-146 | 5.9E-143 |
| **149** | Progenitors 1 | ccna2 | 27.85132 | 5.193192959 | 7.5E-140 | 3.8E-137 |
| **150** | Progenitors 1 | calm2b | 27.76758 | 2.044607878 | 8.4E-142 | 4.6E-139 |
| **151** | Progenitors 1 | smc2 | 27.57939 | 4.493327618 | 3.6E-139 | 1.8E-136 |
| **152** | Progenitors 1 | abhd6a | 27.46467 | 3.580729485 | 3.1E-142 | 1.7E-139 |
| **153** | Progenitors 1 | si:ch73-1a9.3 | 27.4144 | 1.864255905 | 1.5E-132 | 7E-130 |
| **154** | Progenitors 1 | lig1 | 27.32397 | 3.173058033 | 5.5E-141 | 2.9E-138 |
| **155** | Progenitors 1 | sumo3b | 27.30627 | 2.569580078 | 1.5E-140 | 7.4E-138 |
| **156** | Progenitors 1 | ddx39ab | 27.13763 | 1.862550378 | 3.2E-132 | 1.4E-129 |
| **157** | Progenitors 1 | her15.1-1 | 27.03404 | 3.495584965 | 6.7E-138 | 3.2E-135 |
| **158** | Progenitors 1 | top2a | 27.01279 | 4.9675107 | 1.2E-132 | 5.9E-130 |
| **159** | Progenitors 1 | ppm1g | 26.82956 | 2.378648281 | 3.1E-135 | 1.5E-132 |
| **160** | Progenitors 1 | ranbp1 | 26.72907 | 1.870456576 | 2.8E-132 | 1.3E-129 |
| **161** | Progenitors 1 | si:ch211-137a8.4 | 26.31446 | 2.234991789 | 5.6E-131 | 2.5E-128 |
| **162** | Progenitors 1 | anp32a | 26.25086 | 1.995781898 | 6.2E-130 | 2.7E-127 |
| **163** | Progenitors 1 | cks1b | 26.14723 | 4.261714935 | 1.9E-128 | 8E-126 |
| **164** | Progenitors 1 | mibp | 26.05821 | 4.348234653 | 3.8E-127 | 1.6E-124 |
| **165** | Progenitors 1 | anp32e | 25.97471 | 1.89624691 | 1.4E-125 | 5.5E-123 |
| **166** | Progenitors 1 | fen1 | 25.95778 | 3.096474171 | 4.8E-129 | 2E-126 |
| **167** | Progenitors 1 | zgc:110540 | 25.95219 | 4.499548912 | 8.6E-126 | 3.4E-123 |
| **168** | Progenitors 1 | h2afx | 25.71205 | 2.896784782 | 6.2E-127 | 2.5E-124 |
| **169** | Progenitors 1 | id1 | 25.69344 | 3.157251835 | 6.8E-127 | 2.7E-124 |
| **170** | Progenitors 1 | baz1b | 25.62452 | 2.442528725 | 5.4E-126 | 2.1E-123 |
| **171** | Progenitors 1 | cbx5 | 25.46941 | 2.536276817 | 8.2E-125 | 3.1E-122 |
| **172** | Progenitors 1 | tuba1c | 25.10795 | 2.305203199 | 2.9E-121 | 1E-118 |
| **173** | Progenitors 1 | CABZ01058261.1 | 25.01974 | 4.617056847 | 1.1E-117 | 3.7E-115 |
| **174** | Progenitors 1 | her4.1 | 24.99796 | 3.3596313 | 5.2E-121 | 1.8E-118 |
| **175** | Progenitors 1 | hdac1 | 24.89681 | 1.754298449 | 3.2E-116 | 1.1E-113 |
| **176** | Progenitors 1 | setb | 24.74794 | 1.799617887 | 2.1E-116 | 7.1E-114 |
| **177** | Progenitors 1 | nutf2l | 24.6904 | 3.120162249 | 2.3E-118 | 8.1E-116 |
| **178** | Progenitors 1 | smc1al | 24.66069 | 1.887411833 | 7.2E-116 | 2.4E-113 |
| **179** | Progenitors 1 | magoh | 24.58812 | 1.993138671 | 7.8E-116 | 2.5E-113 |
| **180** | Progenitors 1 | cdk1 | 24.54513 | 4.643316269 | 1.1E-113 | 3.5E-111 |
| **181** | Progenitors 1 | hnrnpa0l-1 | 24.51803 | 1.10616529 | 1.3E-110 | 3.9E-108 |
| **182** | Progenitors 1 | ncapg | 24.48101 | 4.886284828 | 1E-113 | 3.3E-111 |
| **183** | Progenitors 1 | smc4 | 24.46729 | 3.825938225 | 4.3E-115 | 1.4E-112 |
| **184** | Progenitors 1 | syncrip | 24.43125 | 1.531150579 | 8.3E-111 | 2.4E-108 |
| **185** | Progenitors 1 | snrpd2 | 24.40524 | 1.759106874 | 3.9E-113 | 1.2E-110 |
| **186** | Progenitors 1 | taf15 | 24.2833 | 1.957806468 | 5.4E-113 | 1.7E-110 |
| **187** | Progenitors 1 | dhfr | 24.23506 | 3.964292526 | 4.5E-113 | 1.4E-110 |
| **188** | Progenitors 1 | cenpf | 24.17227 | 4.719338894 | 8.5E-111 | 2.5E-108 |
| **189** | Progenitors 1 | actl6a | 24.06294 | 2.283429861 | 1.4E-112 | 4.2E-110 |
| **190** | Progenitors 1 | dut | 23.99543 | 3.155126333 | 2.3E-112 | 6.8E-110 |
| **191** | Progenitors 1 | h2afva | 23.85158 | 1.949113607 | 2.6E-110 | 7.3E-108 |
| **192** | Progenitors 1 | mad2l1 | 23.68154 | 4.58840847 | 1.3E-107 | 3.6E-105 |
| **193** | Progenitors 1 | nasp | 23.63137 | 3.372167587 | 2.7E-109 | 7.6E-107 |
| **194** | Progenitors 1 | marcksb | 23.56229 | 1.464284182 | 2.8E-104 | 7.3E-102 |
| **195** | Progenitors 1 | tpx2 | 23.27903 | 5.679986 | 5.3E-103 | 1.4E-100 |
| **196** | Progenitors 1 | srsf2a | 23.27819 | 1.755615711 | 5.9E-105 | 1.6E-102 |
| **197** | Progenitors 1 | lmnb2 | 23.11164 | 2.727295876 | 1.6E-105 | 4.2E-103 |
| **198** | Progenitors 1 | aurkb | 23.02791 | 4.603858471 | 1.4E-102 | 3.5E-100 |
| **199** | Progenitors 1 | si:ch73-21g5.7 | 23.00684 | 3.866025209 | 2.2E-103 | 5.7E-101 |
