## Supplementary_file_6 for "Single cell RNA sequencing unravels the transcriptional network underlying zebrafish retina regeneration"

**Differentially expressed genes between Müller glia_Progenitors 1 and Progenitors 2 clusters**

| | group | names | scores | logfoldchanges | pvals | pvals_adj |
|---|---|---|---|---|---|---|
| 0 | Progenitors 1 | id1 | 33.20395 | 4.413954258 | 5.8E-190 | 1.6E-185 |
| 1 | Progenitors 1 | si:dkey-238o13.4 | 30.18699 | 2.934452057 | 2.4E-168 | 3.4E-164 |
| 2 | Progenitors 1 | f3a | 29.49284 | 2.489398479 | 1.3E-161 | 1.2E-157 |
| 3 | Progenitors 1 | crlf1a | 29.26157 | 3.758566618 | 1.7E-155 | 9.5E-152 |
| 4 | Progenitors 1 | pim1 | 28.90828 | 2.224119425 | 9.2E-156 | 6.3E-152 |
| 5 | Progenitors 1 | si:ch1073-303k11.2 | 27.48717 | 4.214794159 | 1.3E-138 | 5.2E-135 |
| 6 | Progenitors 1 | acbd7 | 26.48776 | 2.141279697 | 9.1E-135 | 3.1E-131 |
| 7 | Progenitors 1 | her4.1 | 26.04419 | 3.313039064 | 6.2E-130 | 1.9E-126 |
| 8 | Progenitors 1 | nrarpa | 25.8404 | 2.959989309 | 3.4E-128 | 9.5E-125 |
| 9 | Progenitors 1 | her12 | 25.70443 | 4.233252048 | 4.1E-124 | 1E-120 |
| 10 | Progenitors 1 | akap12b | 24.70734 | 2.195334911 | 2.6E-119 | 6E-116 |
| 11 | Progenitors 1 | notch3 | 23.9893 | 2.590406418 | 2.6E-113 | 5.6E-110 |
| 12 | Progenitors 1 | six3b | 23.81036 | 1.77921629 | 6.9E-110 | 1.4E-106 |
| 13 | Progenitors 1 | ppdpfb | 23.02896 | 1.329558372 | 8.8E-102 | 1.43E-98 |
| 14 | Progenitors 1 | myl6 | 22.7005 | 2.129921675 | 7.1E-103 | 1.2E-99 |
| 15 | Progenitors 1 | ckbb | 22.66209 | 1.653145909 | 8.7E-101 | 1.33E-97 |
| 16 | Progenitors 1 | her4.2 | 22.06587 | 2.83970499 | 1.86E-97 | 2.71E-94 |
| 17 | Progenitors 1 | s1pr1 | 21.78406 | 2.389015436 | 1.44E-95 | 1.99E-92 |
| 18 | Progenitors 1 | crabp1a | 21.38036 | 1.167667389 | 2.7E-90 | 2.49E-87 |
| 19 | Progenitors 1 | abhd6a | 21.29357 | 2.585156679 | 9.24E-92 | 1.02E-88 |
| 20 | Progenitors 1 | mdka | 21.22327 | 1.112773299 | 2.56E-89 | 2.29E-86 |
| 21 | Progenitors 1 | clcf1 | 21.19512 | 2.955190897 | 2.18E-90 | 2.08E-87 |
| 22 | Progenitors 1 | lfng | 21.14992 | 2.12735033 | 1.07E-90 | 1.06E-87 |
| 23 | Progenitors 1 | zgc:165604 | 20.9498 | 2.277417898 | 3.97E-89 | 3.44E-86 |
| 24 | Progenitors 1 | sgk1 | 20.57258 | 2.153960943 | 2.68E-86 | 2.25E-83 |
| 25 | Progenitors 1 | cnn2 | 20.57133 | 2.752839804 | 8.36E-86 | 6.61E-83 |
| 26 | Progenitors 1 | flna | 20.52126 | 2.272691727 | 7.19E-86 | 5.86E-83 |
| 27 | Progenitors 1 | tjp2b | 19.55174 | 2.835801125 | 3.3E-78 | 2.4E-75 |
| 28 | Progenitors 1 | CU467822.1 | 19.54188 | 1.552370071 | 3.78E-78 | 2.68E-75 |
| 29 | Progenitors 1 | rtn4a | 19.51699 | 1.212570786 | 1.01E-77 | 6.97E-75 |
| 30 | Progenitors 1 | zgc:158343 | 19.33073 | 2.217153788 | 5.58E-77 | 3.68E-74 |
| 31 | Progenitors 1 | mmp9 | 19.04083 | 3.244989395 | 2.58E-74 | 1.63E-71 |
| 32 | Progenitors 1 | marcksl1b | 18.82026 | 1.07450664 | 5.22E-72 | 3.08E-69 |
| 33 | Progenitors 1 | nr2e1 | 18.09925 | 2.058402538 | 2.03E-68 | 1.15E-65 |
| 34 | Progenitors 1 | yap1 | 18.01408 | 1.959831595 | 7.13E-68 | 3.95E-65 |
| 35 | Progenitors 1 | gfap | 17.93883 | 1.542437434 | 4.78E-67 | 2.54E-64 |
| 36 | Progenitors 1 | si:ch211-251b21.1 | 17.4214 | 1.659507871 | 7.45E-64 | 3.75E-61 |
| 37 | Progenitors 1 | tpm4a | 17.3204 | 3.760930061 | 2.43E-62 | 1.18E-59 |
| 38 | Progenitors 1 | her9 | 17.23831 | 1.610290527 | 2.09E-62 | 1.03E-59 |
| 39 | Progenitors 1 | sox9b | 17.1651 | 1.834976196 | 4.36E-62 | 2.08E-59 |
| 40 | Progenitors 1 | f3b | 16.94719 | 2.167218685 | 1.26E-60 | 5.82E-58 |
| 41 | Progenitors 1 | btg2 | 16.88596 | 1.013802886 | 1.18E-59 | 5.28E-57 |
| 42 | Progenitors 1 | errfi1a | 16.83897 | 3.918623209 | 4.94E-59 | 2.09E-56 |
| 43 | Progenitors 1 | zgc:109949 | 16.77024 | 1.86362648 | 1.46E-59 | 6.42E-57 |
| 44 | Progenitors 1 | midn | 16.36706 | 1.434379101 | 6.78E-57 | 2.68E-54 |
| 45 | Progenitors 1 | si:ch73-21g5.7 | 16.01612 | 2.184960127 | 9.71E-55 | 3.68E-52 |
| 46 | Progenitors 1 | nrarpb | 15.56761 | 2.937930107 | 1.14E-51 | 4.16E-49 |
| 47 | Progenitors 1 | cspg5a | 15.4924 | 2.866997004 | 2.78E-51 | 9.88E-49 |
| 48 | Progenitors 1 | her4.2-1 | 15.35021 | 2.399408817 | 1.69E-50 | 5.79E-48 |
| 49 | Progenitors 1 | si:dkey-7j14.6 | 15.33142 | 1.080294251 | 3.95E-50 | 1.32E-47 |
| 50 | Progenitors 1 | sox2 | 15.09371 | 1.383063436 | 6.7E-49 | 2.16E-46 |
| 51 | Progenitors 1 | slc12a4 | 15.05737 | 3.736255169 | 2.61E-48 | 8.12E-46 |
| 52 | Progenitors 1 | lgals2a | 14.99862 | 1.881000519 | 1.54E-48 | 4.85E-46 |
| 53 | Progenitors 1 | sparc | 14.86773 | 2.276907206 | 1.07E-47 | 3.22E-45 |
| 54 | Progenitors 1 | BX465834.1 | 14.79454 | 1.47226131 | 2.85E-47 | 8.48E-45 |
| 55 | Progenitors 1 | igfbp5b | 14.72426 | 2.648566484 | 1.16E-46 | 3.35E-44 |
| 56 | Progenitors 1 | txn | 14.70218 | 1.462990165 | 8.83E-47 | 2.57E-44 |
| 57 | Progenitors 1 | myl9b | 14.54179 | 1.423460126 | 8.91E-46 | 2.49E-43 |
| 58 | Progenitors 1 | qkia | 14.47649 | 1.297585011 | 2.38E-45 | 6.58E-43 |
| 59 | Progenitors 1 | bzw1b | 14.4581 | 0.96661377 | 4.71E-45 | 1.29E-42 |
| 60 | Progenitors 1 | dab2ipb | 14.29183 | 2.518965721 | 2.49E-44 | 6.62E-42 |
| 61 | Progenitors 1 | her15.1-1 | 14.10235 | 1.568661213 | 2.82E-43 | 7.16E-41 |
| 62 | Progenitors 1 | fxyd6l | 14.07612 | 1.640132904 | 3.47E-43 | 8.73E-41 |
| 63 | Progenitors 1 | notch1b | 13.86108 | 2.192939758 | 5.87E-42 | 1.44E-39 |
| 64 | Progenitors 1 | cdh2 | 13.75786 | 0.83481735 | 4.1E-41 | 9.8E-39 |
| 65 | Progenitors 1 | cnn3a | 13.69488 | 1.521155357 | 4.73E-41 | 1.12E-38 |
| 66 | Progenitors 1 | her4.4 | 13.66234 | 2.617347002 | 8.68E-41 | 2.02E-38 |
| 67 | Progenitors 1 | angptl4 | 13.35335 | 1.231126189 | 4.22E-39 | 9.41E-37 |
| 68 | Progenitors 1 | rasgef1bb | 13.24782 | 1.089429498 | 1.64E-38 | 3.59E-36 |
| 69 | Progenitors 1 | mCherry | 13.21127 | 1.574850559 | 2.05E-38 | 4.48E-36 |
| 70 | Progenitors 1 | tmsb4x | 13.05165 | 0.654317379 | 3.96E-37 | 8.5E-35 |
| 71 | Progenitors 1 | LO018196.1 | 12.83982 | 2.251034737 | 1.95E-36 | 4.12E-34 |
| 72 | Progenitors 1 | dusp5 | 12.81894 | 1.963020921 | 2.4E-36 | 5.03E-34 |
| 73 | Progenitors 1 | her15.1 | 12.81567 | 1.637444735 | 2.44E-36 | 5.08E-34 |
| 74 | Progenitors 1 | fosab | 12.7846 | 1.466152191 | 3.55E-36 | 7.33E-34 |
| 75 | Progenitors 1 | nocta | 12.73726 | 1.494959116 | 6.29E-36 | 1.28E-33 |
| 76 | Progenitors 1 | jdp2b | 12.71978 | 1.816286087 | 8.08E-36 | 1.62E-33 |
| 77 | Progenitors 1 | gpm6aa | 12.50174 | 0.74019593 | 1.88E-34 | 3.53E-32 |
| 78 | Progenitors 1 | col18a1a | 12.48219 | 1.59650743 | 1.28E-34 | 2.45E-32 |
| 79 | Progenitors 1 | fhl3b | 12.38253 | 2.873163939 | 5.13E-34 | 9.42E-32 |
| 80 | Progenitors 1 | fosl1a | 12.37457 | 1.607260108 | 4.5E-34 | 8.37E-32 |
| 81 | Progenitors 1 | emilin1a | 12.23998 | 4.57828331 | 5.43E-33 | 9.77E-31 |
| 82 | Progenitors 1 | rbpms2b | 12.17257 | 1.659262776 | 4.67E-33 | 8.45E-31 |
| 83 | Progenitors 1 | BX284638.1 | 12.12708 | 1.292871237 | 8.64E-33 | 1.5E-30 |
| 84 | Progenitors 1 | slmapb | 12.12079 | 1.79136312 | 8.49E-33 | 1.49E-30 |
| 85 | Progenitors 1 | atp1b1a | 12.07991 | 1.148278236 | 1.48E-32 | 2.54E-30 |
| 86 | Progenitors 1 | zic2b | 11.99187 | 1.104104042 | 4.22E-32 | 7.16E-30 |
| 87 | Progenitors 1 | si:ch73-335l21.4 | 11.95505 | 1.246983647 | 5.98E-32 | 1.01E-29 |
| 88 | Progenitors 1 | nrp2b | 11.8913 | 1.182368517 | 1.33E-31 | 2.2E-29 |
| 89 | Progenitors 1 | eml1 | 11.88188 | 1.894070268 | 1.28E-31 | 2.13E-29 |
| 90 | Progenitors 1 | sb:cb81 | 11.86705 | 3.127805233 | 2.25E-31 | 3.67E-29 |
| 91 | Progenitors 1 | rhbg | 11.85136 | 2.234927654 | 1.89E-31 | 3.12E-29 |
| 92 | Progenitors 1 | ccdc80 | 11.84705 | 5.136306763 | 4.75E-31 | 7.56E-29 |
| 93 | Progenitors 1 | cxcl14 | 11.83785 | 1.708183408 | 2.09E-31 | 3.43E-29 |
| 94 | Progenitors 1 | crip3 | 11.83164 | 3.947635412 | 4.14E-31 | 6.67E-29 |
| 95 | Progenitors 1 | CR848047.1 | 11.81629 | 1.902240515 | 2.67E-31 | 4.33E-29 |
| 96 | Progenitors 1 | efhd1 | 11.80226 | 3.130785942 | 4.37E-31 | 7E-29 |
| 97 | Progenitors 1 | bcar1 | 11.7913 | 3.947091818 | 6.35E-31 | 9.99E-29 |
| 98 | Progenitors 1 | cited4a | 11.71951 | 0.827466905 | 1.21E-30 | 1.87E-28 |
| 99 | Progenitors 1 | eif4ebp3l | 11.7086 | 1.782539248 | 8.85E-31 | 1.38E-28 |

| | group | names | scores | logfoldchanges | pvals | pvals_adj |
|---|---|---|---|---|---|---|
| 100 | Progenitors 2 | insm1a | 29.74285 | 4.062833309 | 8.5E-139 | 3.9E-135 |
| 101 | Progenitors 2 | pou2f2a-1 | 25.10969 | 3.67982173 | 1.1E-107 | 2.1E-104 |
| 102 | Progenitors 2 | rbpjb | 22.31904 | 2.4372015 | 3.5E-92 | 4.04E-89 |
| 103 | Progenitors 2 | si:dkey-56m19.5 | 22.09177 | 1.84433341 | 3.4E-93 | 4.09E-90 |
| 104 | Progenitors 2 | h3f3b.1 | 21.74553 | 1.010189891 | 5.04E-95 | 6.65E-92 |
| 105 | Progenitors 2 | tubb2b | 21.56813 | 1.212004542 | 6.26E-94 | 7.88E-91 |
| 106 | Progenitors 2 | stmn1a | 21.35702 | 1.334182978 | 3.58E-91 | 3.68E-88 |
| 107 | Progenitors 2 | hmgb2a | 21.26676 | 0.923674226 | 1.41E-91 | 1.5E-88 |
| 108 | Progenitors 2 | im:7152348 | 20.84255 | 2.880055189 | 3.99E-81 | 3.07E-78 |
| 109 | Progenitors 2 | tent5ba | 20.08026 | 3.247291088 | 2.5E-75 | 1.61E-72 |
| 110 | Progenitors 2 | mex3b | 20.07191 | 2.233929873 | 2.8E-77 | 1.89E-74 |
| 111 | Progenitors 2 | neurod4 | 19.89341 | 4.068521023 | 1.63E-73 | 1E-70 |
| 112 | Progenitors 2 | stmn1b | 19.65258 | 1.195197701 | 2.2E-78 | 1.65E-75 |
| 113 | Progenitors 2 | sox11a | 19.12948 | 1.846146345 | 2.73E-72 | 1.64E-69 |
| 114 | Progenitors 2 | hes6 | 18.84141 | 1.767419577 | 2.31E-70 | 1.33E-67 |
| 115 | Progenitors 2 | nusap1 | 18.2124 | 1.839676142 | 4.71E-67 | 2.54E-64 |
| 116 | Progenitors 2 | golga7ba | 18.13729 | 2.684294224 | 3.23E-64 | 1.66E-61 |
| 117 | Progenitors 2 | mycla | 17.77845 | 2.757133484 | 6.22E-62 | 2.92E-59 |
| 118 | Progenitors 2 | ndrg1b | 17.5824 | 4.818793297 | 1.87E-59 | 8.09E-57 |
| 119 | Progenitors 2 | mki67 | 17.57771 | 1.183243871 | 1.11E-64 | 5.79E-62 |
| 120 | Progenitors 2 | foxn4 | 17.39317 | 2.271308422 | 2.42E-60 | 1.1E-57 |
| 121 | Progenitors 2 | cdkn1ca | 17.2424 | 3.572548866 | 5.75E-58 | 2.38E-55 |
| 122 | Progenitors 2 | itm2cb | 17.06698 | 1.813652277 | 4.99E-59 | 2.09E-56 |
| 123 | Progenitors 2 | nsg2 | 17.01987 | 2.492872 | 8.78E-58 | 3.57E-55 |
| 124 | Progenitors 2 | chd7 | 16.56265 | 1.074338913 | 3.52E-57 | 1.41E-54 |
| 125 | Progenitors 2 | cenpf | 16.49356 | 1.50218153 | 1.3E-56 | 5.07E-54 |
| 126 | Progenitors 2 | sox11b | 16.28317 | 1.535938144 | 8.72E-55 | 3.35E-52 |
| 127 | Progenitors 2 | hmga1a | 15.71892 | 0.713748991 | 7.69E-53 | 2.88E-50 |
| 128 | Progenitors 2 | top2a | 15.68546 | 1.314869165 | 6.31E-52 | 2.33E-49 |
| 129 | Progenitors 2 | atoh7 | 15.63489 | 3.633435249 | 2.9E-49 | 9.56E-47 |
| 130 | Progenitors 2 | si:ch211-137a8.4 | 15.55755 | 0.957719445 | 2.97E-51 | 1.04E-48 |
| 131 | Progenitors 2 | zswim5 | 15.55344 | 1.32071054 | 1.99E-50 | 6.72E-48 |
| 132 | Progenitors 2 | tuba8l4 | 15.51112 | 0.796337485 | 1.75E-51 | 6.28E-49 |
| 133 | Progenitors 2 | ccna2 | 15.48092 | 1.218245983 | 1.13E-50 | 3.93E-48 |
| 134 | Progenitors 2 | ube2c | 15.25562 | 1.485815406 | 5.35E-49 | 1.74E-46 |
| 135 | Progenitors 2 | si:ch211-255i3.4 | 15.19922 | 1.93849504 | 8.56E-48 | 2.6E-45 |
| 136 | Progenitors 2 | cdk1 | 15.18764 | 1.355599523 | 7.58E-49 | 2.41E-46 |
| 137 | Progenitors 2 | histh1l | 15.01661 | 1.247156382 | 7.09E-48 | 2.18E-45 |
| 138 | Progenitors 2 | si:ch211-222l21.1 | 14.71231 | 0.667203367 | 7.84E-47 | 2.31E-44 |
| 139 | Progenitors 2 | h3f3b.1-2 | 14.68723 | 0.852724254 | 3.42E-46 | 9.66E-44 |
| 140 | Progenitors 2 | si:ch73-281n10.2 | 14.63193 | 0.750804663 | 2.76E-46 | 7.87E-44 |
| 141 | Progenitors 2 | dhx32b | 14.50244 | 1.76413238 | 3.27E-44 | 8.61E-42 |
| 142 | Progenitors 2 | CABZ01058261.1 | 14.49623 | 1.203343987 | 6.88E-45 | 1.85E-42 |
| 143 | Progenitors 2 | otx2b | 14.43487 | 3.108237505 | 6.7E-43 | 1.67E-40 |
| 144 | Progenitors 2 | tubb4b | 14.4129 | 0.70184356 | 6.07E-45 | 1.65E-42 |
| 145 | Progenitors 2 | onecut2 | 14.3852 | 3.557531118 | 1.42E-42 | 3.52E-40 |
| 146 | Progenitors 2 | znrf1 | 14.34256 | 1.800945878 | 2.61E-43 | 6.7E-41 |
| 147 | Progenitors 2 | tuba8l | 14.25486 | 1.170437932 | 1.19E-43 | 3.12E-41 |
| 148 | Progenitors 2 | tuba1a | 14.23932 | 0.852048218 | 1.27E-43 | 3.29E-41 |
| 149 | Progenitors 2 | tfap2d | 14.04352 | 5.688826084 | 1.92E-40 | 4.43E-38 |
| 150 | Progenitors 2 | pou2f2a | 13.93332 | 3.183808088 | 2.64E-40 | 6.03E-38 |
| 151 | Progenitors 2 | aurkb | 13.92913 | 1.201517344 | 8.89E-42 | 2.16E-39 |
| 152 | Progenitors 2 | alcamb | 13.89505 | 1.827475429 | 6.68E-41 | 1.57E-38 |
| 153 | Progenitors 2 | lbr | 13.81296 | 0.898884833 | 1.89E-41 | 4.54E-39 |
| 154 | Progenitors 2 | birc5a | 13.66098 | 1.532848597 | 6.51E-40 | 1.48E-37 |
| 155 | Progenitors 2 | cnp | 13.48519 | 1.066300035 | 3.23E-39 | 7.28E-37 |
| 156 | Progenitors 2 | dek | 13.32753 | 0.782036602 | 7.34E-39 | 1.63E-36 |
| 157 | Progenitors 2 | h1f0 | 13.20192 | 1.04755187 | 5.63E-38 | 1.22E-35 |
| 158 | Progenitors 2 | dclk1a | 13.09317 | 1.696277022 | 8.81E-37 | 1.88E-34 |
| 159 | Progenitors 2 | pif1 | 13.01037 | 2.443202496 | 4.76E-36 | 9.77E-34 |
| 160 | Progenitors 2 | bhlhe22 | 12.95476 | 3.421462774 | 2.1E-35 | 4.12E-33 |
| 161 | Progenitors 2 | tpx2 | 12.83766 | 1.256715894 | 7.3E-36 | 1.48E-33 |
| 162 | Progenitors 2 | mad2l1 | 12.75631 | 1.149626374 | 1.5E-35 | 2.96E-33 |
| 163 | Progenitors 2 | cited4b | 12.75119 | 2.145797729 | 6.66E-35 | 1.3E-32 |
| 164 | Progenitors 2 | tuba1c | 12.72924 | 0.816478729 | 1.26E-35 | 2.51E-33 |
| 165 | Progenitors 2 | cdc20 | 12.69682 | 1.614559889 | 7.25E-35 | 1.4E-32 |
| 166 | Progenitors 2 | neurod1 | 12.6789 | 2.726300716 | 3.05E-34 | 5.71E-32 |
| 167 | Progenitors 2 | tp53inp2 | 12.57528 | 1.101814985 | 1.53E-34 | 2.91E-32 |
| 168 | Progenitors 2 | lepr | 12.56878 | 1.283785105 | 1.38E-34 | 2.64E-32 |
| 169 | Progenitors 2 | hmgn2 | 12.3784 | 0.512889981 | 4.68E-34 | 8.63E-32 |
| 170 | Progenitors 2 | tsc22d1 | 12.32308 | 0.950090408 | 2.3E-33 | 4.18E-31 |
| 171 | Progenitors 2 | cep55l | 12.26544 | 1.510048032 | 7.61E-33 | 1.34E-30 |
| 172 | Progenitors 2 | kifc1 | 12.26017 | 1.367429972 | 6.83E-33 | 1.22E-30 |
| 173 | Progenitors 2 | srrm4 | 12.2442 | 1.469046593 | 8.77E-33 | 1.52E-30 |
| 174 | Progenitors 2 | ncapg | 12.21664 | 1.066959977 | 7.59E-33 | 1.34E-30 |
| 175 | Progenitors 2 | rimkla | 12.10791 | 1.129550576 | 2.92E-32 | 4.99E-30 |
| 176 | Progenitors 2 | kif22 | 12.04421 | 1.292233944 | 7.47E-32 | 1.25E-29 |
| 177 | Progenitors 2 | barhl2 | 11.87954 | 3.747173786 | 2.33E-30 | 3.54E-28 |
| 178 | Progenitors 2 | tfap2c | 11.84417 | 2.713200808 | 2.11E-30 | 3.24E-28 |
| 179 | Progenitors 2 | dlgap5 | 11.83674 | 1.166193247 | 5.63E-31 | 8.91E-29 |
| 180 | Progenitors 2 | robo3 | 11.82311 | 2.347760439 | 2.12E-30 | 3.24E-28 |
| 181 | Progenitors 2 | cxxc5a | 11.78868 | 1.031479836 | 1.03E-30 | 1.6E-28 |
| 182 | Progenitors 2 | prdm1b | 11.64075 | 3.057979107 | 2.1E-29 | 3.02E-27 |
| 183 | Progenitors 2 | scml2 | 11.63646 | 2.177360296 | 1.38E-29 | 2E-27 |
| 184 | Progenitors 2 | calm2b | 11.62686 | 0.624697924 | 2.67E-30 | 4.04E-28 |
| 185 | Progenitors 2 | si:dkey-28b4.8 | 11.61454 | 1.312385201 | 8.66E-30 | 1.28E-27 |
| 186 | Progenitors 2 | si:ch211-69g19.2 | 11.59842 | 1.706728339 | 1.28E-29 | 1.87E-27 |
| 187 | Progenitors 2 | arpp19b | 11.58961 | 0.917288542 | 6.81E-30 | 1.01E-27 |
| 188 | Progenitors 2 | anp32a | 11.58766 | 0.658562183 | 4.87E-30 | 7.32E-28 |
| 189 | Progenitors 2 | plcxd3 | 11.55526 | 2.793503523 | 4.62E-29 | 6.53E-27 |
| 190 | Progenitors 2 | hmgb3a | 11.35004 | 0.783460557 | 8.71E-29 | 1.21E-26 |
| 191 | Progenitors 2 | myt1a | 11.34812 | 1.462108612 | 1.46E-28 | 2.01E-26 |
| 192 | Progenitors 2 | pik3r3b | 11.2848 | 1.71014452 | 4.09E-28 | 5.53E-26 |
| 193 | Progenitors 2 | hes2.2 | 11.24021 | 1.975386739 | 5.44E-28 | 7.31E-26 |
| 194 | Progenitors 2 | kif11 | 11.21864 | 1.493881106 | 6.47E-28 | 8.66E-26 |
| 195 | Progenitors 2 | lrrfip1a | 11.02948 | 2.591686726 | 8.21E-27 | 1.05E-24 |
| 196 | Progenitors 2 | h2afva | 11.0211 | 0.676322818 | 2.32E-27 | 3.06E-25 |
| 197 | Progenitors 2 | plk1 | 10.98766 | 1.178636551 | 4.87E-27 | 6.36E-25 |
| 198 | Progenitors 2 | si:ch211-207i1.2 | 10.97547 | 1.301731825 | 5.85E-27 | 7.57E-25 |
| 199 | Progenitors 2 | kmt5ab | 10.92091 | 1.760930061 | 1.58E-26 | 2.01E-24 |
